# Substrate and target selectivity of 4′-fluoroadenosine against viral and host polymerases

**DOI:** 10.64898/2026.05.22.727251

**Authors:** Simon M. Walker, Arlo J. Loutan, Egor P. Tchesnokov, Dana Kocincova, Calvin J. Gordon, Ruby A. Escobedo, Nathaniel Jackson, Olivia A. Vogel, Kim Morsheimer, Suncheol Park, Anant Gharpure, Ixchel Urbano, Mina Heacock, Zhong Cheng, Kushboo Pathak, Karen C. Wolff, Lauren Huerta, Malina A. Bakowski, Laura Riva, Anil K. Gupta, Chenguang Yu, Kalyan Das, Luis Martinez-Sobrido, Christopher F. Basler, Robert Davey, Ian A. Wilson, Andrew B. Ward, Sumit Chanda, Arnab K. Chatterjee, Matthias Götte

## Abstract

Developing safe and effective treatments against emerging RNA viruses is an important goal in pandemic preparedness efforts. 4′-fluorouridine (4′-FlU) is a broad-spectrum antiviral that was shown to inhibit viral RNA-dependent RNA polymerases (RdRps). Given its notable range of antiviral activity, this class of nucleoside analogs warrants further investigation. Here, we studied the antiviral activity and underlying mechanism of inhibition of 4′-fluoroadenosine (4′-FlA). Like 4′-FlU, 4′-FlA demonstrates a broad-spectrum of antiviral activity against eight prototypic viruses representing diverse families. Enzyme kinetics show that the triphosphate (4′-FlA-TP) is efficiently incorporated by viral RdRps. A cryo-EM structure of RdRp of severe acute respiratory syndrome coronavirus 2 (SARS-CoV-2) in complex with double-stranded RNA and the incorporated monophosphate (4′-FlA-MP) characterizes interactions at the active site. The incorporated analog elicits heterogeneous inhibition patterns in primer extension reactions. In contrast, templates with embedded 4′-FlA-MP inhibit incorporation of complementary UTP across the viral RdRps. However, incorporation of 4′-FIA-TP is not limited to viral polymerases and likewise includes human mitochondrial RNA polymerase. These results demonstrate the general potential for 4′-fluorinated nucleotides as antiviral drugs and guide the development of more selective derivatives for medical use in appropriate settings.

## Introduction

Diverse positive- and negative-sense RNA viruses are associated with human diseases and have the potential to cause outbreaks, epidemics, or pandemics (1,2). Severe acute respiratory syndrome coronavirus 2 (SARS-CoV-2) and the coronavirus disease 2019 (COVID-19) pandemic provide a recent example and highlight the need for effective treatments during outbreaks. The viral RNA-dependent RNA polymerase (RdRp) is essential for viral replication and provides a logical target for the development of antiviral drugs (3). While structural features of these enzymes vary across viral families, the active site is generally conserved to accommodate nucleoside triphosphates (NTPs) (4–6). This feature provides opportunities to develop broad-spectrum antivirals that could be rapidly employed against emerging pathogens.

Nucleoside and nucleotide analogs (NAs) mimic natural NTPs and commonly inhibit viral RNA synthesis once incorporated into the growing RNA strand (4,7). Remdesivir (RDV), a 1′-cyano modified *C*-linked adenosine monophosphate prodrug, interferes with RNA synthesis by SARS-CoV-2 RdRp and has been approved for the treatment of COVID-19 (8,9). RDV shows antiviral activity against a broad spectrum of virus families, including the *Corona-*, *Picorna-*, *Flavi-*, *Filo-*, *Paramyxo-*, and *Pneumoviridae* (10–14). RDV is less active or not active against several segmented negative-sense viruses, such as members of the *Nairoviridae* (Crimean-Congo hemorrhagic fever or CCHFV), *Arenaviridae*, and *Orthomyxoviridae* (influenza viruses)(10,11). Molnupiravir, a cytidine analog and prodrug of β-D-N^4^-hydroxycytidine (NHC), has also demonstrated a broad range of activity, including several segmented negative-sense RNA viruses (15–24). The investigational 4′-fluorouridine (4′-FlU) shows a similar profile (25–33). While molnupiravir acts as a mutagenic nucleotide, RDV and 4′-FlU inhibit viral RNA synthesis (34).

Data that identify the biochemical attributes underlying the broad-spectrum activity of 4′-FlU remain elusive. Previous work has demonstrated that the active triphosphate form, 4′-FlU-TP, is used as a substrate by the viral RdRp of respiratory syncytial virus (RSV), SARS-CoV-2, and influenza A (FluA) (29,30). For both RSV and FluA RdRp, the natural counterpart UTP outcompetes 4′-FlU-TP approximately 4-fold. Once incorporated, 4′-FlU displays heterogeneous effects on primer-strand extension. Against RSV RdRp, inhibition at the site of incorporation “i” and further downstream point to contributions from immediate and delayed chain-termination, respectively (30). For SARS-CoV-2 RdRp, a single incorporation of 4′-FlU still generates the full-length product (30). In the case of FluA RdRp, there is evidence for immediate chain-termination (29). Overall, the mechanism of action of 4′-FlU appears heterogeneous and dependent on the nature of the viral RdRp.

Collectively, these studies on 4′-FlU raise several important questions. Does the 4′-fluoro modification also mediate antiviral effects in a different base context? How specific is the inhibition of viral polymerases? Is there a common underlying mechanism of inhibition against diverse polymerase targets? To address these questions, we synthesized a purine derivative of 4′-FlU, 4′-fluoroadenosine (4′-FlA) (Figure 1A), and show that the spectrum of antiviral activity is comparable to that of 4′-FlU. The active form, 4′-FlA-TP, is efficiently incorporated by an array of corresponding viral RNA polymerases. We determined a cryo-EM structure of SARS-CoV-2 RdRp with the nucleotide analog to identify its binding pose at the active site. Comparison with other viral polymerase structures demonstrates broad-spectrum antiviral activity. Several host polymerases also use 4′-FlA-TP as a substrate, which needs to be addressed in promising drug development programs.

**Figure 1.**
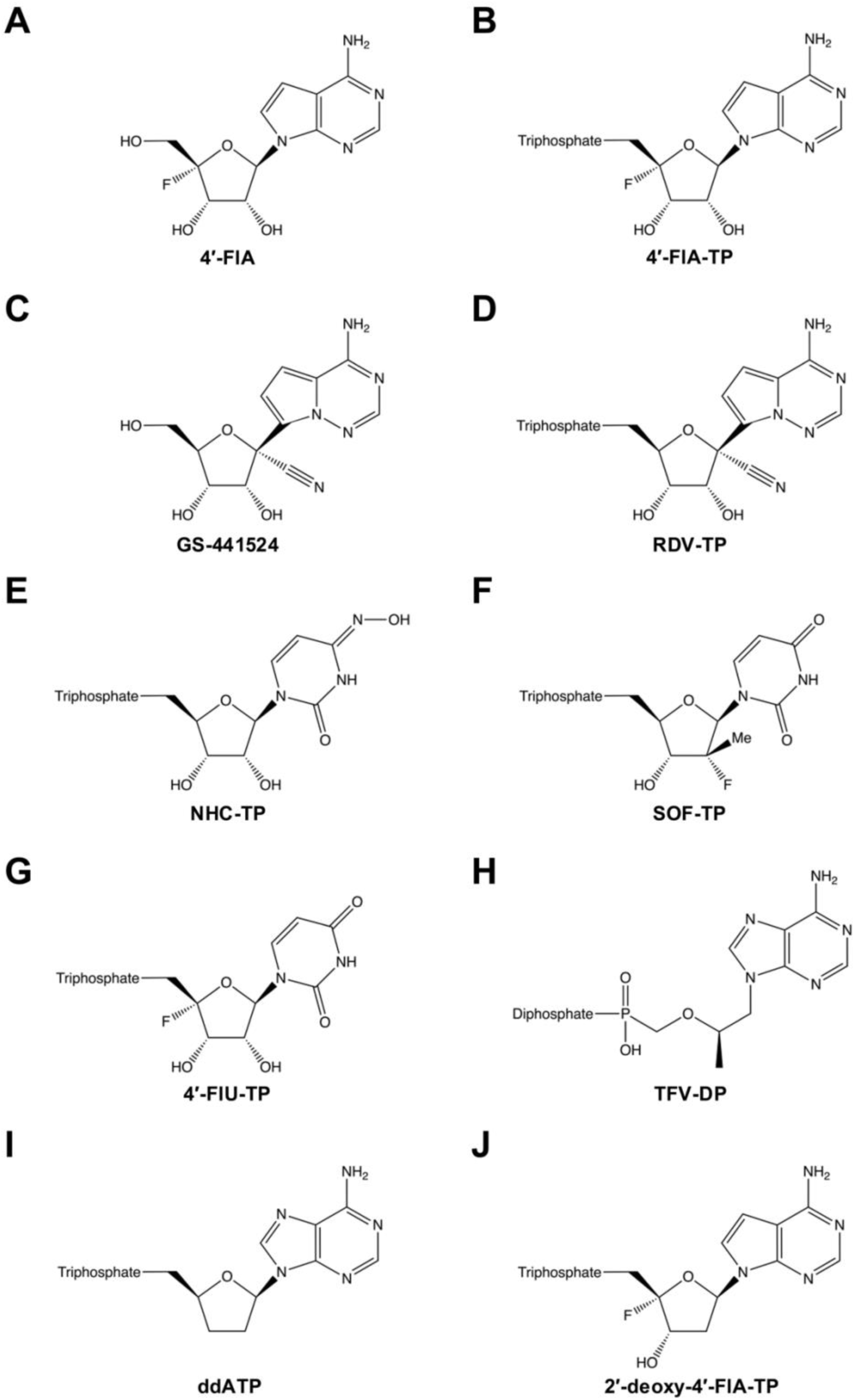
Chemical structures of nucleoside and nucleotide analogs investigated in this study. 4′-fluoroadenosine (4′-FlA) (**A**), 4′-fluoroadenosine triphosphate (4′-FlA-TP) (**B**), GS-441524 (**C**), RDV-TP (**D**), the active form of molnupiravir or NHC-TP (**E**), the active form of sofosbuvir or SOF-TP (**F**), the active form of 4′-FlU or 4′-FlU-TP (**G**), the active form of tenofovir or TFV-DP (**H**), ddATP (**I**), and 2′-deoxy-4′-FlA-TP (**J**).

## Materials and methods

### Chemicals

NTPs were purchased from GE Healthcare (Mississauga, ON, Canada). [α-^32^P]-GTP was purchased from PerkinElmer (Revvity, Waltham, MA, USA). NHC-TP and baloxavir acid were purchased from MedChemExpress (Monmouth Junction, NJ, USA). ddATP was purchased from Roche (Mannheim, Germany). kCMN996 (RDV, GS-441524) was purchased from Combi Blocks (San Diego, CA, USA). TFV-DP was purchased from Toronto Research Chemicals (Vaughan, ON, Canada). mCOT466 (4′-FlA), mCOU991 (4′-FlA-TP), mCQT310 (2′-d-4′-FlA-TP), and kCOT700 (4′-FlU-TP) were synthesized as follows: mCOT466 (4′-FlA): WO2024227159 – 4′-substituted nucleosides and nucleotides as antiviral agents – Google Patents (35); mCOU991 (4′-FlA-TP): See “**Synthetic chemistry**” in supporting information (Figure S1); mCQT310 (2′-d-4′-FlA-TP): See “**Synthetic chemistry**” in supporting information (Figure S2); kCOT700 (4′-FlU-TP): Synthesized according to a previously published protocol (36).

### DENV-2, HRV-16, SARS-CoV-2, RSV, and FluB cell-based assays

The preparation of dengue virus (Dengue virus type 2 or DENV-2 16681 strain), human rhinovirus (HRV-16), SARS-CoV-2 (USA/WA1/2020), and RSV-A (A2 strain) viral stocks and subsequent assessment and analysis of the antiviral activity and cytotoxicity of mCOT466 (4′-FlA) against these viruses in various cell lines have been previously described in Google Patent WO2024227159, “4′-substituted nucleosides and nucleotides as antiviral agents” (35). The assay for evaluating the efficacy of 4′-FlA against FluB in Madin-Darby canine kidney (MDCK) cells has been previously described (37).

### EBOV antiviral assays

HeLa cells (ATCC) were seeded at 4,000 cells/well 24h prior to infection. Experimental compounds were prepared as 5mM stocks in Dimethyl sulfoxide (DMSO, Sigma-Aldrich) and further diluted to 200 µM. Cell culture media was aspirated and replaced with 40 µL of DMEM (Dubecco’s Modified Eagle Medium, Gibco) containing 2% FBS (Fetal Bovine Serum, GeminiBio) and 100 units/mL of P/S (Penicillin-Streptomycin, Gibco). Compounds were dispensed onto the cell-containing plates using an iDOT liquid handler (Dispendix) in a 10-point dose-response curve in triplicate, ranging from 20 µM to 0.04 µM. All wells were backfilled to a final volume of 0.2 μL DMSO. Outer wells were excluded from compound addition due to edge effects. Following compound treatment, cells were challenged with Ebola Mayinga virus (EBOV; isolate Ebola virus/H.sapiens-tc/COD/1976/Yambuku-Mayinga) within the BSL-4 laboratory. Virus inoculum was prepared in DMEM supplemented with 2% FBS and 1xPenicillin/Streptomycin solution to yield a final multiplicity of infection (MOI) of 0.2, and 10 µL was added per well. The plates were then incubated for 40 h at 37°C before being inactivated in 10% formalin. Following inactivation by formalin, plates were removed from the BSL-4 laboratory and washed in 1x phosphate-buffered saline (PBS, Thermo Fisher Scientific) permeabilized with 0.1% (v/v) Triton X (Acros Organics) for 15 min and blocked with 3.5% bovine serum albumin (BSA, 1 h). Cells were stained with mouse anti-EBOV glycoprotein (4F3, IBT Bioservices) for 3 h at room temperature. Following a wash with 1xPBS, the secondary antibody, donkey anti-mouse AlexaFluor 488 (Invitrogen), was added and incubated overnight at 4°C. Plates were then washed and counterstained with Hoechst 33342 (Thermo Fisher Scientific). Plates were imaged using a Cytation 1 automated microscope (BioTek) using the blue and green fluorescence channels. Image analysis was performed using CellProfiler (Broad Institute, v4.2.6), where pipelines were configured to quantify total EBOV-infected cells and nuclei. Infection rates were calculated for each well and normalized to the DMSO control wells. Cell viability was assessed per well, with values normalized to the DMSO controls. Dose-response curves were generated using GraphPad Prism (v10.2.3) and the four-parameter logistic model ([agonist] vs response, variable slope). 4′-FlA was also screened against the EBOV minigenome assay as previously described (38).

### Rescue of recombinant CCHFV and subsequent dose-response inhibition

Recombinant Crimean-Congo hemorrhagic fever virus expressing the Zs-Green fluorescent protein (rCCHFV-ZsGreen) was rescued as previously described (39,40). Briefly, baby hamster kidney cells (BHK-21; ATCC CCL-10) were seeded in 6-well plates (1 × 10⁶ cells per well) one day prior to transfection. Culture medium was removed and replaced with Opti-MEM for plasmid transfection. Cells were transfected with a pT7 plasmids encoding the CCHFV IbAr10200 gene segments S, M, and L (1 µg, 2.5 µg, and 1 µg, respectively), along with pCAGGS helper plasmids encoding CCHFV NP and L proteins, and a codon-optimized bacteriophage T7 RNA polymerase (0.66 µg, 0.33 µg, and 1 µg, respectively), using Lipofectamine 3000 () according to the manufacturer’s instructions. The S genome segment contains Zs-Green inserted upstream of the NP gene separated by a porcine teschovirus-1 2A peptide linker sequence (P2A). Plasmids were provided by Éric Bergeron and César G. Albariño (Centers for Disease Control and Prevention, CDC). At 24 hours post-transfection (hpt), Opti-MEM was replaced with cell culture medium. Cell culture supernatants (CCS) were collected at 96 hpt and used to infect African green monkey kidney epithelial cells (Vero E6; ATCC CRL-1586) in T75 flasks for virus amplification and generation of viral stocks.

Vero E6 cells were seeded in 96-well clear-bottom black plates (2 x 10^4^ cells/well) in quadruplicate one day before infection. Cells were infected with rCCHFV Zs-Green at a MOI of 0.005. Following 1 h of viral adsorption, media containing 2-fold serial dilutions of inhibitors (starting at 50 µM), were added directly to the viral inoculum. At 72 h post-infection (hpi), cells were inactivated with 10% formalin and stained with DAPI. Fluorescence intensities of Zs-Green and DAPI were quantified using a GloMax plate reader (Promega) and Zs-Green signal was normalized to mock-treated, infected cells, whereas DAPI signal was normalized to mock-treated, mock-infected cells for EC_50_ and CC_50_ determinations, respectively. All work involving rCCHFV were performed in BSL-4 facilities at the Texas Biomedical Research Institute (San Antonio, TX, USA).

### FluA/A549 high-content imaging assay and cytotoxicity counter screen

Influenza A virus (FluA) infection was quantified by immunofluorescence-based high-content imaging multi-cycle assay as previously described for influenza antiviral screening assays, with minor modifications (41). Compounds were acoustically transferred using a Labcyte Echo liquid handler into collagen I-coated 384-well µclear-bottom plates (Greiner, 781090-2B). A549 cells were mixed with Influenza A virus A/WSN/1933 (H1N1) (WSN/33) at a MOI of 0.01 and seeded at 4.0 x 10³ cells per well in 30 µL F-12K medium supplemented with 1% BSA (Sigma), and 0.5 µg/mL TPCK-treated irradiated trypsin (Worthington). Plates were incubated for 48 h at 37°C with 5% CO₂. Cells were fixed with 4% formaldehyde for 15 min at room temperature, washed with PBS containing 0.05% Tween-20 (PBS-T), permeabilized with 0.25% Triton X-100 for 10 min, and blocked with SuperBlock T20 (Thermo Fisher Scientific) for 1 h. Plates were incubated overnight at 4°C with goat anti-influenza A polyclonal antibody (United States Biological, I7650-05E-ML550; 1:1,000 dilution) and 8 µM DAPI in blocking buffer. After washing with PBS-T, plates were sealed and imaged using an ImageXpress Micro Confocal High-Content Imaging System (Molecular Devices) with a 10x objective (four fields per well). Images were analyzed using MetaXpress (v6.7.2.290). DAPI staining was used to identify total nuclei, and influenza A immunofluorescence was used to quantify infected cells. For cytotoxicity assessment, A549 (ATCC CCL-185) cells were maintained at 37°C with 5% CO₂ in humidified incubators and cultured in F-12K medium supplemented with 10% heat-inactivated FBS, 100 IU/mL penicillin, and 100 µg/mL streptomycin. 300 A549 cells per well in assay media (same as maintenance media but with 2% FBS) were seeded in 1,536-well white tissue culture-treated plates (Corning 9006BC) containing compounds transferred acoustically in three-fold serial dilutions starting at 80 µM. Cells were incubated for 48 hours, after which viability was quantified using CellTiter-Glo (Promega G7573; 2 µL of 50% reagent diluted in water) and luminescence measured on a CLARIOstar plate reader (BMG Labtech). High-content image data and cytotoxicity data were processed using Genedata Screener (v21.0.3). IAV/A549 assay data were normalized to DMSO controls and inhibitor controls (0.2 µM baloxavir for antiviral activity and 10 µM puromycin for infected-cell toxicity). For cytotoxicity assays, 80 µM puromycin served as the positive control. Dose-response curves were generated from technical triplicates using a four-parameter Hill equation.

### Nucleic acids

The following synthetic oligos were used in this study. The portion of the template that is underlined is complementary to the primer used. Oligos requiring radiolabeling were 5′-monophosphorylated. For assays with most viral RdRps, the RNA template 3′-<u>UGCG</u>CUAGAAAAAAp-5′ was used for analog selectivity, pattern of inhibition, and UTP incorporation at position “i + 1” experiments. Analog selectivity, pattern of inhibition, and UTP incorporation at position “i + 1” experiments for EBOV RdRp were evaluated using 3′-<u>UGCG</u>CUAGUUUAUUp-5′. Pattern of inhibition for RSV RdRp was also evaluated using 3′-<u>UGCG</u>CUAGUUUAUUp-5′. 3′-<u>UGCG</u>CUUUUUXUUGUUGUUUp-5′ was used for the evaluation of nucleotide incorporation opposite a template-embedded analog in comparison to the natural scenario, where X represents either templated adenosine (Template “A”) or templated 4′-FlA (Template “F”). These RNA templates were produced as previously described (42). For h-mtRNAP, the DNA template 3′-<u>GCGCGGT</u>TGCAAAAAAA-5′ and the RNA primer 5′-Cy5-UUUUGCCGCGCCA-3′ were used for 4′-FlA-TP selectivity, pattern of inhibition, and CTP incorporation at position “i + 1” experiments. 3′-<u>GCGCGGT</u>GACTATTATTA-5′ DNA template and 5′-FAM-UUUUGCCGCGCCA-3′ RNA primer were used for NHC-TP selectivity. 3′-<u>GCGCGGT</u>AGCTTTTTTT-5′ DNA template and 5′-Cy5-UUUUGCCGCGCCA-3′ RNA primer were used for 4′-FlU-TP selectivity. For DNA Pol γ and KF, 3′-TAAGAC<u>TGATTTTCCCAGACTCCCTA</u>TCAGAGAACG-5′ DNA template and 5′-Cy5-ACTAAAAGGGTCTGAGGGAT-3′ DNA primer, based on Baldwin *et al*. (43), were used for analog selectivity and pattern of inhibition experiments. 5′-monophosphorylated RNA primers and templates were purchased from Dharmacon (Lafayette, CO, USA). 5′-triphosphorylated RNAs were purchased from ChemGenes (Wilmington, MA, USA), then capped and radiolabeled using [α-^32^P]-GTP, the mRNA cap 2′ O-methyltransferase, and vaccinia capping system from New England BioLabs (Whitby, ON, Canada). 5′-fluorescently labelled (Cy5) primers and their complementary templates were purchased from IDT (Coralville, IA, USA).

### Expression and purification of enzymes

The design rationale, expression, and purification of DENV-2, HRV-16, SARS-CoV-2, RSV, EBOV, CCHFV, FluB, h-mtRNAP, and Pol γ used in this study have been previously described (12,42,44–49). DNA polymerase Klenow fragment (3′→5′ no exonuclease) was purchased from New England Biolabs (M0212S, Ipswich, MA, USA). The baculovirus expression system was also employed for the FluA H3N2 PA/PB1/PB2 complex. The pFastBac-1 (Invitrogen, Burlington, ON, Canada) plasmid with codon-optimized synthetic DNA sequences (GenScript, Piscataway, NJ, USA) encoding for FluA PA (PA_AAA43597.1), PB1 (PB1_AAA43579.1), and PB2 (PB2_AAA43613.1) was designed and used as starting material for protein expression, based on previously reported by others (50–52). We utilized the MultiBac (Geneva Biotech, Indianapolis, IN, USA) system for protein expression in Sf9 insect cells (Invitrogen) according to the published protocols (53,54). The FluA RdRp components (PA, PB1, PB2) were expressed as individual recombinant proteins controlled by individual polyhedrin promoters. The resulting FluA RdRp complex is comprised of the PA subunit, the PB1 subunit, and the PB2 subunit with a C-terminal 8xHis tag and Protein A tag. Sf9 cell pellets expressing FluA RdRp complex were lysed, and the polymerase was purified as previously reported (49,55). Protein purity and identity were confirmed by sodium dodecyl-sulfate polyacrylamide gel electrophoresis (SDS-PAGE) and mass spectrometric analysis (Dr. Jack Moore, Alberta Proteomics and Mass Spectrometry Facility, University of Alberta).

### In vitro gel-based assays

RNA synthesis assays, data acquisition, and quantification were performed as we previously described (44–46,48,55). Reactions containing viral RdRp and non-5′-labeled primer were provided with 0.1 µM [α-^32^P]-GTP for visualization of RNA synthesis. For experiments determining kinetic parameters, enzyme concentrations and the time point for each reaction were optimized to ensure incorporation of the first nucleotide was within the linear range of product formation. Standard reaction mixtures contained purified polymerase complex, 25 mM Tris-HCl pH 8.0, 200 µM primer, 2 µM template (10 µM for CCHFV in analog selectivity, pattern of inhibition, and “i + 1” experiments), 0.1 µM [α-^32^P]-GTP, 0.5 mM EDTA (0.25 mM for HRV-16, SARS-CoV-2, EBOV, Pol γ, and KF), and various nucleotide concentrations. For EBOV, glycerol was added to reaction mixtures to equal 5%. For h-mtRNAP evaluation of 4′-FlA-TP and 4′-FlU-TP, 50 nM of primer and 100 nM of template were provided, while NHC-TP investigation utilized primer-template strands which were annealed before their addition into the mixture (50 nM) as previously described (44). For Pol γ and KF, 50 nM of pre-annealed primer-template was supplemented. Reactions with SARS-CoV-2, h-mtRNAP, Pol γ, and KF also contained 50 mM NaCl. Mixtures were incubated for 5 to 10 minutes at 23°C (h-mtRNAP), 30°C (viral RdRps), or 37°C (Pol γ and KF), and RNA synthesis was initiated by the addition of 2.5 mM (HRV-16, SARS-CoV-2, EBOV, h-mtRNAP, and Pol γ) or 5 mM (DENV-2, RSV, CCHFV, FluB, and KF) MgCl_2_ and incubated. The reaction duration varied among enzymes: DENV-2, HRV-16, RSV, and FluB reactions were stopped after 10 minutes; SARS-CoV-2 and Pol γ reactions were stopped after 2 minutes; KF reactions were stopped after 1 minute; CCHFV reactions were stopped after 15 minutes; EBOV reactions were stopped after 20 minutes; and h-mtRNAP reactions with 4′-FlA-TP and NHC-TP were stopped after 4 minutes, whereas reactions with 4′-FlU-TP were stopped after 8 minutes. The capped primer extension assay utilized purified FluA or FluB RdRp enzyme, 25 mM Tris-HCl pH 8.0, 50 mM NaCl, 5 mM MgCl_2_, 0.5 mM EDTA, 10 µM baloxavir acid (BXA), 1 µM vRNA (equimolar 5′ (5′ pAGUAGUAACAAGAG 3′) and 3′ (5′ UAUACCUCUGCUUCUGCU 3′) RNAs) incubated for 5 minutes at 30°C. Reactions were started with the addition of 0.1 µM cap1 radiolabeled primer (5’m^7^G-AmAUCUAUAAUAGC; synthesized according to manufacturer protocols and purified as described previously) and nucleotides at various concentrations, then allowed to proceed for 10 minutes at 30°C (56). All reactions were stopped by the addition of an equal volume of a formamide/EDTA (50 mM) mixture. Reaction products were then incubated at 95°C for 5 minutes and resolved by 20% polyacrylamide gel electrophoresis (PAGE) containing 8 M urea, 89 mM Tris base, 89 mM Boric acid, and 2 mM EDTA, using 1X TBE running buffer. Radiolabeled or fluorescent signals were then visualized using an Amersham Typhoon biomolecular imager (Cytiva) and quantified in Cytiva ImageQuant TL software. The data were analyzed using GraphPad Prism 10 (GraphPad Software, Inc). To calculate selectivity, the fraction of products generated at position “i” out of the total signal in the lane was determined and plotted against the concentration of the nucleotide or nucleotide analog. The kinetic parameter *V*_max_ was defined as the product fraction of extended primer relative to the total signal observed in each lane. *K*_m_ reflects the nucleotide concentration at half the *V*_max_ value. *V*_max_/*K*_m_ determines the efficiency of a single nucleotide incorporation for a given nucleotide. To compare the efficiency of nucleotide incorporation, a selectivity value is determined as the ratio of the incorporation efficiency of the natural nucleotide to the incorporation efficiency of the analog. Incorporation efficiency of the subsequent nucleotide following either NTP or analog incorporation (at position “i + 1”) was calculated as the fraction of RNA synthesized products generated at position “i + 1” and beyond, divided by the total signal in the lane, and plotted against the concentration of NTP incorporated at “i + 1”. Inhibition (fold) compares the efficiency of NTP incorporation following 4′-FlA-TP incorporation to the efficiency of NTP incorporation following ATP incorporation.

### Cryo-EM sample preparation

For cryo-EM of SARS-CoV-2 RdRp complex with RNA and 4′-FlA-TP, the nsp12, nsp7, and nsp8 subunits were expressed separately. The nsp12 gene with a C-terminal TEV cleavage site and a Twin-Strep-Tag was codon-optimized and inserted into the pEZT-BM vector. HEK293F cells were transfected with 1 mg DNA and 3 mg PEI per liter of cells. After 5 days, cells were harvested, and pellets were resuspended in lysis buffer containing 25 mM Tris, pH 7.4, 300 mM NaCl, 1 mM MgCl₂, 0.1% IGEPAL CA-630, 1 mM DTT, 10% glycerol, and protease inhibitor cocktail. Following cell lysis by sonication, the lysate was clarified by centrifugation at 30,000 xg for 30 minutes. The supernatant was applied to Strep-Tactin XT resin (IBA Lifesciences) for affinity purification. After extensive washing, the bound protein was eluted overnight with Buffer BXT (IBA Lifesciences) and further purified by size exclusion chromatography on a Superdex 200 10/300 column (Cytiva) equilibrated with buffer containing 25 mM Tris pH 7.4, 300 mM NaCl, 0.1 mM MgCl₂, and 1 mM DTT. The nsp7 and nsp8 subunits were expressed in BL21(DE3) *Escherichia coli* cells. Protein expression was induced with 1 mM IPTG at an OD₆₀₀ of 0.7, and cultures were incubated overnight at 18°C. Harvested cells were lysed by sonication in a buffer containing 10 mM Tris, pH 7.4, 300 mM NaCl, 30 mM imidazole, and 5 mM β-mercaptoethanol. His-tagged proteins were purified using Ni-NTA affinity chromatography followed by TEV protease cleavage and dialysis to remove the His-tag. For RdRp complex assembly, purified nsp12 was incubated overnight with nsp7 and nsp8 at a molar ratio of 1:5:5. The assembled complex was further purified by size exclusion chromatography and concentrated in a final buffer containing 25 mM Tris, pH 7.4, 100 mM NaCl, 2 mM MgCl₂, and 1 mM DTT. For cryo-EM sample preparation, the RdRp complex was incubated with dsRNA substrate (Integrated DNA Technologies; template strand: 5’-UUUUUUUUUUAUAACUUAAUCUCACAUAGC-3’; primer strand: 5’-GCUAUGUGAGAUUAAGUUAU-3’) as previously described by Yin et al. (57), and 4′-FlA-TP at a molar ratio of 1:2:10 for 2 hours at 4°C before grid preparation. Grid preparation was performed by mixing 3 μl of RdRp complex containing 4′-FlA-TP and dsRNA at 3 mg/ml with 0.5 μl of n-dodecyl β-D-maltoside (DDM) to achieve a final concentration of 0.00128%. The sample was immediately applied to glow-discharged 1.2/1.3 300-mesh UltraAuFoil grids and plunge-frozen using a Vitrobot Mark IV (Thermo Fisher Scientific) under controlled conditions at 4°C chamber temperature, 100% humidity, 10 seconds wait time, and 3 seconds blotting time before plunging into liquid ethane.

### Cryo-EM data collection and image processing

Cryo-EM data acquisition was performed on a 200 kV Glacios (Thermo Fisher Scientific) equipped with a Falcon 4i direct electron detector. Automated image collection was conducted using EPU software (Thermo Fisher Scientific) at a nominal magnification of 190,000× with a pixel size of 0.718 Å. A total of 7,824 movies were collected and processed using CryoSPARC (58). Motion correction with dose weighting and CTF estimation were performed using CryoSPARC Live. For image processing, particles were initially picked using blob picker followed by 2D classification. Template-based particle picking was then performed using selected 2D class averages as templates. Multiple rounds of 2D classification were conducted to obtain clean particle datasets. Selected particles were subjected to ab initio reconstruction followed by heterogeneous refinement to separate different conformational states. The 3D model corresponding to the RdRp complex with RNA was selected from heterogeneous refinement and subjected to non-uniform refinement. An additional 3D classification was performed to further clean the particle set. To select the RdRp complex containing the 4′-FlA molecule, a mask was applied to the active site, and 3D variability analysis (3DVA) was performed, resulting in 48,482 particles used for the final map reconstruction. Finally, local refinement was conducted with a mask applied to the core region of RdRp after removing flexible domains, yielding a final structure at 3.38 Å resolution.

### Model building and refinement

The atomic model was initially built using PDB: 7BV2 as a starting template (57). Model refinement was carried out through Rosetta relaxed refinement protocols (59), followed by iterative rounds of manual adjustment in Coot and real-space refinement using Phenix (60,61). Model quality was monitored throughout the refinement process by calculating EMRinger and MolProbity scores after each refinement cycle (62,63). Structural visualization and figure preparation were performed using ChimeraX and PyMOL software (64,65). Comprehensive model statistics and validation data are presented in Table S1 and Figure S3C and 3D.

### h-mtRNAP modelling studies

Modelling of 4′-FlA-TP and RDV-TP within the N-site of h-mtRNAP was performed using a previously determined cryo-EM structure (PDB: 9R96) (66). The incoming GTP was replaced with either 4′-FlA-TP and RDV-TP, and modelled manually using Coot and Phenix, for the analogs to maintain base-pairing and chelation with the catalytic Mg^2+^ ions at the enzyme active site (60,61). Structural visualization and figure preparation were performed using PyMOL software (65).

## Results

### Experimental strategy

– The main objective of this work was to evaluate the efficacy and selectivity of 4′-FlA against RNA viruses that represent families of epidemic or pandemic potential. Prototypic viruses from diverse families were considered: *Flaviviridae* (DENV-2), *Coronaviridae* (SARS-CoV-2), *Picornaviridae* (HRV-16), *Pneumoviridae* (RSV), *Filoviridae* (EBOV), *Nairoviridae* (CCHFV), and *Orthomyxoviridae* (FluA and FluB). Biochemical enzymatic assays and structural evaluation were used to assess 4′-FlA-TP (Figure 1C) incorporation and the mechanism of inhibition of RNA synthesis. RDV-TP (Figure 1D) was used for comparative purposes, and host polymerases were considered to evaluate the potential for off-target effects.

### Antiviral activity and cytotoxicity of 4′-FlA

4′-FlA demonstrates antiviral effects against a panel of prototypic viruses (Table 1). Antiviral efficacy by increasing effective inhibitory concentrations (EC_50_) values follows the order DENV-2 (0.016 µM) > RSV (0.061 µM) > FluA (0.067 µM) > SARS-CoV-2 (0.071 µM) > CCHFV (0.076 µM) > FluB (< 0.1 µM) > HRV-16 (0.14 µM) > EBOV (0.22 µM). Against the EBOV minigenome, 4′-FlA similarly demonstrates antiviral activity (0.17 µM). The 50% cytotoxic concentration (CC_50_) for 4′-FlA varies due to the cell types employed, ranging from 1.0 to >50 µM, and corresponding selectivity index (SI) values ranging from ∼10 to >100, indicating the observed antiviral activities are predominantly driven by polymerase inhibition.

**Table 1.** Antiviral activity and cytotoxicity of GS-441524 and 4′-FlA^a^.

| Virus | Cell line | GS-441524 |  |  | 4'-FIA |  |  |
| --- | --- | --- | --- | --- | --- | --- | --- |
|  |  | EC<sub>50</sub> (μM) | CC<sub>50</sub> (μM) | Selectivity index (SI)<sup>b</sup> | EC<sub>50</sub> (μM) | CC<sub>50</sub> (μM) | Selectivity index (SI) |
| DENV-2 | HepG2 | 4.9 ± 1.7<br>9.46<sup>1</sup> | > 99 | >20 | 0.016 ± 0.004 | 3.3 ± 0.8 | 210 |
| HRV-16 | HeLa | 14.6 ± 6.9<br>(n=2) | > 99 | >6.8 | 0.14 ± 0.16 (n=2) | 2.1 ± 0.7 | 15 |
| SARS-CoV-2 | HeLa-ACE2 | 8.7 ± 0.4<br>0.28 – 1.65<sup>2</sup> | > 99 | >11 | 0.071 ± 0.010 | 1.00 ± 0.03 | 14 |
| RSV | Hep2 | 0.89 ± 0.20<br>0.63<sup>3</sup> | > 99 | >110 | 0.061 ± 0.027 | 3.2 ± 1.0 | 52 |
| EBOV | HeLa | 0.17 (n=1) |  |  | 0.22 ± 0.06 (n=2) | 5.5 ± 1.1 | 25 |
|  |  | 0.51 ± 0.04<sup>c</sup> | 8.02 ± 0.04<sup>c</sup> | 16 | 0.17 ± 0.12<sup>c</sup> | 1.6 ± 0.1<sup>c</sup> | 9.4 |
|  |  | 1.0 – 3.1<sup>3</sup> | > 49<sup>3</sup> | 16 – 50<sup>3</sup> |  |  |  |
| CCHFV | Vero-E6 | No inhibition<sup>3</sup> |  |  | 0.076 (n=1) | >50 | >660 |
| FluA | A549 | >10 (n=2)<br>27.9<sup>1</sup> | >99<br>>30<sup>1</sup> | >9.9<br>>1.1<sup>1</sup> | 0.067 ± 0.008 | 2.8 ± 0.3 | 42 |
| FluB | MDCK | - | - | - | < 0.1 (n=1) | 30 | 300 |

<sup>a</sup> The antiviral activity of GS-441524 and mCOT466 was evaluated against a panel of viruses in different cell types as described in Materials and Methods to determine the half-maximum effective concentration (EC<sub>50</sub>) and half-maximal cytotoxic concentration (CC<sub>50</sub>). All data are at least $n \geq 3$ replicates (unless otherwise indicated) and reported as the average ± SD.
<sup>b</sup> Selectivity index = CC<sub>50</sub>/EC<sub>50</sub>.
<sup>c</sup> Half maximal inhibitory concentration (IC<sub>50</sub>) of 4'-FIA determined in the EBOV minigenome assay. Data are at least $n=2$ replicates and reported as the average ± SD.
<sup>1</sup> Data from Cho *et al.*, 2012 (67).
<sup>2</sup> Data from Pruijssers *et al.*, 2020 (13).
<sup>3</sup> Data from Lo *et al.*, 2017 (11).

GS-441524, the parent nucleoside of RDV, was also evaluated for comparative purposes (Figure 1C). In cell-based assays, EC_50_ values for RDV and GS-441524 against several of the viruses discussed here have been previously determined (10–14,67). EC_50_ values for RDV and GS-441524 are in the sub-to-low micromolar range across many viral families, though the prodrug has greater activity, likely due to higher levels of active 5′-triphosphate metabolite formation within the cell (13). Here, we determined EC_50_ values for GS-441524 against DENV-2, HRV-16, SARS-CoV-2, RSV, and EBOV (Table 1). Our observed EC_50_ values for GS-441524 are largely consistent with previous reports, as indicated in the Table. Taken together, although 4′-FlA demonstrates a broader spectrum of antiviral activities as compared to RDV, the lower CC_50_ values point to potential off-target effects.

### Incorporation of 4′-FlA-TP by representative viral polymerases

4′-FlA-TP, the active triphosphate form of 4′-FlA, is anticipated to compete with natural ATP pools for incorporation by the viral polymerase. Furthermore, previous studies have demonstrated that efficient incorporation of an NA is an important parameter to assess the potential for antiviral activity (45,55,68). We compared the incorporation efficiency of ATP and 4′-FlA-TP with purified recombinant polymerases and polymerase complexes from the viruses for which 4′-FlA was tested against (Table 2 and Table S2). For DENV-2, HRV-16, SARS-CoV-2, RSV, EBOV, CCHFV, and FluB, *in vitro* enzyme RNA synthesis was evaluated using short model primer/templates as previously described (12,45,46,48,55) (Figure S4). FluA RdRp RNA synthesis activity requires the use of a genomic vRNA template, utilizing a radiolabeled 13-nt 5′-methyl-7-guanosine capped primer (Figure S5). Steady-state kinetic parameters *V*_max_ and *K*_m_ were measured to determine substrate selectivity, defined as the ratio of the incorporation efficiency of natural nucleotide over the nucleotide analog. A selectivity value below one suggests that the analog is more efficiently incorporated than the natural nucleotide. The relative efficiency of 4′-FlA-TP incorporation follows the order HRV-16 (0.48) > SARS-CoV-2 (0.99) > CCHFV (1.4) ∼ FluB (1.4) > EBOV (4.2) > RSV (7.1) > DENV-2 (7.6) (Table 2). Using the 13-nt capped primer system, FluA and FluB demonstrate selectivity values of 1.5 and 0.52, respectively (Table 2 and Table S3). The low selectivity values for CCHFV, FluA, and FluB RdRp are a distinct property of 4′-FlA-TP in comparison to RDV-TP. Collectively, these data suggest that the 4′-fluoro modification is well tolerated across all viral polymerases tested and that 4′-FlA-TP can efficiently compete with the natural nucleotide ATP.

**Table 2.** Incorporation of RDV-TP and 4′-FlA-TP by selected viral RdRp enzymes.

| RNA Sense | Family | Virus | RDV-TP | 4'-FlA-TP |
| --- | --- | --- | --- | --- |
|  |  |  | Selectivity<sup>a</sup> (fold) |  |
| Positive ssRNA | *Flaviviridae* | DENV-2 | 11.1<sup>1</sup> | 7.6 |
|  | *Picornaviridae* | HRV-16 | 1.03<sup>2</sup> | 0.48 |
|  | *Coronaviridae* | SARS-CoV-2 | 0.28<sup>3</sup> | 0.99 |
| Nonsegmented negative ssRNA | *Pneumoviridae* | RSV | 2.7<sup>4</sup> | 7.1 |
|  | *Filoviridae* | EBOV | 4.0<sup>3</sup> | 4.2 |
| Segmented negative ssRNA | *Nairoviridae* | CCHFV | 41<sup>5</sup> | 1.4 |
|  | *Orthomyxoviridae* | FluB | 68<sup>5</sup> | 1.4 |
| Segmented negative ssRNA | *Orthomyxoviridae* | FluA* | 1800 | 1.5 |
|  |  | FluB* | 1400 | 0.52 |

<sup>a</sup>Selectivity of a viral polymerase for a nucleotide analog is calculated as the ratio of the $V_{\max}/K_m$ values for ATP and the analog, respectively.
<sup>1</sup>Data from Walker *et al.*, 2026 (12).
<sup>2</sup>Data from Gordon *et al.*, 2024 (45).
<sup>3</sup>Data from Gordon *et al.*, 2020 (47).
<sup>4</sup>Data from Tchesnokov *et al.*, 2019 (48).
<sup>5</sup>Data from Gordon *et al.*, 2022 (55).
\*Selectivity values generated for these viral RdRp were performed using a 13-nt labelled primer/template system, whereas all other values were generated using the unlabeled 4-nt primer/template system.

### Structure of incorporated 4′-FlA-MP by SARS-CoV-2 RdRp

We determined the cryo-EM structure of the SARS-CoV-2 RdRp complex comprising nsp12, nsp8, and nsp7 in complex with RNA and incorporated 4′-FlA-MP at an overall resolution of 3.4 Å (Figure 2 and Table S1). Structural analysis reveals that 4′-FlA-MP occupies the +1 or “i” active site position, equivalent to the binding site previously described for RDV in the SARS-CoV-2 RdRp (57)(Figure 2B, 2C, and S3A, S3B). At this position, 4′-FlA forms a base pair with the U10 nucleotide of the template strand (Figure 2C). Although the precise orientation of the 4′-fluorine modification on 4′-FlA cannot be unambiguously defined due to limited local resolution, it is positioned in close proximity to residues S759 and N691 of nsp12 that are also involved in RDV-TP binding (69). Comparisons to structures of DENV2 RdRp (PDB: 7XD8) (70) and FluB RdRp (PDB: 6QCT) (71) suggest that 4′-FlA could bind in a similar pose at their respective active sites, further highlighting the potential for broad-spectrum activity (Figure S3E, S3F).

**Figure 2.**
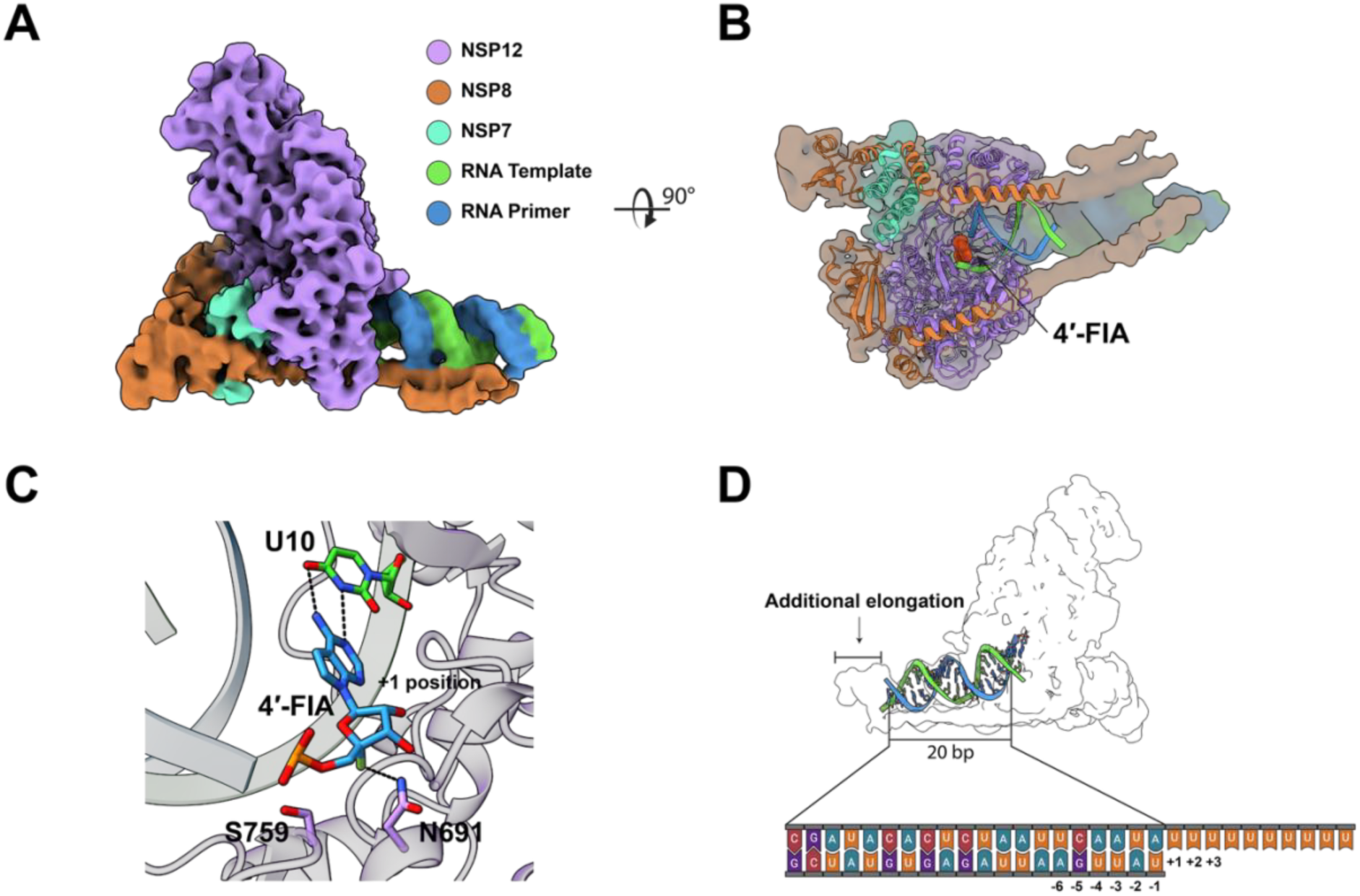
Cryo-EM structure of the SARS-CoV-2 RdRp complex bound to RNA and 4′-FlA. (**A** and **B**) Cryo-EM density map of the SARS-CoV-2 RdRp complex, low-pass filtered to 5 Å. The left panel (**A**) shows a side view, and the right panel (**B**) shows a bottom view of the complex. The atomic model is displayed as a cartoon representation fitted within the transparent surface map. 4′-FlA is shown as a red sphere model at the active site. (**C**) Close-up view of the RdRp active site showing 4′-FlA at the +1 position, base-paired with the template strand nucleotide U10. Hydrogen bonds between 4′-FlA and U10 are indicated by dashed lines. Residue N691 of NSP12, located within 3.5 Å of 4′-FlA, is indicated by dashed lines and labelled. (**D**) Schematic representation of the RNA scaffold used for cryo-EM structure determination. The 20 bp double-stranded region is indicated. Additional elongation beyond the designed dsRNA region is shown, with overhanging uracil residues on the template strand. Nucleotide positions are labelled (−6 to −1 and +1 to +3).

### Heterogeneous effects of 4′-FlA-TP in the primer-strand

Incorporation of an NA into the growing RNA primer-strand is commonly required for downstream inhibition. We evaluated the patterns of RNA synthesis following the incorporation of a single 4′-FlA-TP (Figure 3). For DENV-2 RdRp, 4′-FlA-TP does not appear to inhibit primer extension (Figure 3B). Similarly, 4′-FlA-TP demonstrates no inhibitory effect during primer-strand synthesis against HRV-16 and SARS-CoV-2 (Figure S6). In contrast, 4′-FlA-TP incorporation elicits immediate chain termination against RdRp from FluB and CCHFV (Figure 3C, Figure S7). 4′-FlA-TP incorporation by RSV RdRp appears to elicit an inhibitory effect downstream at position “i+3” (Figure S8A). A template allowing for a second 4′-FlA-TP incorporation enhances the inhibitory effect at this position (Figure S8B), which is also observed for EBOV (Figure S9). Using the 13-nt radiolabeled capped primer and vRNA template, 4′-FlA-TP incorporation results in immediate chain termination against both influenza RdRp (Figure S10). A second inhibitory site at position “i + 2” can also be seen, which may be due to a second incorporation event of 4′-FlA-TP. Notably, against all viral RdRp, longer RNA products past the site of a single incorporation of 4′-FlA-TP can be generated. As observed with influenza, RSV, and EBOV RdRps, inhibition due to multiple incorporations appears more absolute.

**Figure 3.**
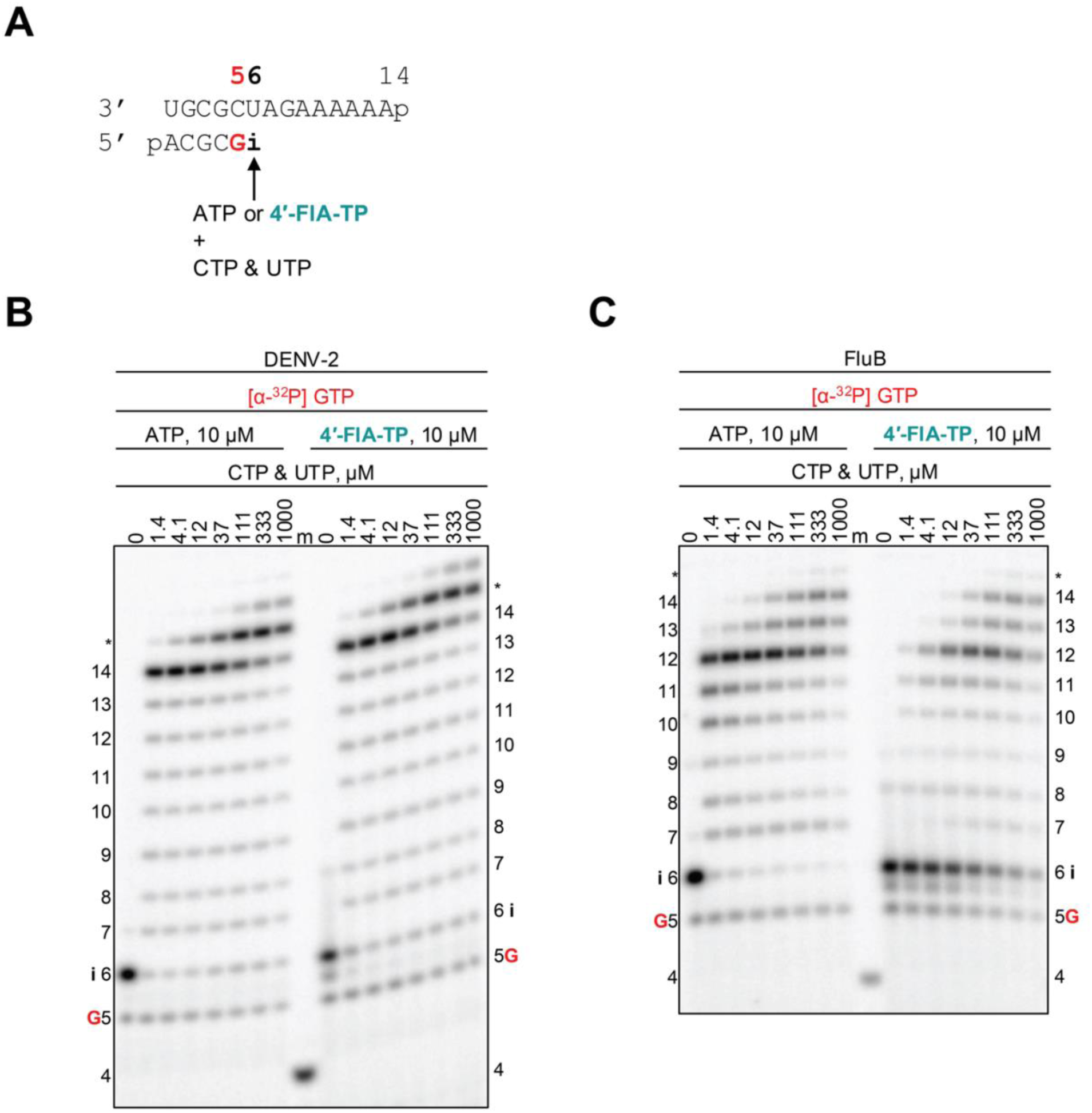
DENV-2 and FluB RdRp-catalyzed RNA synthesis and inhibition patterns following a single incorporation of ATP or 4′-FlA-TP as a function of nucleotide concentration. (**A**) RNA primer/template supporting a single incorporation of ATP or 4′-FlA-TP at position 6 (“i”). G indicates incorporation of [α-^32^P]-GTP at position 5. Migration pattern of RNA products resulting from (**B**) DENV-2 RdRp- and (**C**) FluB RdRp-catalyzed RNA extension following incorporation of ATP or 4′-FlA-TP at position 6 (“i”) at increasing CTP and UTP concentrations. A 5′-^32^P-labelled 4-nt primer (4) serves as a size marker (m). RNA products formed at and beyond the asterisk may be a result of sequence-dependent slippage events or RdRp nucleotide transferase activity.

To translate these findings into quantitative terms, we determined the inhibitory efficiency of subsequent nucleotide incorporation following the incorporation of 4′-FlA-TP. We determined the kinetic parameters of UTP incorporation at position “i + 1” with AMP- and 4′-FlA-MP-terminated primer strands to determine inhibition values (Figure S11)(Table S4). For the positive-sense RNA viruses, the extension of 4′-FlA-terminated primers is only slightly impaired, 3- to 5-fold, compared to AMP. Against RSV and EBOV, the 4′-FlA-terminated primer displays 13- and 8-fold inhibition. For CCHFV, UTP incorporation with 4′-FlA-MP-terminated primers is inhibited roughly 33-fold. Against FluB, the extension of 4′-FlA-MP-terminated primer requires a marked 710-fold increase in UTP concentration. Thus, 4′-FlA-TP exhibits variable inhibition on subsequent nucleotide incorporation. Overall, inhibition within the primer-strand by a single incorporated 4′-FlA-TP appears heterogeneous and not absolute.

In our cryo-EM structure, the RNA density observed in the RdRp-bound complex extends beyond the 20-base-pair double-stranded region used for structure determination, indicating that additional elongation had occurred (Figure 2D). The reaction between the overhanging uracil residues on the template strand and 4′-FlA incubated with the RNA demonstrates that 4′-FlA supports continued primer extension activity. In contrast to RDV against SARS-CoV-2, which exhibits delayed-chain termination due to a clash between the 1′-cyano modification and S861 (42,47,72,73), 4′-FlA appears to mediate a distinct mechanism in the primer-strand. To further evaluate potential steric conflicts introduced by the 4′-fluoro modification at each RNA position, we performed structural modelling using the previously determined 2.5 Å resolution structure (PDB: 7BV2) (57) (Figure S3G, S3H). In the context of the primer-strand, the 4′-fluorine atom is positioned within 3 Å of the main chain atom of C813 and the side chain of D865 at the -2 and -3 positions, respectively, indicating a potential for steric clash. In contrast, when the 4′-fluoro modification is modelled into the template strand, the fluorine atom falls within 3 Å of the main chain atoms of G683, G590, and S592, as well as the side chains of N543 and F594, suggesting a higher likelihood of steric conflict at these positions. Taken together, these observations indicate that the presence of the 4′-fluoro modification within the template strand of SARS-CoV-2 RdRp could interfere with polymerase activity.

### Template-dependent inhibition by an embedded 4′-FlA residue

Efficient incorporation of 4′-FlA-TP into the growing primer-strand and the ability of the viral RdRp to overcome the inhibitory effects provide the opportunity for the RdRp to synthesize full-length RNA copies. Thus, newly synthesized RNA copies could contain embedded analog residues that affect RNA synthesis when used as a templating-base. This mechanism of inhibition has been studied for RDV against numerous viral RdRps and provides a unifying mechanism of inhibition that describes its broad-spectrum antiviral activity (12,42,45,55). Here, we designed RNA templates with a single AMP or 4′-FlA-MP at position 11 (Figure 4A and Figure S12A). For all viral RdRp, low micromolar concentrations (1-10 µM) of NTPs are sufficient to generate full-length product on Template “A” (Figure 4B-D and Figure S12B-D). Comparatively, on Template “F”, inhibitory sites prior to the embedded 4′-FlA-MP and a corresponding decrease in full-template length RNA are seen. This inhibitory effect is maintained despite increasing NTP concentrations, though an increase in full-length product is observed. Further evaluation demonstrated that UTP incorporation opposite a template-embedded 4′-FlA-MP is most efficient among all NTPs; however, this incorporation is reduced relative to a template-embedded AMP at the equivalent position (Figure S13). Thus, 4′-FlA impairs the incorporation of incoming UTP when it is embedded in the template, providing a unifying mechanism of action that helps to explain the broad spectrum of antiviral activity of 4′-FlA.

**Figure 4.**
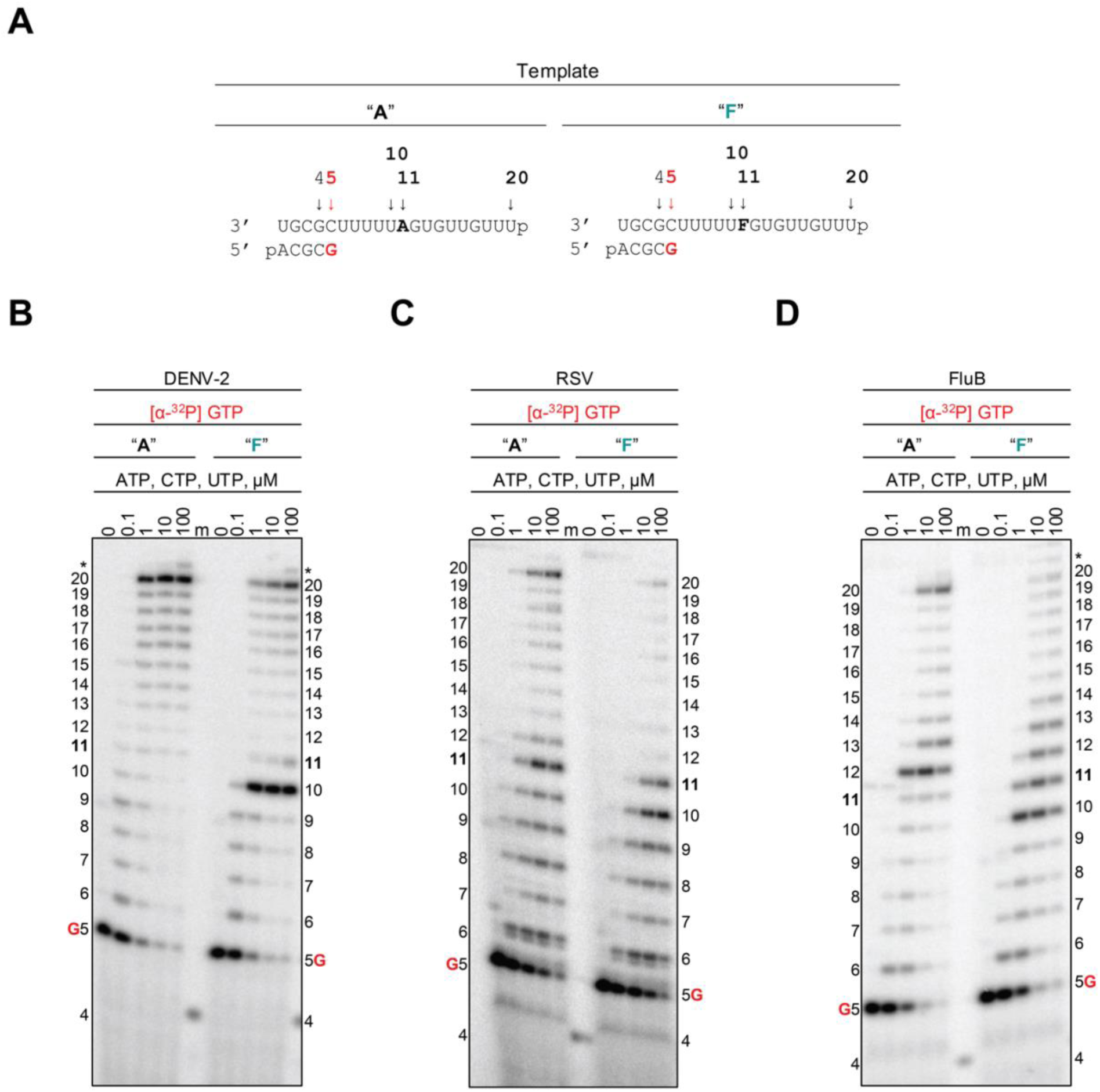
Template-dependent inhibition of DENV-2, RSV, and FluB RdRp by a single embedded 4′-FlA residue at position 11. (**A**) RNA primer/template with an embedded adenosine (Template “A”, *left*) and (Template “F”, *right*) at position 11. G indicates incorporation of [α-^32^P]-GTP at position 5. Migration pattern of RNA products resulting from (**B**) DENV-2 RdRp-, (**C**) RSV RdRp-, and (**D**) FluB RdRp-catalyzed RNA synthesis supplemented with increasing NTP concentrations using Template “A” (*left*) and Template “F” (*right*). A 5′-^32^P-labelled 4-nt primer (4) serves as a size marker (m). RNA products formed at and beyond the asterisk may be a result of sequence-dependent slippage events or RdRp nucleotide transferase activity. DENV-2, dengue virus; RSV, respiratory syncytial virus; FluB, influenza B virus; RdRp, RNA-dependent RNA polymerase.

### Incorporation of nucleotide analogs by host enzymes

Incorporation of NAs by host polymerases can be associated with cytotoxic effects (74–77). Here, we examined the use of 4′-FlA-TP as a substrate for h-mtRNAP (Figure S14, Table S5), comparing selectivity with approved NAs (Figure 1). Included were RDV-TP, NHC-TP, and sofosbuvir (SOF), a 2′-*α*-fluoro, 2′-*β*-methyluridine monophosphate prodrug approved for the treatment of hepatitis C virus (HCV) infection (78,79). Our selectivity values of RDV-TP, NHC-TP, and SOF-TP for h-mtRNAP are consistent with published data. RDV-TP is incorporated 2200-fold less efficiently than ATP, consistent with previous reports (48) (Table 3). These data demonstrate that RDV-TP is a poor substrate for h-mtRNAP, indicating low potential cytotoxicity and corroborating the observed CC_50_ values. For NHC-TP, we determined a selectivity of 22, similar to previous values of 11.5-13.2 and confirming that NHC-TP can be incorporated by h-mtRNAP (80). A selectivity for SOF-TP with h-mtRNAP could not be determined, consistent with SOF-TP being very poorly incorporated by h-mtRNAP (81,82). By comparison, 4′-FlU-TP is used very efficiently by h-mtRNAP, with a selectivity of 4.1. Similarly, h-mtRNAP demonstrates efficient incorporation of 4′-FlA-TP, with a selectivity of 8.4. Efficient incorporation of 4′-FlA-TP by h-mtRNAP may indicate potential cytotoxic off-target effects, which could help interpret the observed CC_50_ values. However, NHC-TP similarly demonstrates efficient incorporation by h-mtRNAP. Indeed, while the incorporation of an analog by h-mtRNAP is an indicator of potential cytotoxicity, it is not the sole determinant, and the threshold for incorporation to elicit toxicity is unknown.

**Table 3.** Incorporation of nucleotide analogs by off-target polymerases.

| Replication strategy | Family | Enzyme | Compound | Selectivity<sup>a</sup> |
| --- | --- | --- | --- | --- |
| Human DNA-dependent RNAP | Human | h-mtRNAP | RDV-TP | 2200<br>508<sup>1</sup> |
|  |  |  | SOF-TP | >5000 |
|  |  |  | NHC-TP | 22 |
|  |  |  | 4'-FIU-TP | 4.1 |
|  |  |  | 4'-FlA-TP | 8.4 |
| Human DNA-dependent DNAP | Human | Pol $\gamma$ | ddATP | 0.66 |
|  |  |  | TFV-DP | 210 |
|  |  |  | 2'-deoxy-4'-FlA-TP | 0.94 |
| *E. coli* DNA-dependent DNAP | *E. coli* | Klenow -exo | ddATP | 750 |
|  |  |  | TFV-DP | >5000 |
|  |  |  | 2'-deoxy-4'-FlA-TP | 4.8 |

<sup>a</sup> The selectivity of a polymerase for a nucleotide analog is calculated as the ratio of the $V_{\max}/K_m$ values for the natural NTP and the analog, respectively.
<sup>1</sup> Data from Tchesnokov *et al.* 2019 (48).

Based on an existing cryo-EM structure of the h-mtRNAP transcription initiation complex (66), we generated a model of 4′-FlA-TP incorporation at the h-mtRNAP NTP-binding site (N-site) (Figure S15). The 4′-fluoro modification of 4′-FlA-TP fits well within the adjacent molecular surface of h-mtRNAP (Figure S15C), supporting the efficient use of 4′-FlA-TP as a substrate. In contrast, the 1′-cyano substitution on RDV-TP clashes with h-mtRNAP residues (Figure S15D), indicating poor incorporation with h-mtRNAP. This aligns with previous modelling and the inefficient incorporation of RDV-TP (45).

Ribonucleoside analogs can be intracellularly converted to serve as substrates for host DNA polymerases. Ribonucleoside 5′-diphosphate (DP) is a metabolic intermediate in the activation pathway that can be used as a substrate by ribonucleotide reductase (RNR)(83). RNR is essential for the production of 2′-deoxyribonucleoside 5′-triphosphates used in DNA synthesis. Previous work demonstrated that NHC-DP could transit the RNR pathway, raising concerns about the potential mutagenic risk in humans (84). Therefore, we evaluated 2′-deoxy-4′-FlA-TP (2′-d-4′-FlA-TP) against the human DNA polymerase gamma (Pol γ)(Figure S16), and *E.coli* DNA polymerase I (Klenow Fragment, KF) to further assess potential cytotoxicity (Table 3 and Table S5). KF was included as an additional measure of off-target evaluation. The active metabolite forms of dideoxyinosine (ddI) and tenofovir (ddATP and TFV-DP, respectively) were included as benchmarks (Figure 1) (85–88). Against Pol γ, ddATP is incorporated very efficiently (selectivity of 0.66), whereas TFV-DP is incorporated 210-fold less efficiently than dATP. This outcome of substrate utilization by Pol γ is consistent with previous literature (75,76). KF does not incorporate TFV-DP and selectively incorporates dATP 750-fold more efficiently than ddATP. Comparatively, we found Pol γ incorporates 2′-d-4′-FlA-TP with a selectivity of 0.94, and KF incorporates 2′-d-4′-FlA-TP 4.8-fold less efficiently than dATP. The ability of 4′-FlA-TP to be utilized as a substrate for RNR is unknown. Should 4′-FlA-TP be efficiently converted to 2′-d-4′-FlA-TP, there is potential for incorporation into host DNA. Ultimately, these findings suggest that, in addition to the efficient incorporation by viral polymerases, 4′-FlA lacks target specificity.

### Inhibition of host polymerases is not evident

While efficient incorporation of 4′-FlA-TP and 2′-d-4′-FlA-TP by human and bacterial polymerases is indicative of off-target effects, cellular RNA synthesis may not necessarily be inhibited. We evaluated the patterns of RNA or DNA synthesis following incorporation of a single 4′-FlA-TP or 2′-d-4′-FlA-TP. For h-mtRNAP DdRp-catalyzed RNA synthesis, no discernible inhibitory effect in the primer-strand is observed following 4′-FlA-TP incorporation at position 14 (“i”) (Figure 5A and 5B). We also evaluated the potential inhibitory effect of subsequent nucleotide incorporation and found that incorporation of CTP following 4′-FlA-TP incorporation is very subtly reduced by 1.8-fold relative to the natural scenario (Table S4). Similarly, no apparent inhibition is observed in the primer-strand following 4′-FlU-TP incorporation (Figure 5C and 5D). For Pol γ and KF, 2′-d-4′-FlA-TP shows no visible inhibition on primer-extension reactions (Figure S17). In comparison, ddATP acts as an immediate chain terminator, disrupting DNA synthesis at the site of incorporation (Figure S17B). Using the assays described above, inhibition of primer-extension reactions is not evident with 4′-fluorinated compounds.

**Figure 5.**
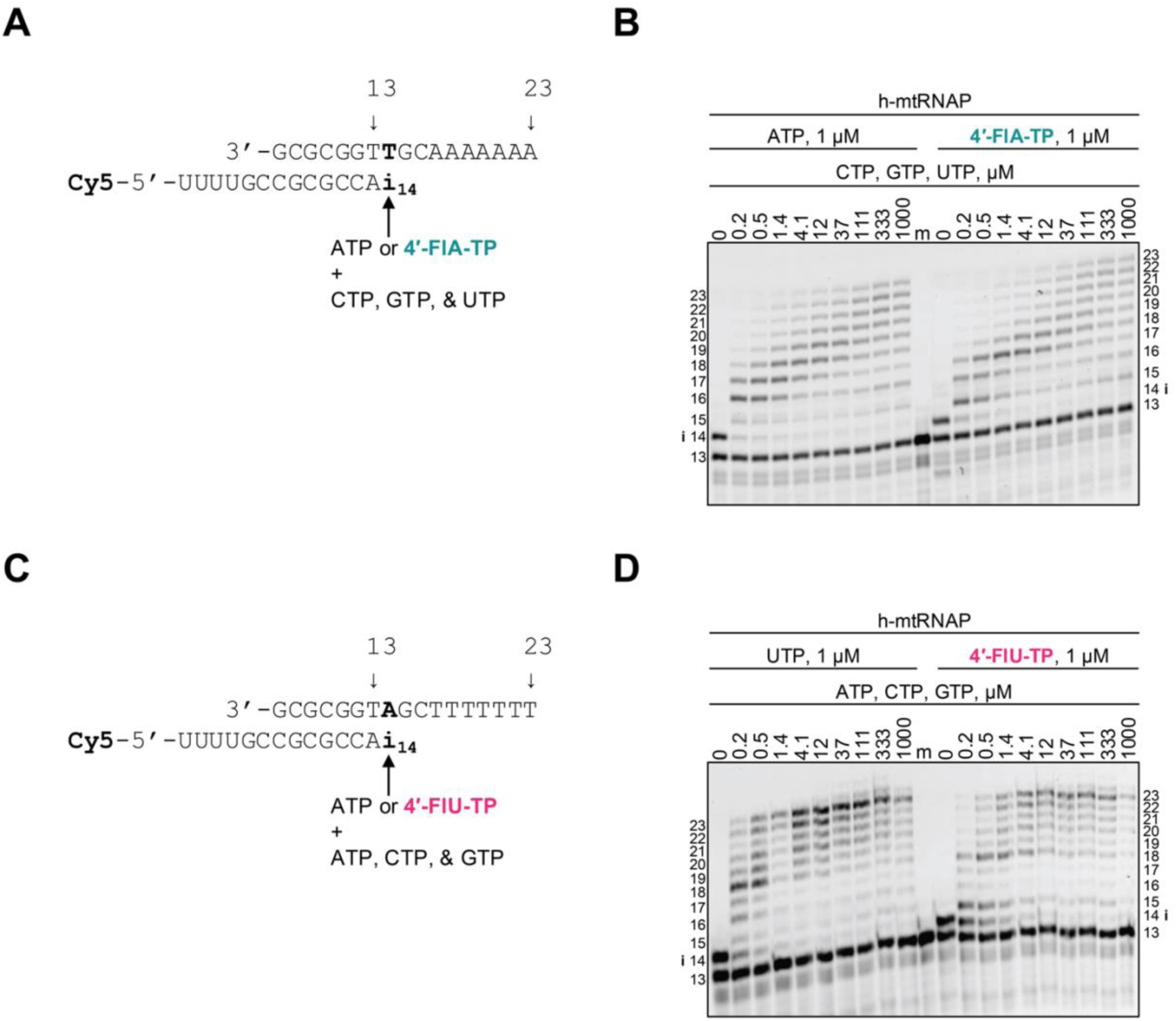
h-mtRNAP DdRp-catalyzed RNA synthesis and inhibition patterns following a single incorporation of ATP or 4′-FlA-TP as a function of nucleotide concentration. (**A**) RNA primer/DNA template supporting a single incorporation of ATP or 4′-FlA-TP at position 14 (“i”). The RNA primer is 5′-Cy5 labelled for signal detection of synthesized RNA products. (**B**) Migration pattern of RNA products resulting from h-mtRNAP DdRp-catalyzed RNA extension following incorporation of ATP or 4′-FlA-TP at position 14 (“i”) at increasing concentrations of CTP, GTP, and UTP. 5′-Cy5-labelled 13-nt primer serves as a size marker (m). (**C**) RNA primer/DNA template supporting a single incorporation of UTP or 4′-FlU-TP at position 14 (“i”). The RNA primer is 5′-Cy5 labelled for signal detection of synthesized RNA products. (**D**) Migration pattern of RNA products resulting from h-mtRNAP DdRp-catalyzed RNA extension following incorporation of UTP or 4′-FlU-TP at position 14 (“i”) at increasing concentrations of ATP, CTP, and GTP. 5′-Cy5-labelled 13-nt primer serves as a size marker (m).

## Discussion

4′-FlU shows a remarkable range of antiviral activity against diverse RNA viruses (25–33). Depending on the pathogen, the *in vitro* potency is often non-inferior or even superior to approved NAs with a broad-spectrum of targets (10–23). These findings warrant further investigation into this class of compounds. Here, we studied the spectrum of antiviral activity of the related 4′-FlA and utilized a comprehensive platform to determine biochemical attributes of the active 4′-FlA-TP form (Figure 6). 4′-FlA demonstrates potent antiviral activity against each of the following viruses: DENV-2, SARS-CoV-2, HRV-16, RSV, LASV, CCHFV, FluA, and FluB. The data also show efficient rates of incorporation of 4′-FlA-TP by the corresponding polymerase complexes. Substrate selectivity measurements suggest that the nucleotide analog can efficiently compete with natural ATP pools. However, efficient incorporation is also seen by host enzymes, which can be indicative of off-target effects (74–77). Both 4′-FlA-TP and 4′-FlU-TP are incorporated by h-mtRNAP, and the deoxy-NTP counterpart of 4′-FlA-TP, 2′-d-4′-FlA-TP, is efficiently incorporated by nonviral DNA polymerases. Together, these data raise concerns regarding target specificity. However, inhibition of host enzymes is not evident, and a favourable SI could be observed in some cases, such as CCHFV.

**Figure 6.**
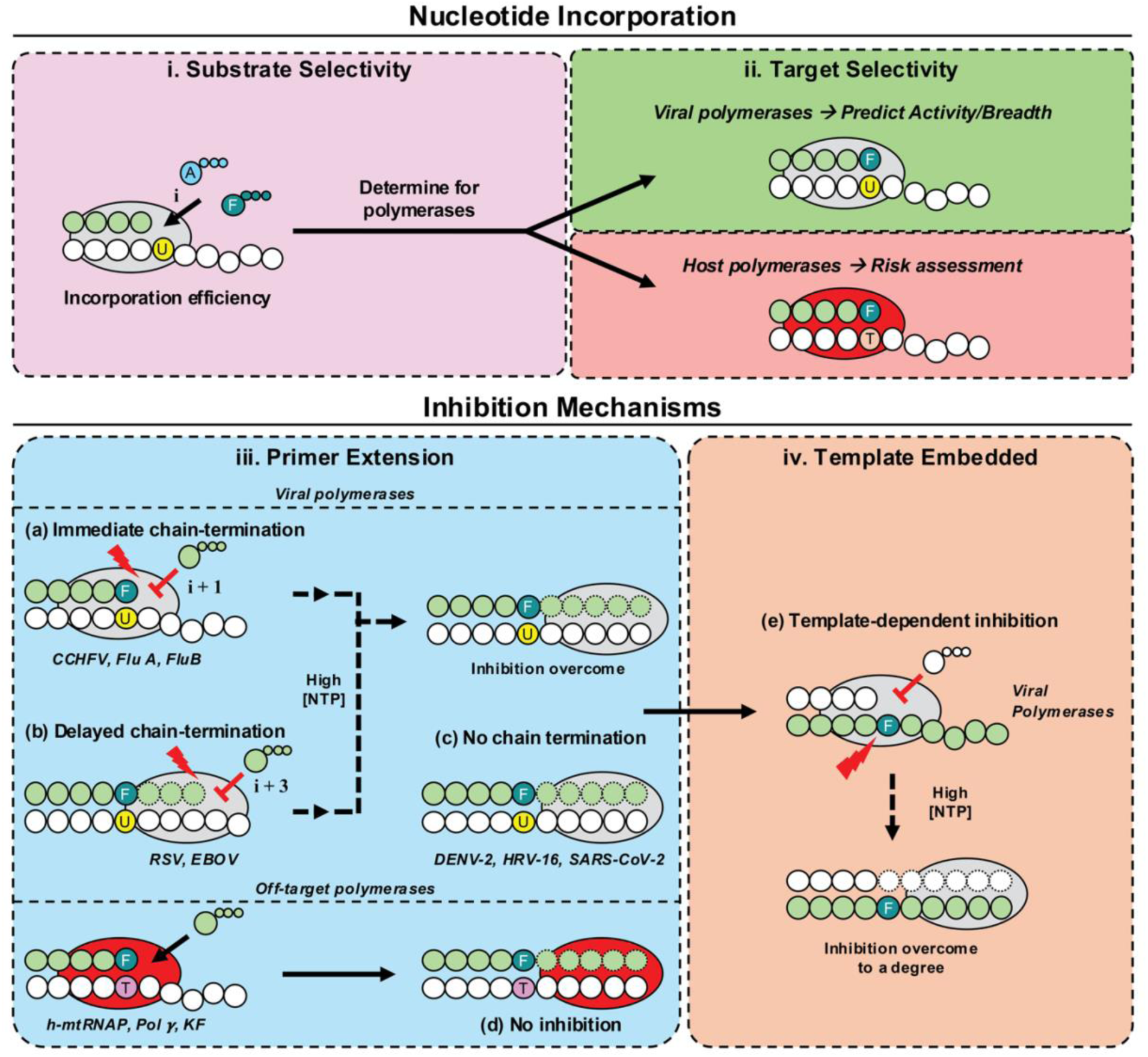
Model of the mechanism of action of 4′-FlA. i) Substrate selectivity. - the investigation of 4′-FlA-TP incorporation efficiency relative to ATP. **ii) Target selectivity** – 4′-FlA-TP is efficiently incorporated by all viral RdRp tested, indicative of potential broad activity. 4′-FlA-TP is also incorporated by nonviral polymerases, indicating potential off-target effects. **iii) Primer extension** – once incorporated, 4′-FlA elicits heterogeneous effects against the viral RdRp, including (a) immediate chain-termination (CCHFV, FluB), (b) delayed chain-termination (RSV, EBOV), and (c) no chain-termination (DENV-2, HRV-16, SARS-CoV-2). When inhibition is observed, increased NTP concentrations appear to be sufficient to overcome inhibition. (d) 4′-FlA shows no inhibitory effect against nonviral polymerases (h-mtRNAP, Pol γ, KF). **iv) Template embedded** – 4′-FlA demonstrates a uniform mechanism against all viral RdRp by inhibiting NTP incorporation opposite the embedded 4′-FlA (e). The priming strand is shown with *green* circles, *white* circles represent residues of the template, ATP is shown in *light blue*, 4′-FlA is shown in *teal*, uridine is shown in *yellow*, thymidine is shown in *pink*, the *red* lightning bolt indicates an inhibitory effect at a given position, the *light grey* oval represents a viral polymerase complex, and the *red* oval represents a nonviral polymerase complex.

The pattern of inhibition within primer-extension reactions appears to depend on the nature of the viral polymerase, which is consistent with reports on 4′-FlU-TP (29,30). The mechanism of action of numerous other 4′-modified nucleotide analogs has been studied, and chain-termination is commonly reported as the dominant mechanism of inhibition (29,44,45,74,89,90). Examples include 4′-azidocytidine (balapiravir) and 2′-deoxy-2′-β-fluoro-4′-azidocytidine (azvudine) that were originally developed for HCV (44,91–94). Lumicitabine, a 2′-fluoro-4′-chloromethyl-cytidine analog, was developed for RSV infection (89,95). GS-7682, an investigational 4′-cyano-modified adenosine analog, is effective against RSV and HRV (96). 4′-ethynyl-2-fluoro-2′-deoxyadenosine (EFdA), developed for the treatment of HIV-1 infection, elicits chain-termination by impairing enzyme translocation (90,97). While 4′-FlU-TP and 4′-FlA-TP show heterogeneous patterns of inhibition of primer extension reactions (29,30), template-dependent inhibition by 4′-FlA provides a unifying mechanism across all viral RdRps included in this study. When embedded in the templating-strand, embedded 4′-FlA-MP inhibits the incorporation of the complementary UTP. Template-dependent inhibition has also been reported for RDV (12,42,45,55). This mechanism needs to be considered for compounds that contain the 3′-hydroxyl group, especially if chain-termination is not absolute. The observation that both 1′-cyano and 4′-fluoro modifications can mediate broad-spectrum antiviral activity through the same mechanism could also be exploited in drug development efforts. The structural differences between these compounds are more sensitive to rates of incorporation, which likely diminishes the antiviral effects of RDV against several segmented negative-sense RNA viruses.

Structural studies are necessary to deepen our understanding of how the 4′-fluoro modification affects substrate use. Our cryo-EM structure provides guidance in this regard and offers insight into the incorporation and subsequent effects of 4′-FlA-TP in the primer strand. It is now of particular interest to determine the structural requirements for template-dependent inhibition. Our structural modelling suggests that this effect may be due to a higher likelihood of steric clashes between the 4′-fluoro modification in the template strand and surrounding residues. However, it is important to note that nucleotide incorporation may accompany conformational changes that our static modelling cannot capture, further highlighting the need for structural studies. It is not evident that such a mechanism may also affect host polymerases. The nature, quantity, and biological relevance of cellular RNA templates with incorporated nucleotide analogs are complex factors that remain to be studied. Modified RNAs generated by host polymerases have numerous functions that may require consideration to assess potential toxicities (98).

In conclusion, the 4′-fluoro modification provides a promising scaffold with a broad-spectrum of antiviral activity that can be ascribed to both efficient incorporation and a unifying template-dependent mechanism of inhibition. Although 4′-FlU and 4′-FlA are viable starting points, substrate utilization by host polymerases raises concerns. The design of derivatives with additional modifications to improve selectivity for viral polymerase targets is warranted. These efforts may be guided by the approach described in this study. Efficacy and safety studies should then accompany the identification of appropriate medical settings that could benefit. Viral infections with high-consequence pathogens are considered in this context.

## Supporting information

Supplementary data are available at NAR online.

## Acknowledgements

The authors would like to thank Emma Woolner for excellent technical and computational assistance, as well as Dr. Jack Moore at the Alberta Proteomics and Mass Spectrometry facility for mass spectrometry analysis. We would also like to thank Dr. Éric Bergeron and Dr. César G. Albariño (Centers for Disease Control and Prevention, CDC) for providing the reverse genetics to generate rCCHFV Zs-Green.

## Author contributions

Conceptualization: SMW, SC, AKC, and MG; Methodology: SMW, AJL, EPT, DK, CJG, KM, RAE, NJ, LMS, KCW, LH, MAB, LR, SP, AG, and MG; Investigation: SMW, AJL, EPT, DK, CJG, IU, MH, ZC, KP, KM, RAE, NJ, LMS, KCW, LH, LR, OAV, SP, AG, AKG, CY; Resources: RAE, NJ, RD, LMS, MG; Data acquisition: SMW, AJL, EPT, IU, MH, ZC, KP, KM, KCW, LH, LR, RAE, NJ, LMS, SP, AKG, CY; Validation: SMW, AJL, KM, KCW, LH, LR, MAB; Formal analysis: SMW, AJL, KM, RAE, NJ, KCW, LH, MAB, LR, OAV, CFB, LMS, SP, and MG; Data curation: SMW, MAB, RAE, NJ, LMS; Visualization: SMW, SP, AG; Writing - original draft: SMW and MG; Writing - review & editing: SMW, AJL, EPT, CJG, MAB, LR, RAE, NJ, LMS, SP, AKG, AG, IAW, and MG; Supervision: LMS, KCW, MAB, RD, IAW, ABW, MG; Project administration: MG; Funding acquisition: CFB, IAW, ABW, AKC, SC, and MG;

## Data availability

The atomic model and cryo-EM density map generated in this study have been deposited in the PDB (accession number: 12SN) and EMDB (accession number: EMD-76733), respectively. The data underlying this article are available in the article itself, its online supplementary material, and the Open Science Framework (OSF) online repository at https://doi.org/10.17605/OSF.IO/A5R4D.

## Supporting Information

**Synthetic chemistry**: 2′-deoxy-4′-FlA-TP (mCQT310 or V2870202)

**Synthetic chemistry**: 4′-FlA-TP (mCOU991 or V2583902)

**Table S1:** Cryo-EM data collection, refinement, and validation statistics for SARS-CoV-2 RdRp complex with dsRNA and 4′-FlA-TP.

| **Data Collection and Processing** |  |
| --- | --- |
| Electron microscope | TFS Glacios 2 |
| Electron detector | Falcon 4i |
| Magnification | 190,000x |
| Voltage (kV) | 200 |
| Electron exposure ($e^-/\text{\AA}^2$) | 45 |
| Defocus range ($\mu\text{m}$) | -0.8 to -1.8 |
| Pixel Size ($\text{\AA}$) | 0.718 |
| Symmetry imposed | C1 |
| Number of movie micrographs | 7,824 |
| Number of particle images in map | 48,482 |
| Map resolution ($\text{\AA}$) | 3.38 |
| FSC threshold | 0.143 |
| Map sharpening $B$ factor ($\text{\AA}^2$) | -78.8 |
| **Model building and validation** |  |
| *Model composition* |  |
| Protein Chains | 4 |
| Protein Residues | 1,176 |
| RNA Chains | 2 |
| RNA Residues | 23 |
| 4'-FIA molecules | 1 |
| *RMSD bonds* |  |
| Bond lengths ($\text{\AA}$) | 0.016 |
| Bond angles ($^\circ$) | 0.838 |
| *Ramachandran plot* |  |
| Favored (%) | 97.84 |
| Outliers (%) | 0.00 |
| *Validation* |  |
| MolProbity score | 1.34 |
| Clashscore | 5.51 |
| Rotamer outliers (%) | 0.58 |
| EMRinger score | 2.50 |
| Map-model correlation coefficient, CC (mask) | 0.78 |

**Table S2:** Selectivity values for mCOU991 (4′-FlA-TP) against viral RdRp.

|  | ATP |  |  | 4'-FlA-TP |  |  |  |
| --- | --- | --- | --- | --- | --- | --- | --- |
| Virus | $V_{\max}^a$<br>(product fraction) | $K_m^b$ (μM) | Efficiency<br>($V_{\max}/K_m$) | $V_{\max}$<br>(product fraction) | $K_m$ (μM) | Efficiency<br>($V_{\max}/K_m$) | Selectivity<sup>c</sup> (fold) |
| **DENV-2** | n=4<sup>d</sup> |  |  |  |  |  |  |
|  | 0.91 | 0.12 | 7.6 | 0.85 | 0.84 | 1.0 | 7.6 |
| $\pm^e$ | 0.01 | 0.01 | | 0.01 | 0.05 | | |
| **HRV-16** | n=3 |  |  |  |  |  |  |
|  | 0.79 | 1.5 | 0.53 | 0.75 | 0.70 | 1.1 | 0.48 |
| ± | 0.02 | 0.2 | | 0.02 | 0.07 | | |
| **SARS-COV-2** | n=3 |  |  |  |  |  |  |
|  | 0.69 | 0.070 | 9.9 | 0.54 | 0.053 | 10 | 0.99 |
| ± | 0.01 | 0.003 | | 0.01 | 0.002 | | |
| **RSV** | n=3 |  |  |  |  |  |  |
|  | 0.72 | 0.69 | 1.0 | 0.49 | 3.5 | 0.14 | 7.1 |
| ± | 0.02 | 0.11 | | 0.02 | 0.5 | | |
| **EBOV** | n=3 |  |  |  |  |  |  |
|  | 0.63 | 1.9 | 0.33 | 0.52 | 6.6 | 0.079 | 4.2 |
| ± | 0.04 | 0.4 | | 0.03 | 1.4 | | |
| **CCHFV** | n=3 |  |  |  |  |  |  |
|  | 0.62 | 0.26 | 2.4 | 0.57 | 0.33 | 1.7 | 1.4 |
| ± | 0.03 | 0.06 | | 0.04 | 0.09 | | |
| **FluB** | n=4 |  |  |  |  |  |  |
|  | 0.80 | 0.19 | 4.2 | 0.80 | 0.26 | 3.1 | 1.4 |
| ± | 0.02 | 0.02 | | 0.02 | 0.04 | | |

<sup>a</sup> $V_{\max}$ is a Michaelis–Menten parameter reflecting the maximal velocity of nucleotide incorporation.
<sup>b</sup> $K_m$ is a Michaelis–Menten parameter reflecting the concentration of the nucleotide substrate at which the velocity of nucleotide incorporation is half of $V_{\max}$ .
<sup>c</sup> Selectivity of a viral RNA polymerase for a nucleotide substrate analog is calculated as the ratio of the $V_{\max}/K_m$ values for NTP and NTP analog, respectively.
<sup>d</sup> All reported values have been calculated based on an 8–data point experiment repeated the indicated number of times (n).
<sup>e</sup> Standard error associated with the fit.

**Table S3:**
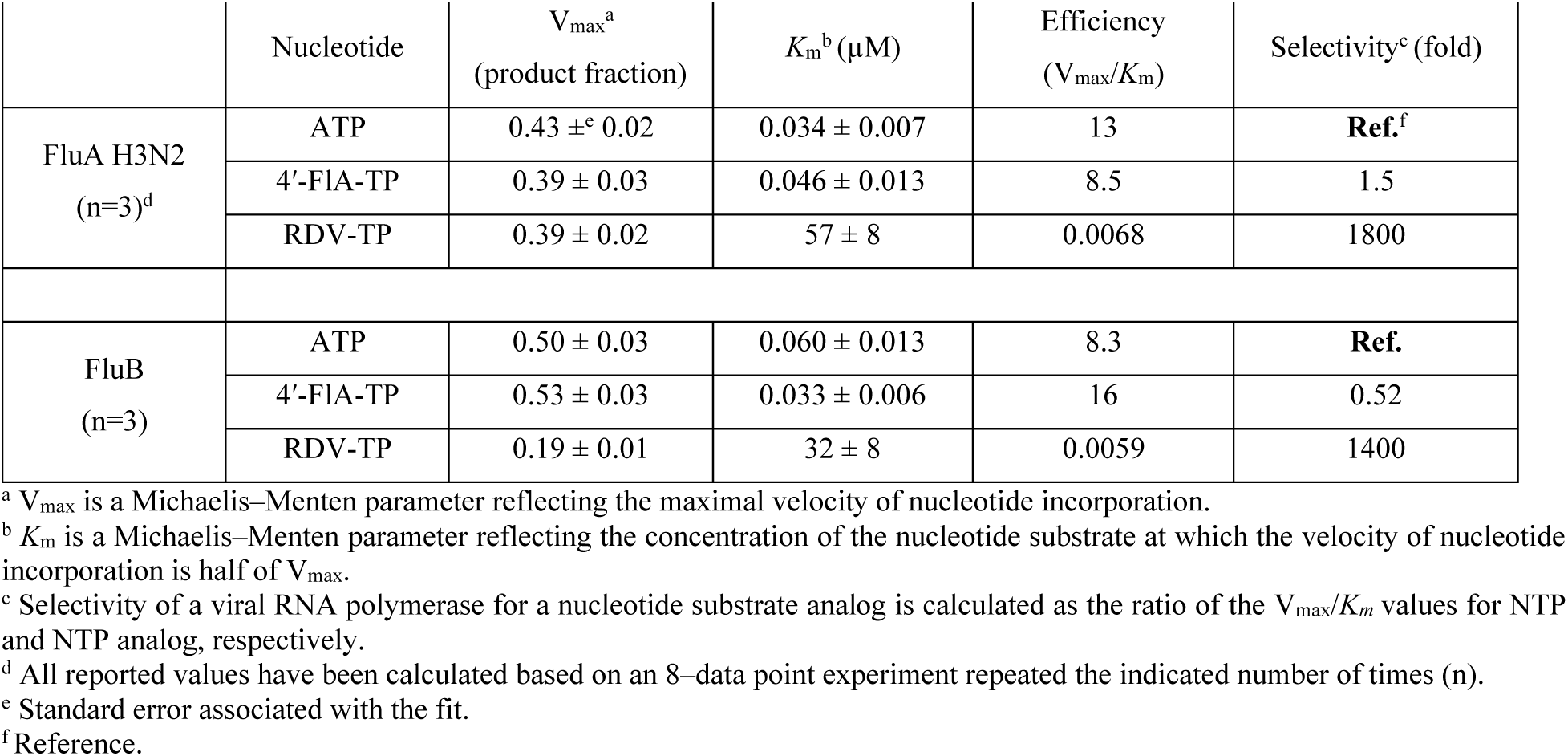
Selectivity values for RDV-TP and 4′-FlA-TP against influenza RdRp using 13-nt capped primer system.

**Table S4:** Inhibitory effect of the incorporated analog-monophosphate on the incorporation of the subsequent nucleotide.

| Enzyme | Primer 3'-end (base) | Substrate | V<sub>max</sub> (product fraction) | K<sub>m</sub><sup>d</sup> (μM) | V<sub>max</sub>/K<sub>m</sub> | Inhibition<sup>e</sup> (fold) |
| --- | --- | --- | --- | --- | --- | --- |
| **DENV-2**<br>(n=3)<sup>f</sup> | AMP<sup>a</sup> (1 μM)<sup>b</sup> | UTP | 0.84 ±<sup>g</sup> 0.03 | 0.060 ± 0.008 | 14 | Ref.<sup>h</sup> |
|  | 4'-FIA-MP (10 μM) |  | 0.84 ± 0.03 | 0.23 ± 0.02 | 3.7 | 3.8 |
| **HRV-16**<br>(n=3) | AMP (10 μM) | UTP | 0.60 ± 0.01 | 0.044 ± 0.004 | 14 | Ref. |
|  | 4'-FIA-MP (10 μM) |  | 0.62 ± 0.02 | 0.13 ± 0.01 | 4.8 | 2.9 |
| **SARS-CoV-2**<br>(n=4) | AMP (1 μM) | UTP | 0.70 ± 0.01 | 0.013 ± 0.001 | 54 | Ref. |
|  | 4'-FIA-MP (1 μM) |  | 0.68 ± 0.02 | 0.068 ± 0.008 | 10 | 5.4 |
| **RSV**<br>(n=3) | AMP (100 μM) | UTP | 0.61 ± 0.03 | 0.69 ± 0.10 | 0.88 | Ref. |
|  | 4'-FIA-MP (100 μM) |  | 0.58 ± 0.02 | 8.8 ± 1.0 | 0.066 | 13 |
| **EBOV**<br>(n=4) | AMP (100 μM) | UTP | 0.31 ± 0.02 | 4.0 ± 1.0 | 0.078 | Ref. |
|  | 4'-FIA-MP (100 μM) |  | 0.29 ± 0.01 | 30 ± 5.2 | 0.0097 | 8.0 |
| **CCHFV**<br>(n=3) | AMP (10 μM) | UTP | 0.63 ± 0.02 | 0.20 ± 0.04 | 3.2 | Ref. |
|  | 4'-FIA-MP (10 μM) |  | 0.43 ± 0.02 | 4.4 ± 0.6 | 0.098 | 33 |
| **FluB**<br>(n=3) | AMP (10 μM) | UTP | 0.69 ± 0.02 | 0.012 ± 0.001 | 58 | Ref. |
|  | 4'-FIA-MP (10 μM) |  | 0.42 ± 0.01 | 5.1 ± 0.5 | 0.082 | 710 |
| **h-mtRNAP**<br>(n=3) | AMP (1 μM) | CTP | 0.36 ± 0.01 | 0.033 ± 0.002 | 11 | Ref. |
|  | 4'-FIA-MP (10 μM) |  | 0.32 ± 0.01 | 0.053 ± 0.004 | 6.0 | 1.8<sup>i</sup> |

<sup>a</sup> Denotes that the 3'-end of the primer is terminated by the monophosphate form of the respective nucleotide.
<sup>b</sup> Concentration of the respective nucleotide that terminates the 3'-end of the primer.
<sup>c</sup> $V_{max}$ is a Michaelis–Menten parameter reflecting the maximal velocity of nucleotide incorporation.
<sup>d</sup> $K_m$ is a Michaelis–Menten parameter reflecting the concentration of the nucleotide substrate at which the velocity of nucleotide incorporation is half of $V_{max}$ .
<sup>e</sup> Inhibition of subsequent nucleotide incorporation is calculated as the ratio of $V_{max}/K_m$ values for UTP determined on primers terminated with AMP or 4'-FIA.
<sup>f</sup> All reported values have been calculated based on an 8–data point experiment repeated the indicated number of times (n).
<sup>g</sup> Standard error associated with the fit.
<sup>h</sup> Reference.
<sup>i</sup> Inhibition of subsequent nucleotide incorporation is calculated as the ratio of $V_{max}/K_m$ values for CTP determined on primers terminated with AMP or 4'-FIA.

**Table S5:** Selectivity values for nucleotide analogs against off-target polymerases.

| | Nucleotide | $V_{\max}^a$<br>(product fraction) | $K_m^b$ ($\mu\text{M}$) | Efficiency<br>($V_{\max}/K_m$) | Selectivity<sup>c</sup> (fold) |
| --- | --- | --- | --- | --- | --- |
| h-mtRNAP | ATP | $0.39^d \pm 0.01$ | $0.015 \pm 0.001$ | 26 | **Ref.**<sup>f</sup> |
| | RDV-TP | $0.14 \pm 0.010$ | $12 \pm 4$ | 0.012 | 2,200 |
| | 4'-FIA-TP | $0.43 \pm 0.004$ | $0.14 \pm 0.01$ | 3.1 | 8.4 |
| | UTP | $0.38 \pm 0.01$ | $0.029 \pm 0.002$ | 13 | **Ref.** |
|  | SOF-TP |  |  |  | B.L.Q.<sup>g</sup> |
| | 4'-FIU-TP | $0.48 \pm 0.01$ | $0.15 \pm 0.02$ | 3.2 | 4.1 |
| | CTP | $0.46 \pm 0.01$ | $0.039 \pm 0.003$ | 12 | **Ref.** |
| | NHC-TP | $0.59 \pm 0.03$ | $1.1 \pm 0.3$ | 0.54 | 22 |
| Pol $\gamma$ | dATP | $0.37 \pm 0.03$ | $0.012 \pm 0.003$ | 31 | **Ref.** |
| | ddATP | $0.28 \pm 0.02$ | $0.0060 \pm 0.0012$ | 47 | 0.66 |
| | TFV-DP | $0.29 \pm 0.01$ | $2.0 \pm 0.3$ | 0.15 | 210 |
| | 2'-d-4'-FIA-TP | $0.36 \pm 0.02$ | $0.010 \pm 0.003$ | 33 | 0.94 |
| Klenow<br>-exonuclease | dATP | $0.38 \pm 0.01$ | $0.025 \pm 0.002$ | 15 | **Ref.** |
| | ddATP | $0.22 \pm 0.02$ | $11 \pm 4$ | 0.020 | 750 |
|  | TFV-DP |  |  |  | B.L.Q. |
| | 2'-d-4'-FIA-TP | $0.29 \pm 0.02$ | $0.094 \pm 0.020$ | 3.1 | 4.8 |

<sup>a</sup> $V_{\max}$ is a Michaelis–Menten parameter reflecting the maximal velocity of nucleotide incorporation.
<sup>b</sup> $K_m$ is a Michaelis–Menten parameter reflecting the concentration of the nucleotide substrate at which the velocity of nucleotide incorporation is half of $V_{\max}$ .
<sup>c</sup> Selectivity of a viral RNA polymerase for a nucleotide substrate analog is calculated as the ratio of the $V_{\max}/K_m$ values for NTP and NTP analog, respectively.
<sup>d</sup> All reported values have been calculated based on an 8–data point experiment repeated at least 3 times (n=3).
<sup>e</sup> Standard error associated with the fit.
<sup>f</sup> Reference.
<sup>g</sup> Below the level of quantification, B.L.Q.

**Figure S1.**
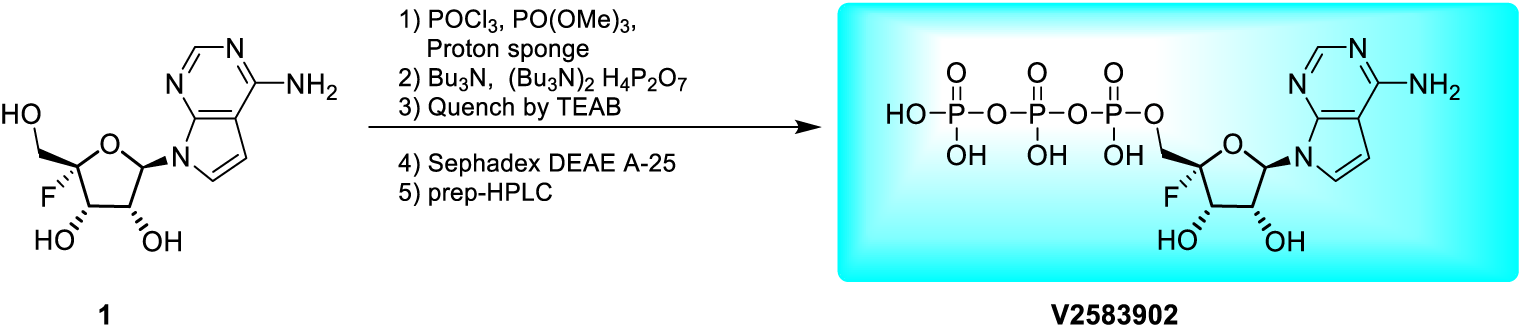
Chemical synthesis scheme of 4′-fluoroadenosine triphosphate (4′-FlA-TP).

**Figure S2.**
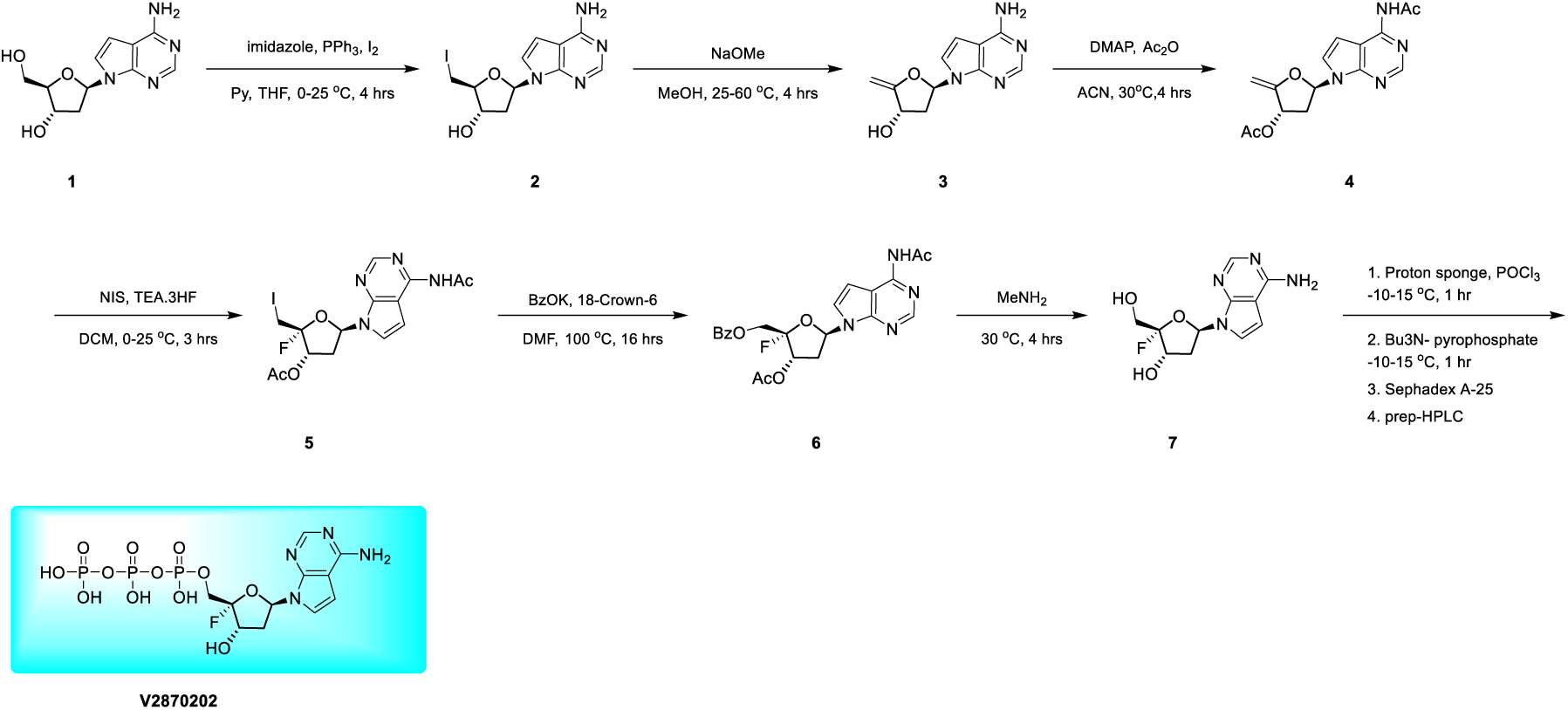
Chemical synthesis scheme of 2′-deoxy-4′-fluoroadenosine triphosphate (2′-d-4′-FlA-TP).

**Figure S3.**
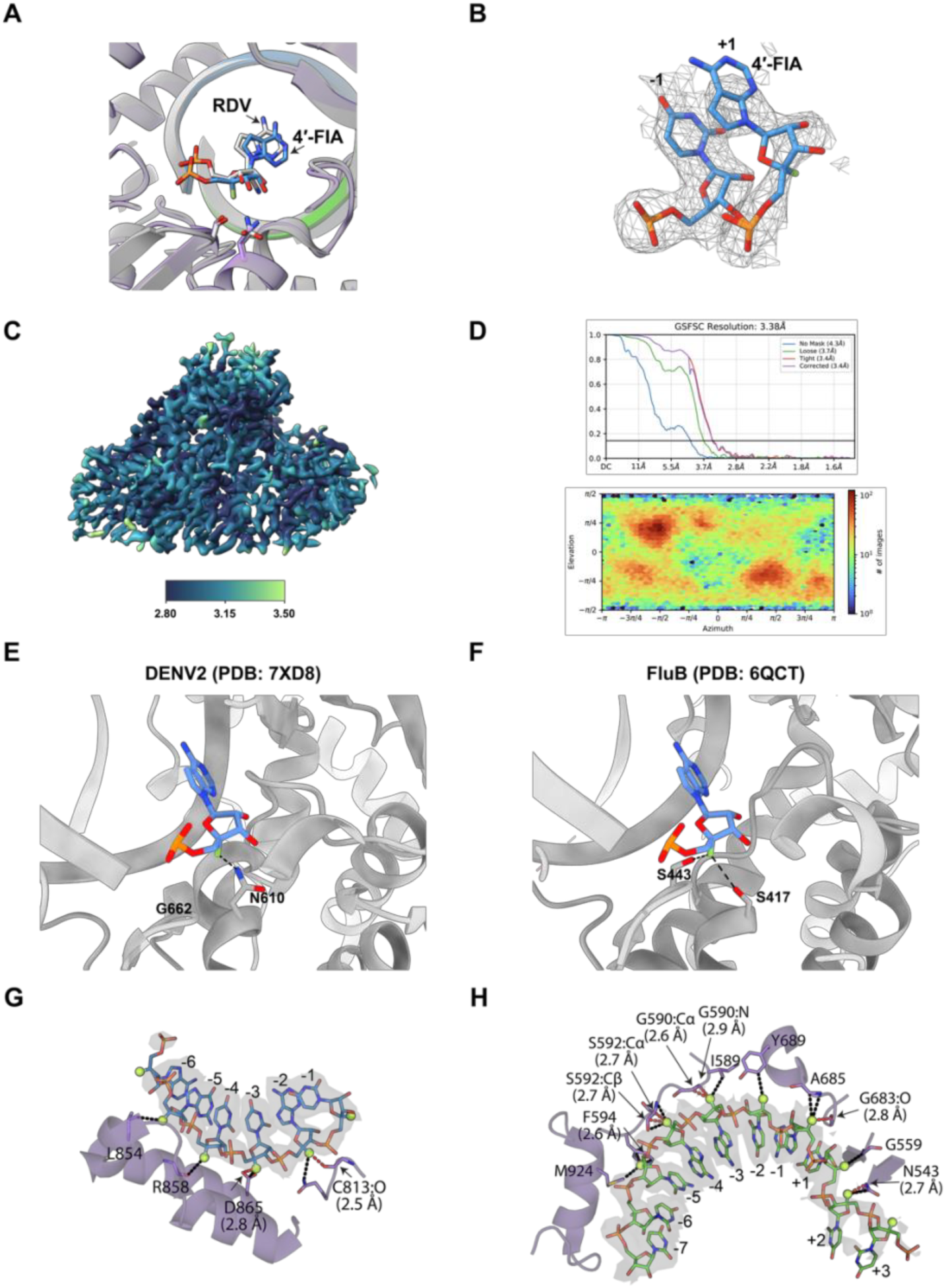
Structural basis for 4′-FlA incorporation by the SARS-CoV-2 RdRp and cryo-EM data quality. (**A**) Structural superposition of the 4′-FlA-bound RdRp complex (purple) with the remdesivir (RDV)-bound RdRp structure (white). Both 4′-FlA and RDV are shown as stick models and are indicated by arrows. (**B**) Cryo-EM density map of 4′-FlA molecule incorporated at the +1 position and the adjacent -1 nucleotide of the primer strand. The map is shown as a mesh, and nucleotides are displayed as stick models. (**C**) Local resolution map of the cryo-EM density. (**D**) Gold-standard Fourier shell correlation (GSFSC) curve (top). Angular distribution plot of particles used for the final reconstruction (bottom). (**E**) Structural superposition of 4′-FlA resolved in this study with DENV-2 RdRp elongation complex. (**F**) Structural superposition of 4′-FlA resolved in this study with FluB RdRp elongation complex. (**G** and **H**) Modelled interactions between the 4′-fluoro modification and nsp12 residues in the primer strand (**G**) and template strand (**H**), based on PDB: 7BV2 (1). RNA nucleotides are shown as green (template) or cyan (primer) sticks, with the 4′-fluorine atom displayed as a green sphere. All contacts within 3.5 Å of the fluorine atom are indicated by dashed lines, with contacts shorter than 3.0 Å shown as red dashed lines with distances labelled. For backbone interactions, the relevant atoms are explicitly displayed. Interacting nsp12 residues are labelled.

**Figure S4.**
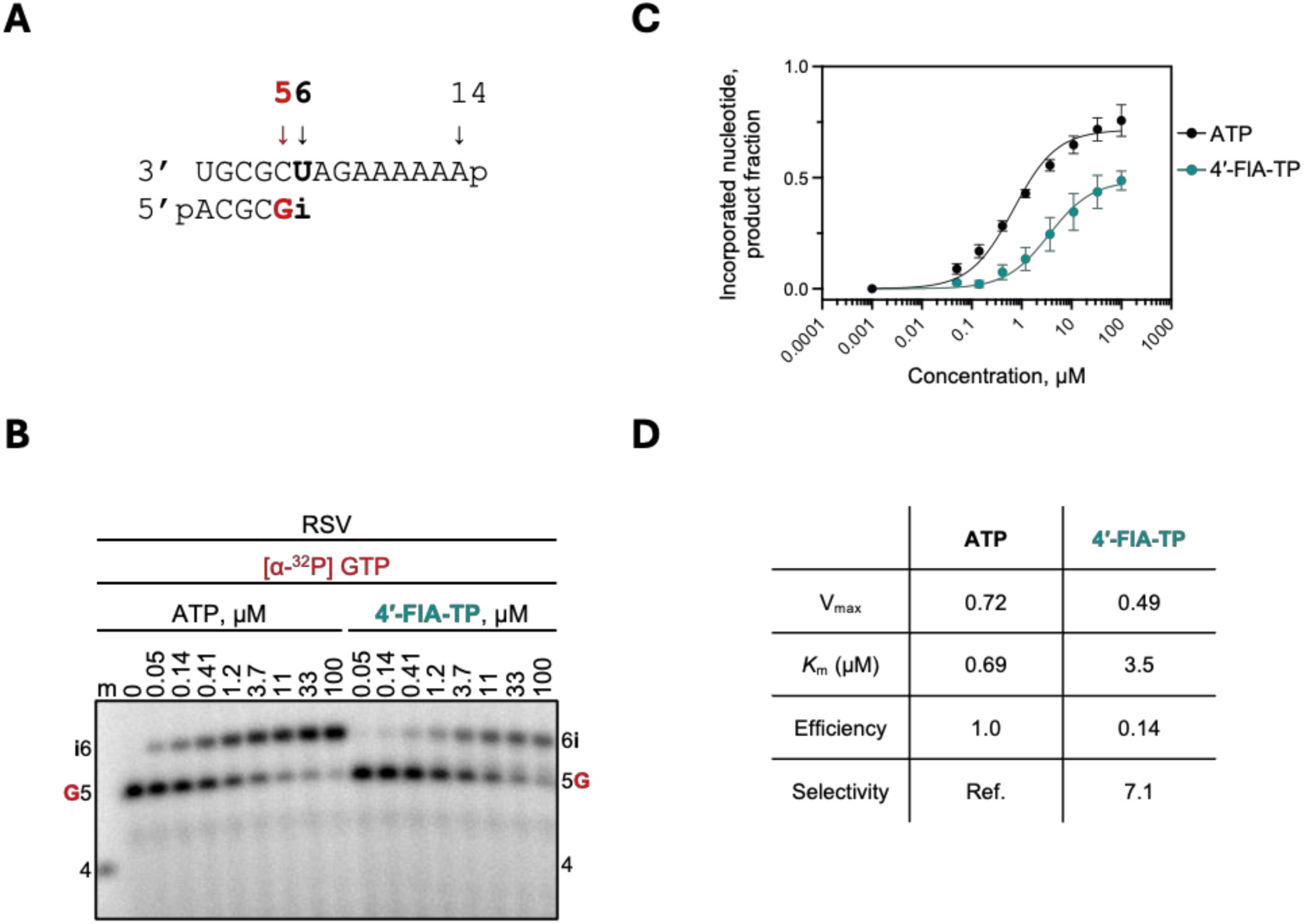
Selective incorporation of 4′-FlA-TP by RSV RdRp. (**A**) RNA primer/template substrate used in the RNA synthesis assays to evaluate 4′-FlA-TP as a substrate for incorporation as an A-analog incorporation is shown above the gel. G indicates incorporation of the radiolabelled nucleotide opposite template position 5. Position (“i”) allows incorporation of ATP or 4′-FlA-TP. (**B**) NTP incorporation was monitored with purified RSV RdRp complex in the presence of [α-^32^P]-GTP, RNA primer/template, MgCl_2_, and increasing concentrations of ATP or 4′-FlA-TP. Lane m illustrates the migration pattern of the radiolabelled 4-nucleotide-long primer. (**C**) Graphic representation of the data for incorporation of ATP and 4′-FlA-TP. (**D**) Kinetic parameters and efficiency of ATP and 4′-FlA-TP incorporation by RSV RdRp. Efficiency is determined as the quotient of V_max_ over *K*_m_. Selectivity is the quotient of ATP efficiency over ATP-analog efficiency.

**Figure S5.**
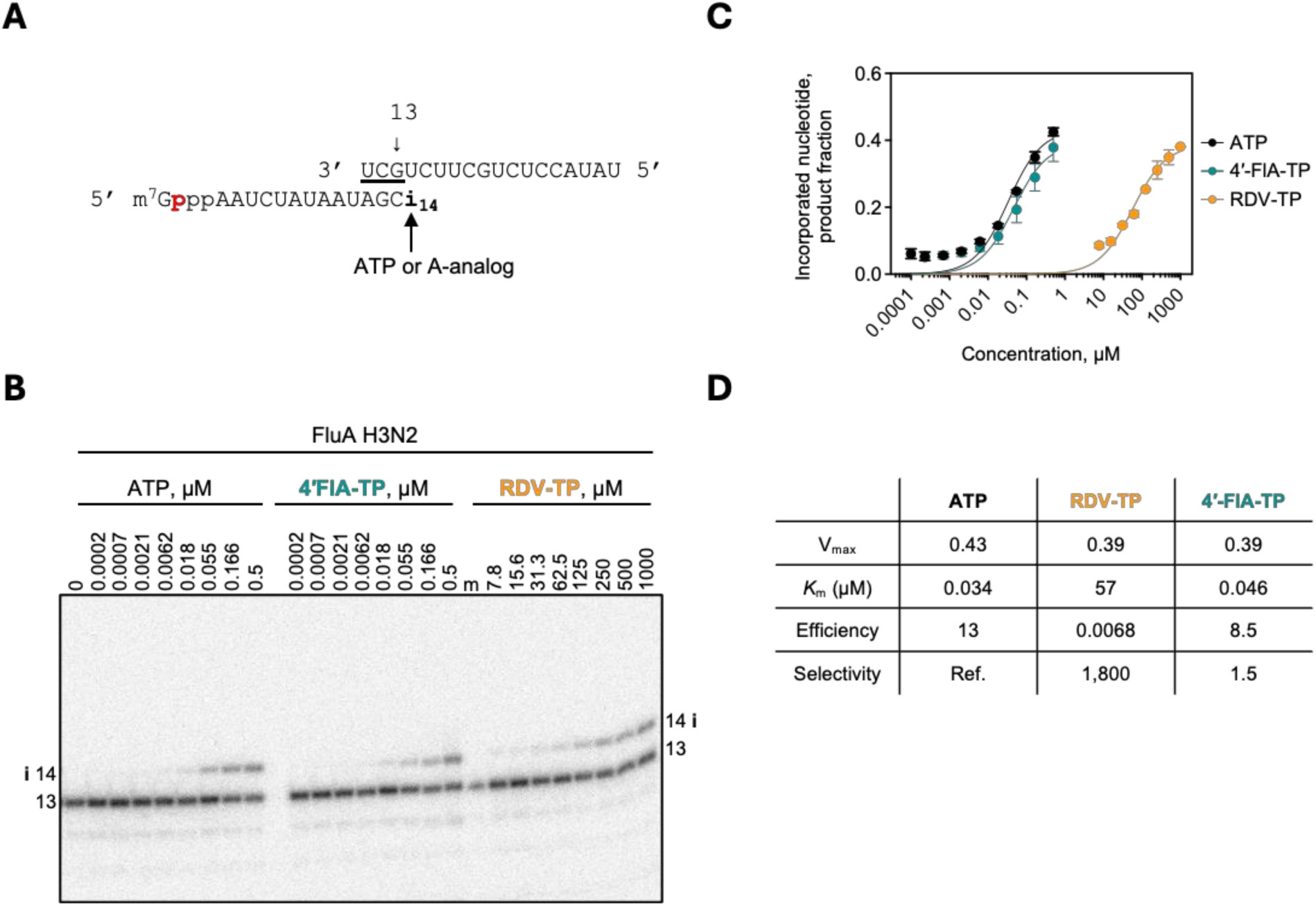
Selective incorporation of 4′-FlA-TP by FluA RdRp in the capped primer extension assay. (**A**) 3′ vRNA template and 13-nt capped radiolabeled RNA primer used to evaluate A-analog incorporation at position 14 (“i”). (**B**) Nucleotide incorporation was monitored with purified FluA RdRp complex in the presence of the radiolabelled RNA primer, vRNA template, MgCl_2,_ BXA (to prevent endonuclease cleavage of the capped primer), and increasing concentrations of the indicated nucleotide (ATP, 4′-FlA-TP, or RDV-TP). Lane m indicates the 13-nt labelled primer used as a marker. (**C**) Graphic representation of ATP and 4′-FlA-TP incorporation. (**D**) Kinetic parameters determined from quantification of product signal, which is ATP or 4′-FlA-TP incorporation by FluA RdRp. Efficiency is determined as the quotient of V_max_ over *K*_m_. Selectivity is the quotient of ATP efficiency over ATP-analog efficiency.

**Figure S6.**
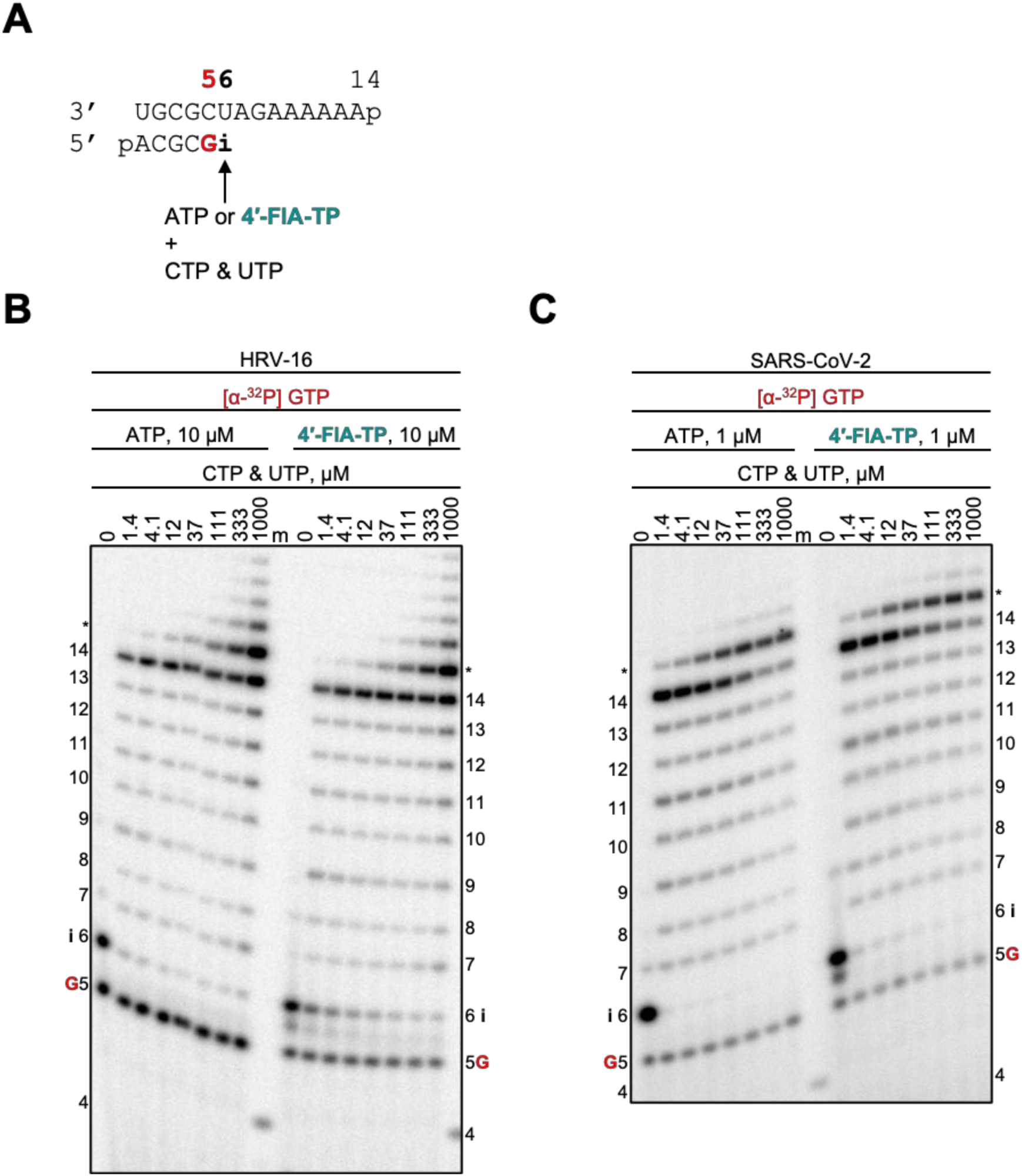
HRV-16 and SARS-CoV-2 RdRp-catalyzed RNA synthesis and inhibition patterns following a single incorporation of ATP or 4′-FlA-TP as a function of nucleotide concentration. (**A**) RNA primer/template supporting a single incorporation of ATP or 4′-FlA-TP at position 6 (“i”). G indicates incorporation of [α-^32^P]-GTP at position 5. Migration pattern of RNA products resulting from (**B**) HRV-16 RdRp- and (**C**) SARS-CoV-2 RdRp-catalyzed RNA extension following incorporation of ATP or 4′-FlA-TP at position 6 (“i”) at increasing CTP and UTP concentrations. A 5′-^32^P-labelled 4-nt primer (4) serves as a size marker (m). RNA products formed at and beyond the asterisk may be a result of sequence-dependent slippage events or RdRp nucleotide transferase activity. HRV, human rhinovirus; SARS-CoV-2, severe acute respiratory syndrome coronavirus 2; RdRp, RNA-dependent RNA polymerase.

**Figure S7.**
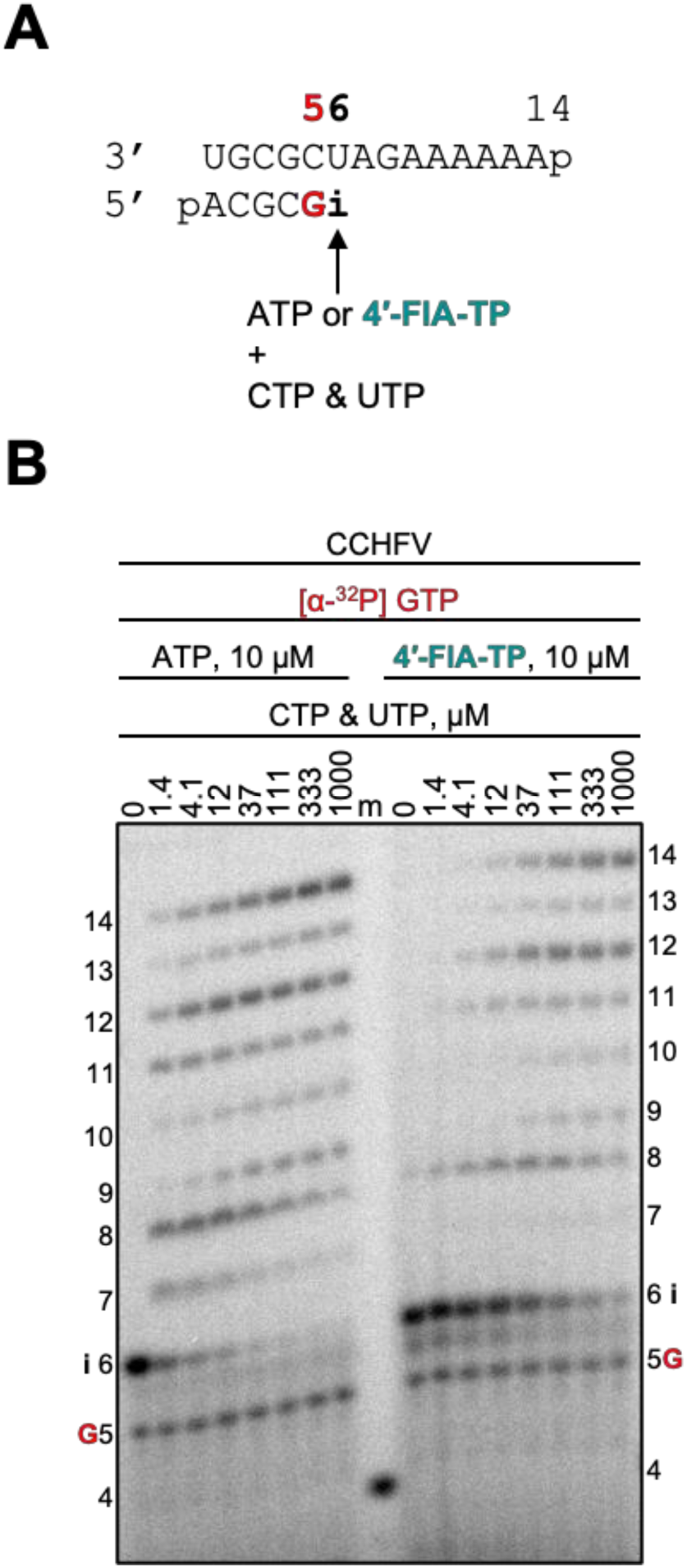
CCHFV RdRp-catalyzed RNA synthesis and inhibition patterns following a single incorporation of ATP or 4′-FlA-TP as a function of nucleotide concentration. (**A**) RNA primer/template supporting a single incorporation of ATP or 4′-FlA-TP at position 6 (“i”). G indicates incorporation of [α-^32^P]-GTP at position 5. Migration pattern of RNA products resulting from (**B**) CCHFV RdRp-catalyzed RNA extension following incorporation of ATP or 4′-FlA-TP at position 6 (“i”) at increasing CTP and UTP concentrations. A 5′-^32^P-labelled 4-nt primer (4) serves as a size marker (m). CCHFV, Crimean-Congo hemorrhagic fever virus; RdRp, RNA-dependent RNA polymerase.

**Figure S8.**
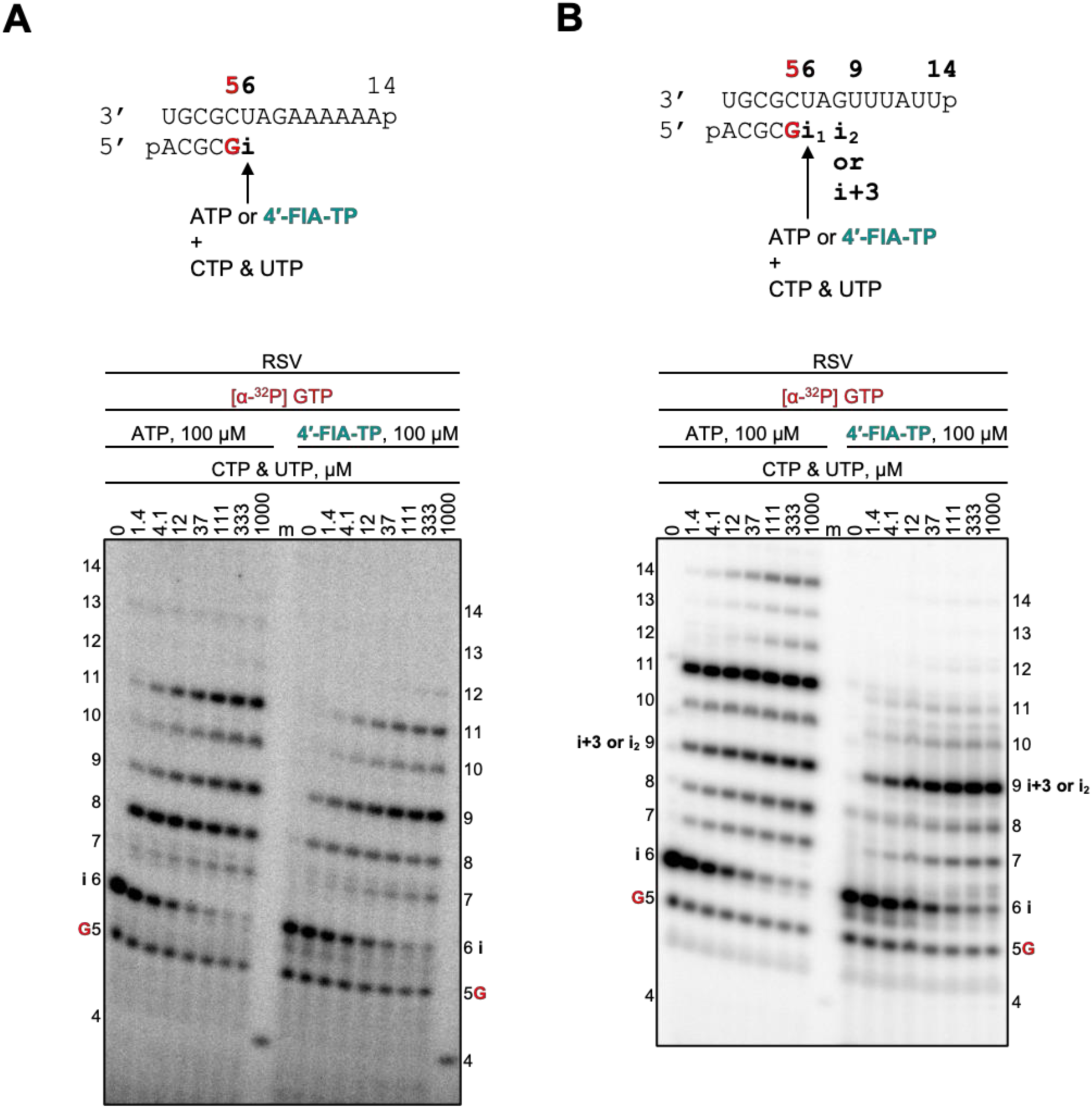
RSV RdRp-catalyzed RNA synthesis and inhibition patterns following a single or multiple incorporations of ATP or 4′-FlA-TP as a function of nucleotide concentration. (**A**) RNA primer/template supporting a single incorporation of ATP or 4′-FlA-TP at position 6 (“i”). G indicates incorporation of [α-32P]-GTP at position 5. Migration pattern of RNA products resulting from RSV RdRp-catalyzed RNA extension following a single incorporation of ATP or 4′-FlA-TP at increasing CTP and UTP concentrations. Investigation of a single incorporated 4′-FlA-TP is difficult, given that full-template length product is not generated even under the natural scenario with ATP, which has been observed previously (2). A 5′-^32^P-labelled 4-nt primer (4) serves as a size marker (m). (**B**) RNA primer/template supporting a first and second incorporation of ATP or 4′-FlA-TP at position 6 (“i”) and position 9 (“i_2_”). G indicates incorporation of [α-^32^P]-GTP at position 5. Migration pattern of RNA products resulting from RSV RdRp-catalyzed RNA extension following multiple incorporations of ATP or 4′-FlA-TP at increasing CTP and UTP concentrations. A 5′-^32^P-labelled 4-nt primer (4) serves as a size marker (m). Multiple 4′-FlA-TP incorporations do not appear to allow production of full-template length product. RSV, Respiratory syncytial virus; RdRp, RNA-dependent RNA polymerase.

**Figure S9.**
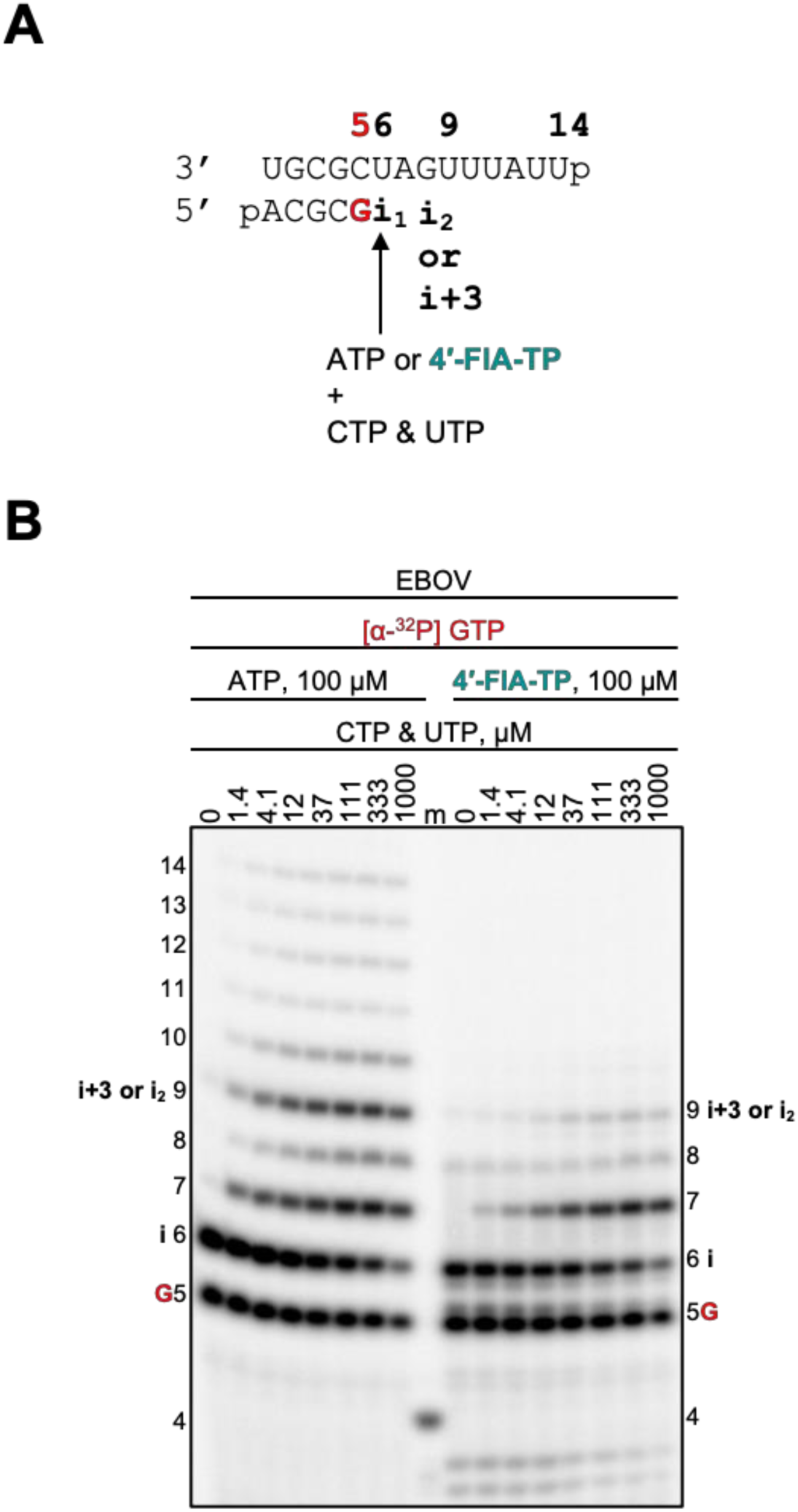
EBOV RdRp-catalyzed RNA synthesis and inhibition patterns following multiple incorporations of ATP or 4′-FlA-TP as a function of nucleotide concentration. (**A**) RNA primer/template supporting a first and second incorporation of ATP or 4′-FlA-TP at position 6 (“i”) and position 9 (“i_2_”). G indicates incorporation of [α-^32^P]-GTP at position 5. (**B**) Migration pattern of RNA products resulting from EBOV RdRp-catalyzed RNA extension following multiple incorporations of ATP or 4′-FlA-TP at increasing CTP and UTP concentrations. A 5′-^32^P-labelled 4-nt primer (4) serves as a size marker (m). Multiple 4′-FlA-TP incorporations do not appear to allow production of full-template length product. EBOV, Ebola virus; RdRp, RNA-dependent RNA polymerase.

**Figure S10.**
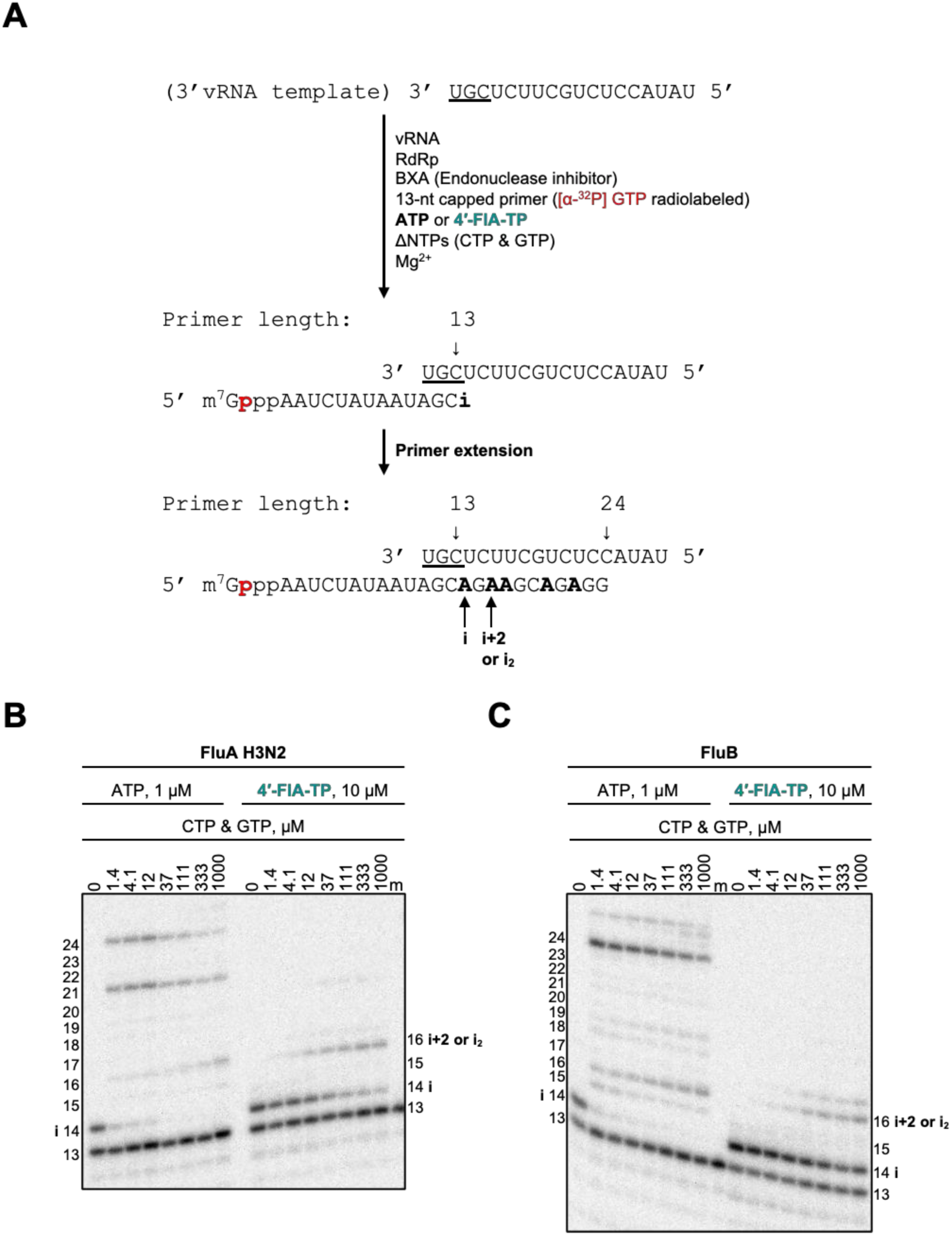
FluA and FluB RdRp-catalyzed RNA synthesis and inhibition patterns following multiple incorporations of ATP or 4′-FlA-TP as a function of nucleotide concentration. (**A**) A 13-nt capped radiolabeled RNA primer and vRNA template required for FluA RdRp activity. Reactions consist of vRNA template, capped primer, viral RdRp, BXA (an influenza endonuclease inhibitor), ATP or 4′-FlA-TP, increasing concentrations of CTP and GTP, and Mg^2+^ (which starts the reaction). The primer-template supports the incorporation of ATP or 4′-FlA-TP at position 14 (“i”). Additional incorporations of ATP or 4′-FlA-TP can occur as indicated. Migration pattern of RNA products resulting from (**B**) FluA RdRp- and (**C**) FluB RdRp-catalyzed RNA extension following incorporation of ATP or 4′-FlA-TP at position 14 (“i”) at increasing CTP and GTP concentrations. 5′-^32^P-labelled 13-nt primer (13) serves as a size marker (m). Investigation of 4′-FlA-TP using this template is difficult given a single incorporation event is not possible. A second site of 4′-FlA-TP incorporation does not appear to be overcome. FluA, Influenza A virus; FluB, Influenza B virus; RdRp, RNA-dependent RNA polymerase.

**Figure S11.**
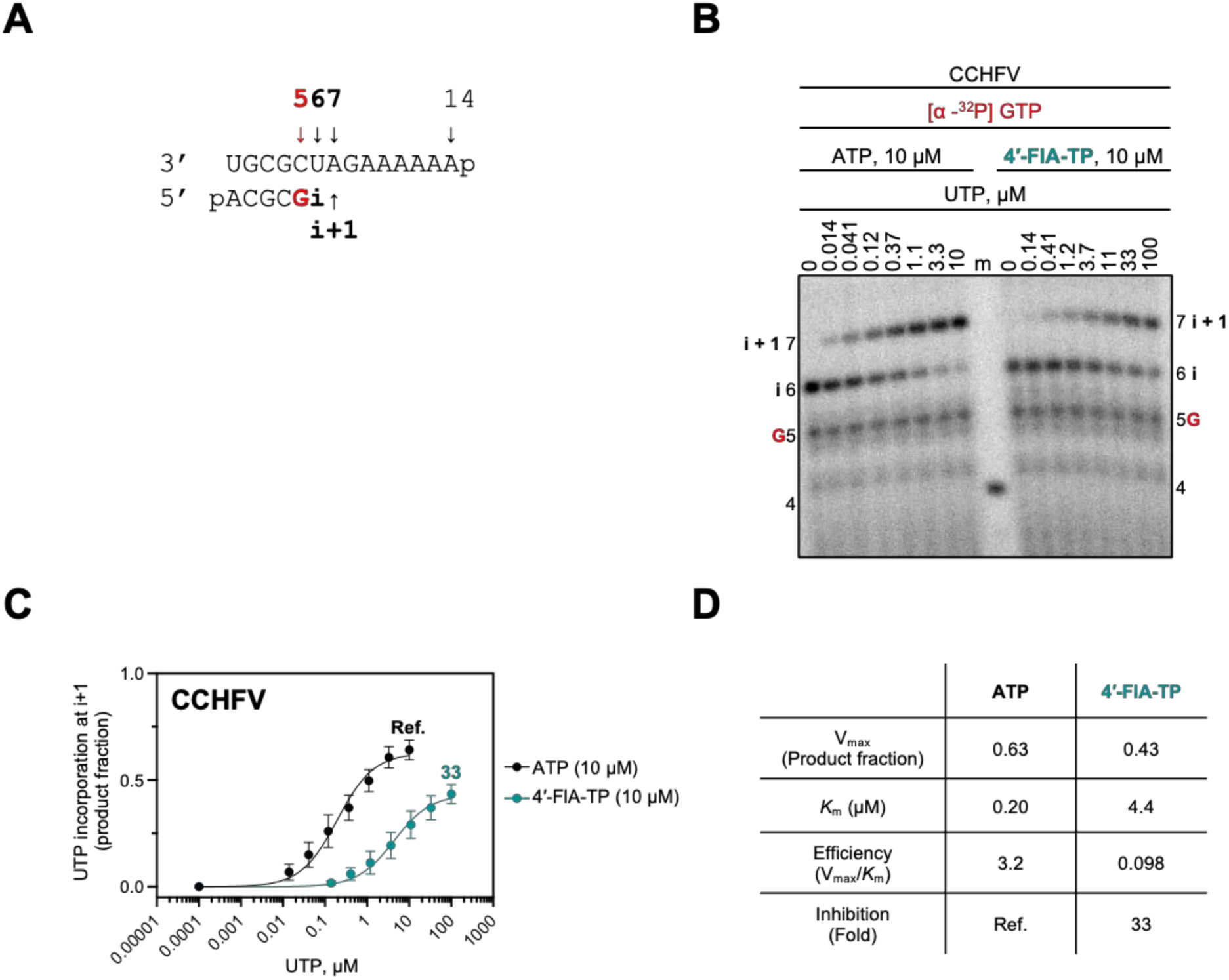
Inhibitory effect of RNA extension by CCHFV RdRp following 4′-FlA-TP incorporation. (**A**) RNA primer/template supporting a single incorporation of ATP or 4′-FlA-TP at position 6 (“i”) and a subsequent incorporation of UTP at position 7 (“i+1”). G indicates incorporation of [α-^32^P]-GTP at position 5. (**B**) CCHFV RdRp-catalyzed RNA extension to position “i+1” following incorporation of ATP or 4′-FlA-TP at position 6 (“i”) at increasing UTP concentrations. ATP or 4′-FlA-TP were kept at fixed concentrations. RNA products were resolved by denaturing PAGE. A 5′-^32^P-labelled 4-nt primer (4) serves as a size marker (m). (**C**) Graphic representation of the data for incorporation of UTP by CCHFV RdRp following the incorporation of ATP and 4′-FlA-TP. (**D**) Kinetic parameters and efficiency of UTP incorporation by CCHFV RdRp following the incorporation of ATP and 4′-FlA-TP. Efficiency is determined as the quotient of V_max_ over *K*_m_. Inhibition is the quotient of UTP incorporation efficiency following ATP incorporation, divided by the efficiency of UTP incorporation following 4′-FlA-TP incorporation. CCHFV, Crimean-Congo hemorrhagic fever virus; RdRp, RNA-dependent RNA polymerase.

**Figure S12.**
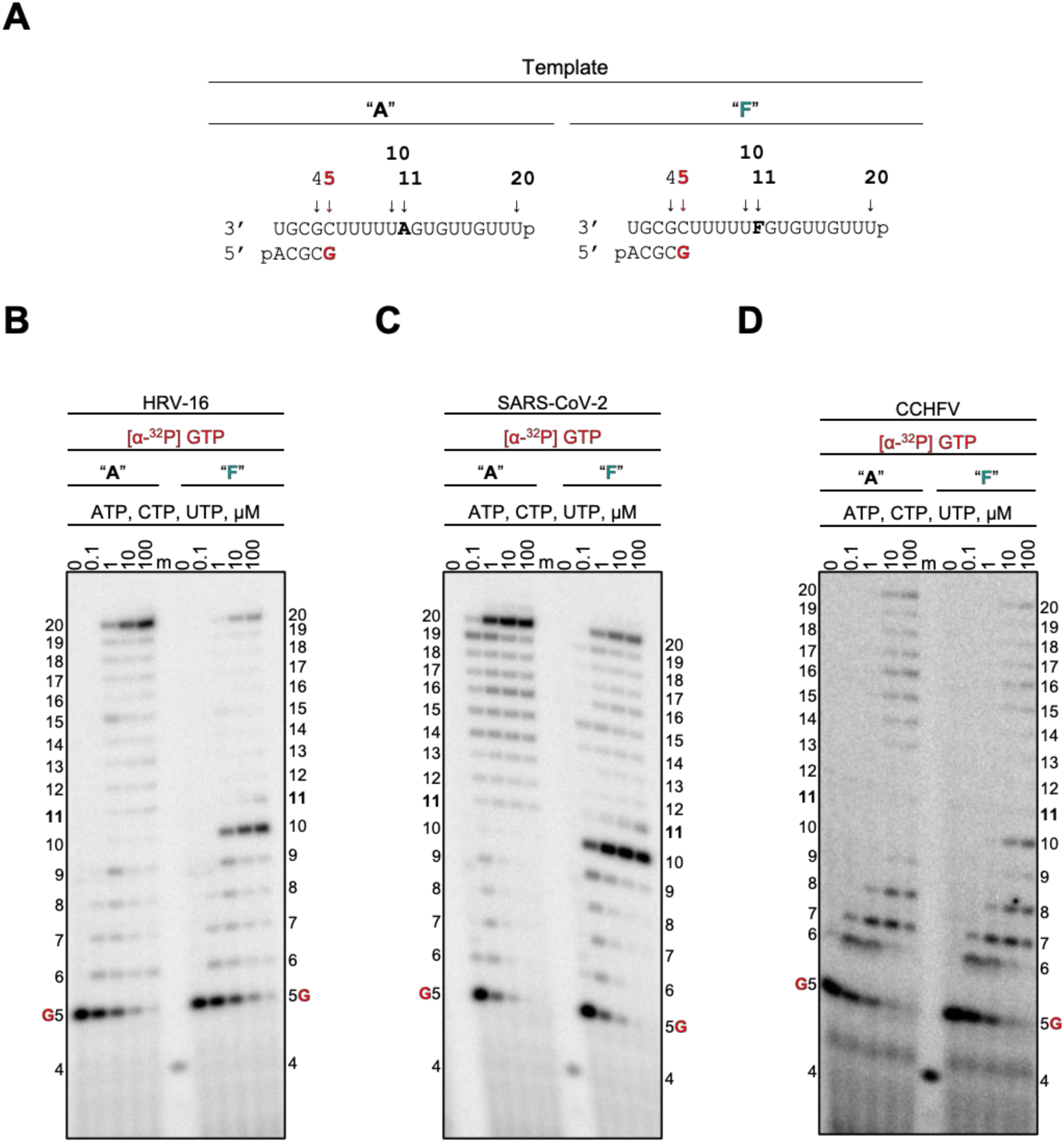
Template-dependent inhibition of HRV-16, SARS-CoV-2, and CCHFV RdRp by a single embedded 4′-FlA residue at position 11. (**A**) RNA primer/template with an embedded adenosine (Template “A”, *left*) and (Template “F”, *right*) position 11. G indicates incorporation of [α-^32^P]-GTP at position 5. Migration pattern of RNA products resulting from (**B**) HRV-16 RdRp-, (**C**) SARS-CoV-2 RdRp-, (**D**) and CCHFV RdRp-catalyzed RNA synthesis supplemented with increasing NTP concentrations using Template “A” (*left*) and Template “F” (*right*). A 5′-^32^P-labelled 4-nt primer (4) serves as a size marker (m). RNA products formed at and beyond the asterisk indicate slippage products that may be a result of sequence-dependent slippage events or RdRp nucleotide transferase activity. HRV, human rhinovirus; SARS-CoV-2, severe acute respiratory syndrome coronavirus 2; CCHFV, Crimean-Congo hemorrhagic fever virus; RdRp, RNA-dependent RNA polymerase.

**Figure S13.**
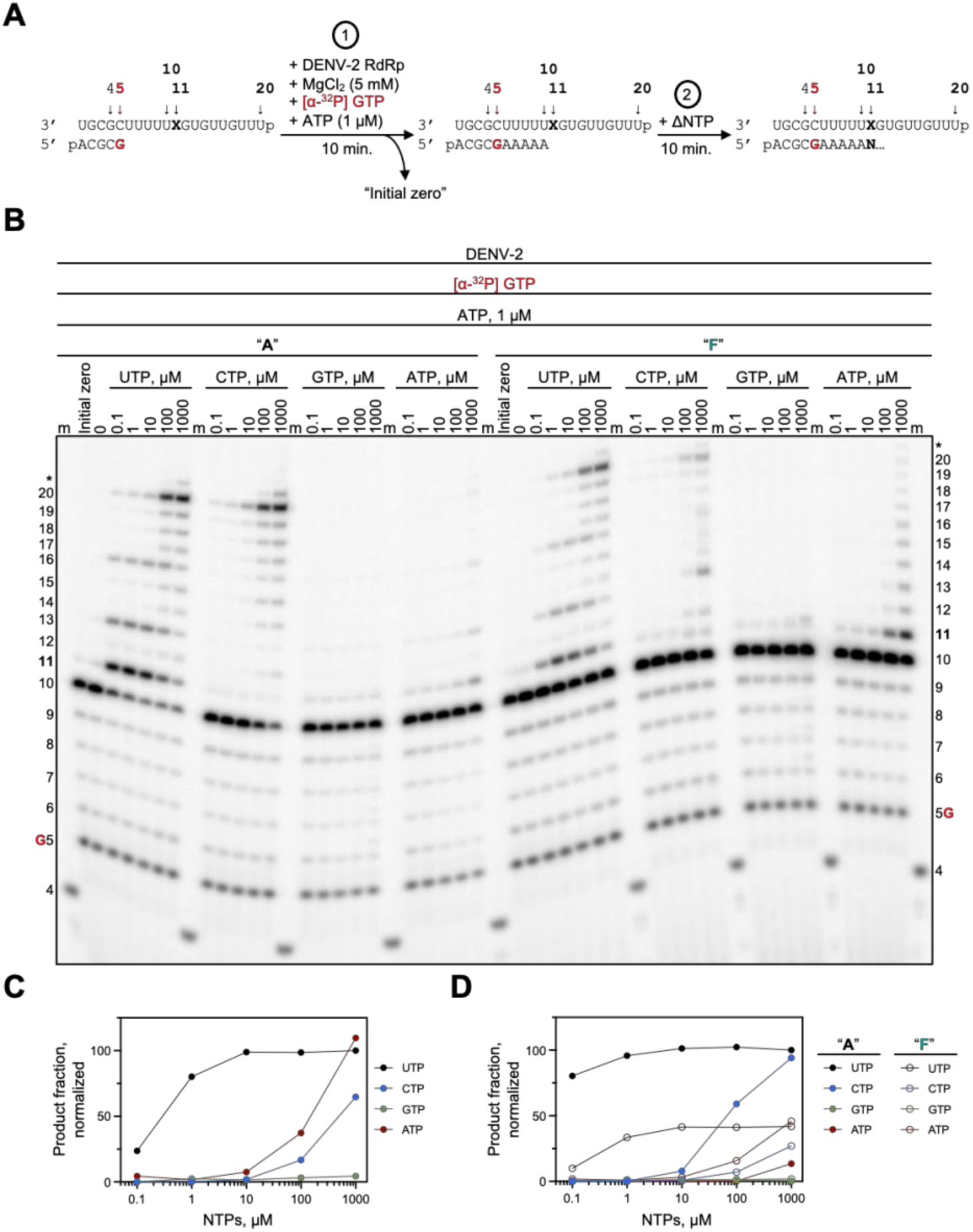
Impairment of NTP incorporation opposite a template-embedded 4′-FlA. (**A**) RNA primer and RNA template with either an embedded adenosine (Template “A”) or (Template “F”) at position 11 (indicated as “X” in the diagram). G indicates incorporation of [α-^32^P]-GTP at position 5. *Step 1*: DENV-2 RdRp, [α-^32^P]-GTP, ATP, and MgCl_2_ are added to the reaction mixture, allowing for RNA synthesis up to position 11. After 10 minutes, the first zero lane reaction (“Initial zero”) is withdrawn. *Step 2*: After the 10 minutes, increasing concentrations (0.1, 1, 10, 100, 1000 µM) of the various NTPs are added to the reaction to allow for incorporation at and beyond position 11. A second zero (“0”) has no NTP added. After 10 minutes, reactions are stopped with formamide/EDTA mixture. (**B**) Migration pattern of RNA products resulting from DENV-2 RdRp-catalyzed RNA synthesis with increasing NTP concentrations using Template “A” (*left*) and Template “F” (*right*). A 5′-^32^P-labelled 4-nt primer (4) serves as a size marker (m). RNA products formed at and beyond the asterisk may be a result of sequence-dependent slippage events or RdRp nucleotide transferase activity. (**C**) Graphic representation of product fraction on Template “F” as a function of NTP concentration. Product fraction is defined as RNA products generated beyond position 10 in each lane, divided by the total signal in that lane, normalized to the UTP lane with the highest product fraction. (**D**) Graphic representation of product fraction on Template “A” and Template “F” as a function of NTP concentration. Product fraction is defined as RNA products generated beyond position 10 in each lane, divided by the total signal in that lane, normalized to the UTP lane on Template “**A**” with the highest product fraction. DENV-2, dengue virus; RdRp, RNA-dependent RNA polymerase.

**Figure S14.**
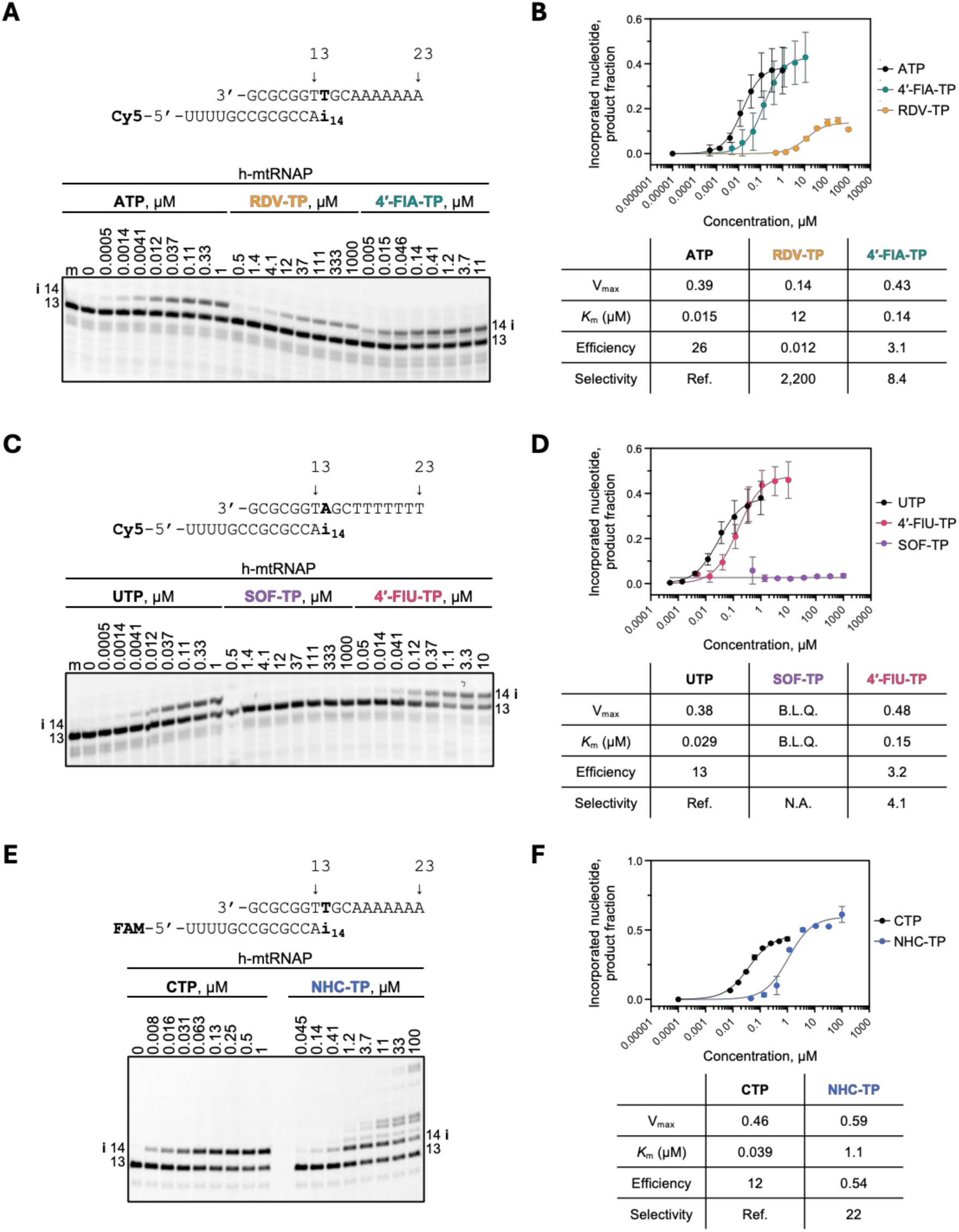
Incorporation of nucleotide analogs by h-mtRNAP. (**A**) RNA primer/DNA template substrate (*Top*) used in the RNA synthesis assays to evaluate RDV-TP and 4′-FlA-TP as a substrate for incorporation as an A-analog. Position 13 indicates the length of the Cy5-labelled primer. Position 14 (“i”) allows incorporation of ATP, RDV-TP, or 4′-FlA-TP. (*Bottom*) NTP incorporation was monitored with purified h-mtRNAP DdRp complex in the presence of RNA primer/DNA template, MgCl_2_, and increasing concentrations of ATP, RDV-TP, or 4′-FlA-TP. Lane m illustrates the migration pattern of the Cy5-labelled 13-nucleotide-long primer. (**B**) (*Top*) Graphic representation of the data for incorporation of ATP, RDV-TP, and 4′-FlA-TP. (*Bottom*) Kinetic parameters and efficiency of ATP, RDV-TP, and 4′-FlA-TP incorporation by h-mtRNAP DdRp. Efficiency is determined as the quotient of V_max_ over *K*_m_. Selectivity is the quotient of ATP efficiency over ATP-analog efficiency. (**C**) RNA primer/DNA template substrate (*Top*) used in the RNA synthesis assays to evaluate SOF-TP and 4′-FlU-TP as a substrate for incorporation. Position 13 indicates the length of the Cy5-labelled primer. Position 14 (“i”) allows incorporation of UTP, SOF-TP, or 4′-FlU-TP. (*Bottom*) NTP incorporation was monitored with purified h-mtRNAP DdRp complex in the presence of RNA primer/DNA template, MgCl_2_, and increasing concentrations of UTP, SOF-TP, or 4′-FlU-TP. Lane m illustrates the migration pattern of the Cy5-labelled 13-nucleotide-long primer. (**D**) (*Top*) Graphic representation of the data for incorporation of UTP, SOF-TP, and 4′-FlU-TP. (*Bottom*) Kinetic parameters and efficiency of UTP, SOF-TP, and 4′-FlU-TP incorporation by h-mtRNAP DdRp. Efficiency is determined as the quotient of V_max_ over *K*_m_. Selectivity is the quotient of UTP efficiency over UTP-analog efficiency. (**E**) RNA primer/DNA template substrate (*Top*) used in the RNA synthesis assays to evaluate NHC-TP as a substrate for incorporation. Position 13 indicates the length of the FAM-labelled primer. Position 14 (“i”) allows incorporation of CTP or NHC-TP. (*Bottom*) NTP incorporation was monitored with purified h-mtRNAP DdRp complex in the presence of RNA primer/DNA template, MgCl_2_, and increasing concentrations of CTP or NHC-TP. Lane m illustrates the migration pattern of the FAM-labelled 13-nucleotide-long primer. (**F**) (*Top*) Graphic representation of the data for incorporation of CTP and NHC-TP. (*Bottom*) Kinetic parameters and efficiency of CTP and NHC-TP incorporation by h-mtRNAP DdRp. Efficiency is determined as the quotient of V_max_ over *K*_m_. Selectivity is the quotient of UTP efficiency over UTP-analog efficiency. Below the level of quantification, B.L.Q.

**Figure S15.**
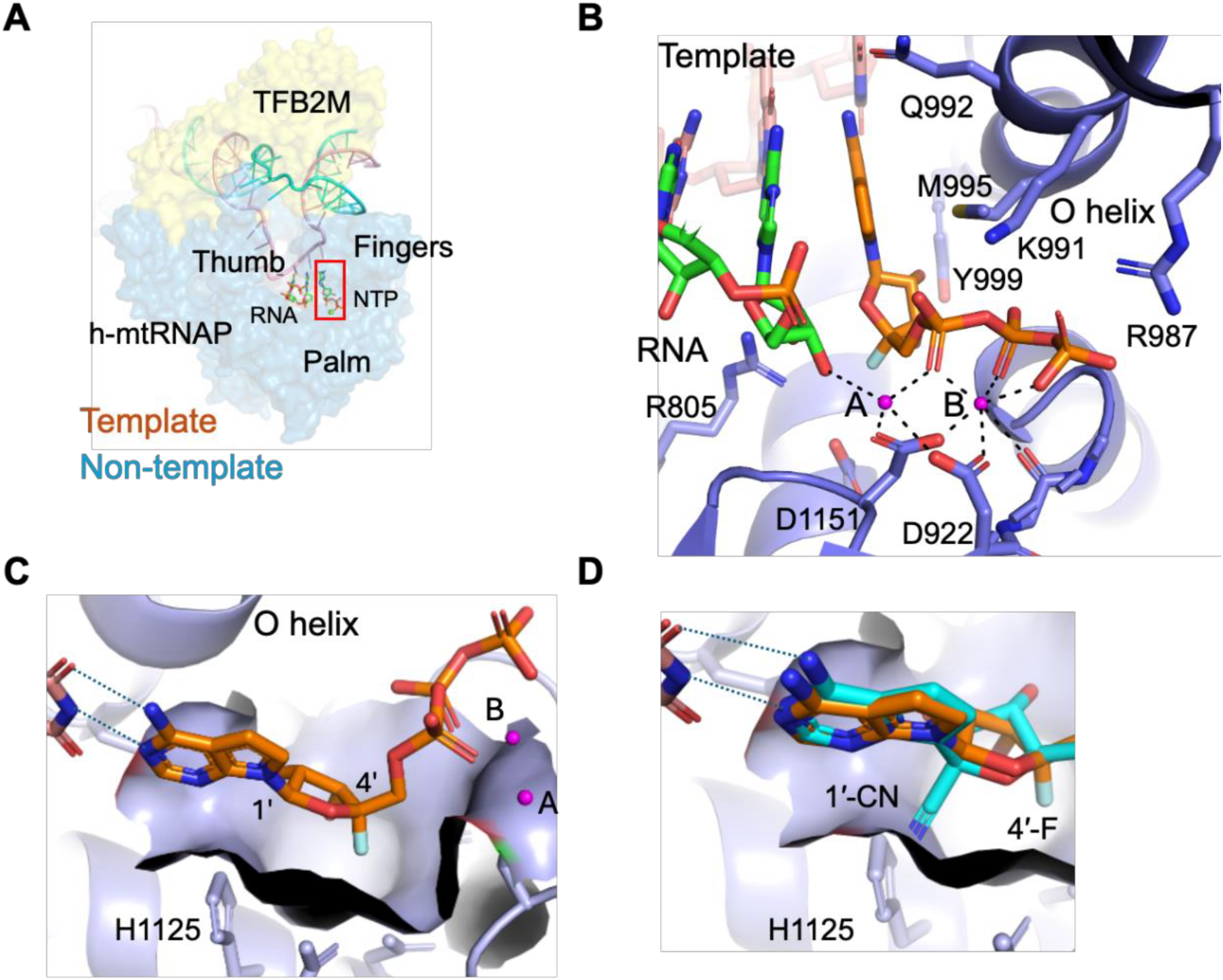
Modelled binding of 4′-FlA-TP in the human mitochondrial RNA polymerase (h-mtRNAP) initiation catalytic complex. (**A**) Structure of the h-mtRNAP initiation complex (PDB: 9R96) (3), which is composed of h-mtRNAP (blue), transcription factor TFB2M (yellow), non-template (cyan), template (pink), and incoming GTP (green). (**B**) The nucleotide analog 4′-FlA-TP is modelled at the N-site within h-mtRNAP. Metal A and B are represented with pink spheres. (**C**) The 4′-fluoro group of 4′-FlA-TP fits well within the adjacent molecular surface of h-mtRNAP. (**D**) In contrast to 4′-FlA-TP, the 1′-cyano group of RDV-TP appears to clash with h-mtRNAP, suggesting it is a poor substrate.

**Figure S16.**
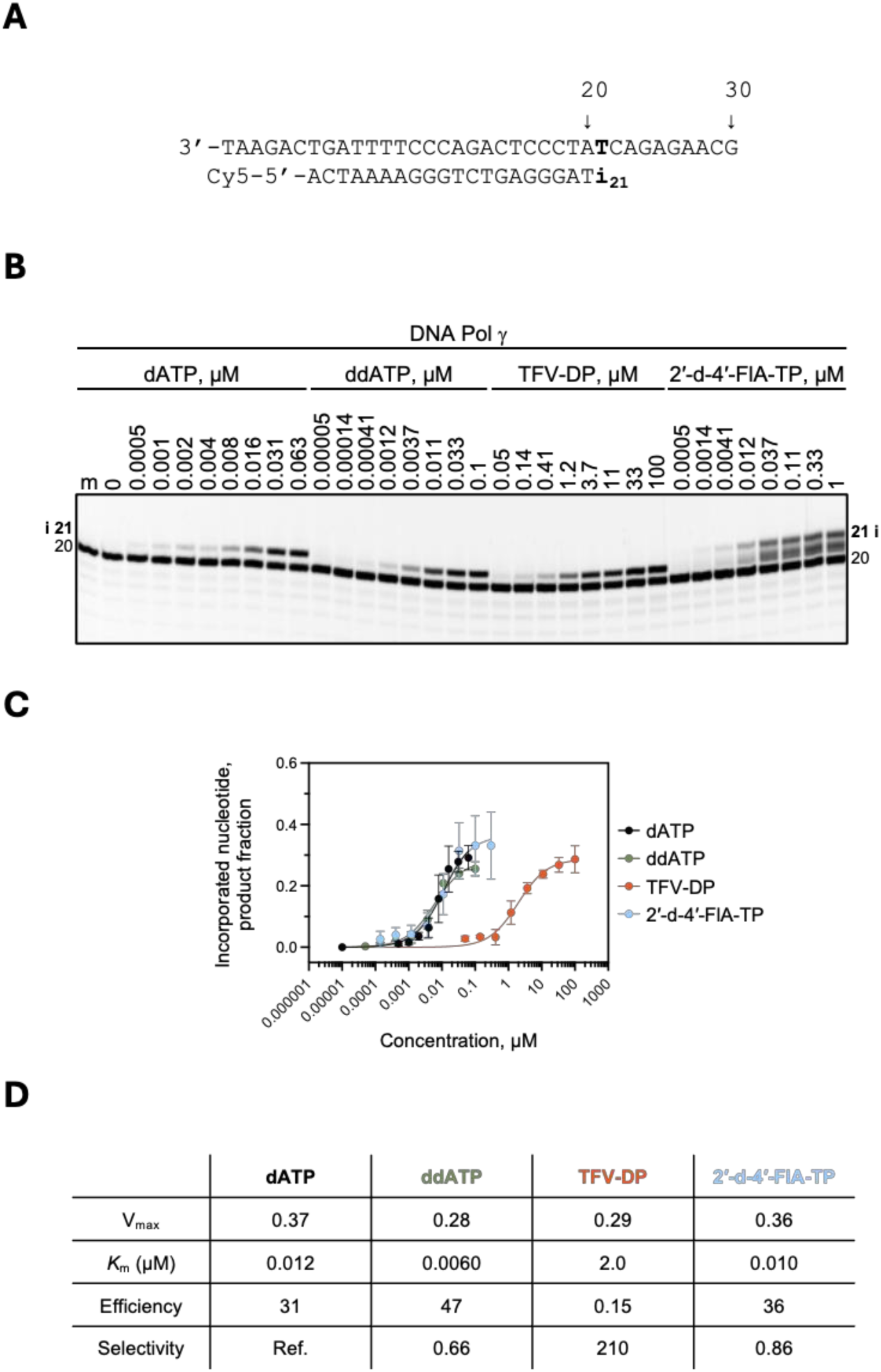
Incorporation of nucleotide analogs by Pol γ. (**A**) DNA primer/template substrate used in the DNA synthesis assays for Pol γ and KF to evaluate ddATP, TFV-DP, and 2′-deoxy-4′-FlA-TP as a substrate for incorporation as an A-analog incorporation is shown above the gel. Position (“i”) allows incorporation of dATP, ddATP, TFV-DP, or 2′-deoxy-4′-FlA-TP. (**B**) Nucleotide incorporation was monitored with purified Pol γ DdDp complex in the presence of Cy5-labelled primer annealed to DNA template, MgCl_2_, and increasing concentrations of dATP, ddATP, TFV-DP, or 2′-deoxy-4′-FlA-TP. Lane m illustrates the migration pattern of the Cy5-labelled primer used as a marker. (**C**) Graphic representation of dATP, ddATP, TFV-DP, and 2′-deoxy-4′-FlA-TP incorporation. (**D**) Kinetic parameters and efficiency of dATP, ddATP, TFV-DP, and 2′-deoxy-4′-FlA-TP incorporation by Pol γ DdDp. Efficiency is determined as the quotient of V_max_ over *K*_m_. Selectivity is the quotient of ATP efficiency over ATP-analog efficiency.

**Figure S17.**
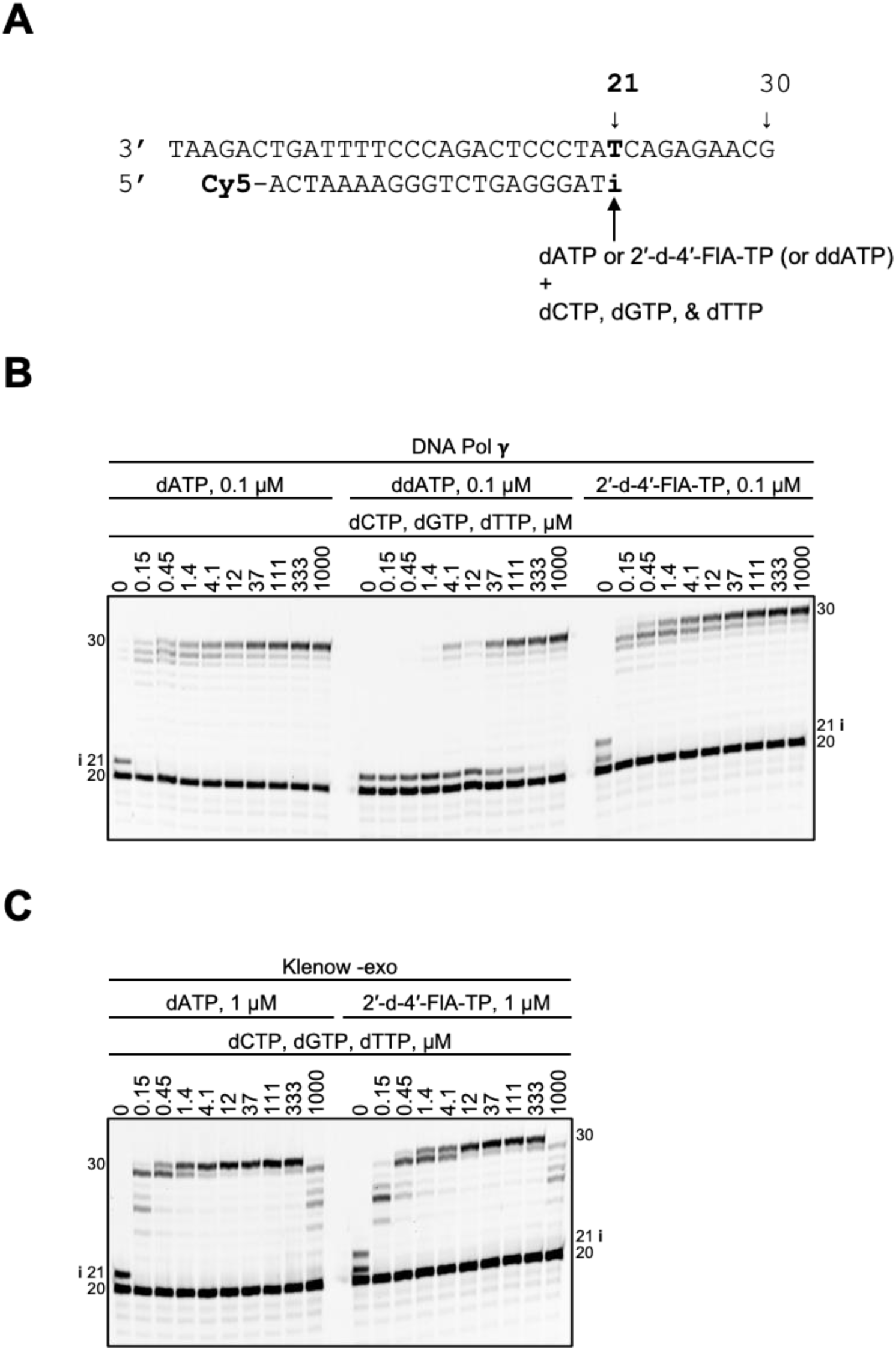
DNA Pol *γ* and Klenow DdDp-catalyzed DNA synthesis and inhibition patterns following a single incorporation of dATP or 2′-d-4′-FlA-TP as a function of nucleotide concentration. (**A**) DNA primer/template supporting a single incorporation of dATP or 2′-d-4′-FlA-TP at position 21 (“i”). The DNA primer is 5′-Cy5 labelled for signal detection of synthesized DNA products. (**B**) Migration pattern of DNA products resulting from DNA Pol *γ* DdDp-catalyzed extension following incorporation of dATP, ddATP, or 2′-d-4′-FlA-TP at position 14 (“i”) at increasing concentrations of dCTP, dGTP, and dTTP. (**C**) Migration pattern of DNA products resulting from Klenow DdDp-catalyzed extension following incorporation of dATP or 2′-d-4′-FlA-TP at position 14 (“i”) at increasing concentrations of dCTP, dGTP, and dTTP.

### Synthetic chemistry

#### 4′-FlA-TP (mCOU991 or V2583902) (Scheme in Figure S16)

To a solution Compound **1** (130 mg, 457 μmol, 1.00 *eq*), proton sponge (97.8 mg, 457 μmol, 1.00 *eq*) in PO (OMe)_3_ (1.30 mL) was added POCl_3_ (104 mg, 685 μmol, 63.5 μL, 1.80 *eq*) at -10 °C. The mixture was stirred at 25 °C for 1 hr under N_2_. **LCMS** (RT = 0.280min) showed compound **1** was consumed completely and desired mass was detected. The crude product (182 mg, crude) was yellow liquid was used into the next step without further purification. To a solution of the crude product (182 mg, 347 μmol, 1.00 *eq*) in PO(OMe)_3_ (1.30 mL) was added (Bu_3_N)_2_H_4_P_2_O_7_ (0.6 M, 2.30 mL, 3.00 *eq*) and Bu_3_N (257 mg, 1388 μmol, 4.00 eq) in at -10 °C. The mixture was stirred at 25 °C for 40 mins. **LCMS** (RT = 0.272 min) showed crude product was consumed completely and one main peak with desired mass was detected. Add 1.00 M TEAB to adjust pH 7. The solution was diluted with H_2_O 2.00 mL and extracted with MTBE (2.00 mL x 3), the aqueous phase purified by DEAE Sephadex column with an elution gradient of 0 to 1.00 M TEAB, evaporated to obtain a colorless oil. The residue was purified by prep-HPLC. Collected and lyophilized. **V2583902** (13.0 mg, 24.0 μmol, 9.75% yield, 97.0% purity) was obtained as a brown solid.

**1H NMR:** 400 MHz, D_2_O

δ ppm 8.15 (s, 1H), 7.32 (d, J = 3.60 Hz, 1H), 6.67 (d, J = 4.0 Hz, 1H), 6.44 (d, J = 2.0Hz, 1H), 4.80-4.78 (m, 1H), 4.39-4.29 (m, 3H).

**31P NMR:** 162 MHz, D_2_O

δ ppm -10.4--10.1 (1P), -11.9--11.8 (1P), -22.9--22.7 (1P)

**19P NMR:** 376 MHz, D_2_O

δ ppm -120.4 (1F)

**1H NMR:** 400 MHz, D_2_O

δ ppm 8.15 (s, 1H), 7.32 (d, J = 3.60 Hz, 1H), 6.67 (d, J = 4.0 Hz, 1H), 6.44 (d, J = 2.0Hz, 1H), 4.80-4.78 (m, 1H), 4.39-4.29 (m, 3H).

**31P NMR:** 162 MHz, D_2_O

δ ppm -10.4--10.1 (1P), -11.9--11.8 (1P), -22.9--22.7 (1P)

**19P NMR:** 376 MHz, D_2_O

δ ppm -120.4 (1F)

#### 2′-deoxy-4′-FlA-TP (mCQT310 or V2870202) (Scheme in Figure S17)

Charge cpd **1** (20.0 g, 79.9 mmol 1.00 eq) into (1000 mL bottle) R1. Charge THF (400 mL, 20.0 V) into R1. Charge imidazole (15.56 g, 229 mmol, 2.86 eq) into R1. Charge PPh_3_ (52.4 g, 200mmol, 2.50 eq) into R1. Charge Py (86.6 g, 1.09 mol, 13.7 eq) into R1. Charge I_2_ (40.57 g, 160mmol, 2.00 eq) into R1 at 0 °C. Stir R1 for 4 hrs at 25 °C. Take sample for analysis. (**LCMS**: RT = 0.877 min). The reaction mixture was quenched with sat. aq. Na_2_S_2_O_3_ (400 mL), extracted with EtOAc (400 mL x 2). The organic layer was dried with Na_2_SO_4_, filtered and concentrated under reduced pressure to give a residue. The residue was purified by column chromatography (SiO_2_, petroleum ether/ethyl acetate=1/1 to 0/1 and DCM/MeOH=10/1). Obtain cpd **2** (12.6 g, 35.2 mmol, 43.6% yield) was white solid.

**TLC**: Dichloromethane: Methanol = 5/1, Rf = 0.56

**LCMS:** RT=0.839 min

**1H NMR:** 400 MHz, MeOH

δ ppm 8.05 (d, *J* = 6.8 Hz, 1H), 7.35 (d, *J* = 2.8 Hz, 1H), 6.62-6.59 (m, 2H), 4.43-4.40 (m, 1 H), 3.92-3.90 (m, 1H), 3.45-3.36 (m, 2H), 2.66-2.64 (m, 1H), 2.34-2.321 (m, 1H).

Charge cpd **2** (7.30 g, 20.3 mmol, 1.00 eq) into (250 mL bottle) R1. Charge sodium;methanolate (73.0 mL) into R1. Stir R1 for 4 hrs at 60 °C. Take sample for analysis. (**LCMS**: RT = 0.758 min). The reaction solution was added acetic acid to adjust pH 7. The mixture solution was concentrated under reduced pressure to give a residue. The residue was purified by column chromatography (SiO_2_, petroleum ether/ethyl acetate=1/1 to 0/1 and DCM/MeOH=10/1). Obtain cpd **3** (3.00 g, 12.9 mmol, 63.7% yield) was white solid.

**TLC**: Petroleum ether/Ethyl acetate = 0/1, Rf = 0.20

**LCMS**: RT=0.737 min

Charge cpd **3** (2.80 g, 12.1 mmol, 1.00 eq) into (100 mL bottle) R1. Charge ACN (28.0 mL, 10.0 V) into R1. Charge DMAP (0.74 g, 6.03 mmol, 0.05 eq) into R1. Charge Ac_2_O (4.923 g, 48.2 mmol, 4.00 eq) into R1. Stir R1 for 4 hrs at 30 °C. Take sample for analysis. (**LCMS**: RT = 0.911 min). The solution was concentrated under reduced pressure to give a residue. The residue was diluted with ethyl acetate (100 mL) and extracted with H_2_O (100 mL x 2). The combined organic layer was dried over anhydrous Na_2_SO_4_, filtered and concentrated under reduced pressure to give a residue. Combine with MT01765-54. The residue was purified by prep-HPLC (neutral condition: column: Welch Xtimate C18 250 x 70 mm x10 um; mobile phase: [A: H_2_O (10mM NH_4_HCO_3_); B: ACN];B%: 10.00%-46.00%,17.00 min;flow rate: 150.00ml/min). Obtain cpd **4** (1.51 g, 4.77mmol, 34.1% yield) was white solid.

**LCMS**: RT = 0.915 min

**1H NMR:** 400 MHz, MeOH

δ ppm 8.52 (s, 1H), 7.48 (m, 1H), 6.96-6.91 (m, 2H), 5.97-5.95 (m, 1 H), 4.46 (d, *J* = 6.8 Hz, 1H), 4.34 (d, *J* = 6.8 Hz, 1H), 2.99-2.95 (m, 1H), 2.66 (m, 1H), 2.28 (s, 3H), 2.12 (s, 3H).

Charge cpd **4** (1.13 g, 3.57 mmol, 1.00 eq) into (100 mL bottle) R1. Charge THF (25.0 mL 22.1 V) into R1. Charge 3HF.TEA (1.73 g, 10.72 mmol, 3.00 eq) into R1 at 0 °C. Charge NIS (1.45 g, 6.43 mmol, 1.80 eq) into R1 at 0 °C. Stir R1 for 3 hrs at 25 °C. Take sample for analysis. (**LCMS**: RT = 1.051 min). The solution was concentrated under reduced pressure to give a residue. The residue was purified by column chromatography (SiO_2_, petroleum ether/ethyl acetate=5/1 to 0/1). Obtain cpd 5 (1.28 g, 2.77 mmol, 77.5% yield) as yellow oil.

**TLC**: Petroleum ether/Ethyl acetate **=** 0:1, Rf = 0.60

**LCMS:** RT = 1.052 min

Charge cpd **5** (1.28 g, 2.77 mmol, 1.00 eq) into (100 mL bottle) R1. Charge DMF (25.0 mL 19.5 V) into R1. Charge potassium;benzoate (2.22 g, 13.8 mmol, 5.00 eq) into R1. Charge 18-crown-6 (1.47 g, 5.54 mmol, 2.00 eq) into R1under N_2_ protection. The reaction mixture was stirred for 16 hrs at 100 °C. Take sample for analysis. (**LCMS**: RT = 1.143 min). The solution was concentrated under reduced pressure to give a residue. Combine with MT01765-61. The residue was diluted with ethyl acetate (50.0 mL) and extracted with H_2_O (50.0 mL x 2). The combined organic layer, was dried over anhydrous Na_2_SO_4_, filtered and concentrated under reduced pressure to give a residue. The residue was purified by prep-HPLC (neutral condition: column: WePure Biotech XP tC18 250 x 70 x 10 um;mobile phase: [A: H_2_O(10 mM NH_4_HCO_3_);B: ACN];B%: 25.0%-60.0%, 20.00min;flow rate:150.00ml/min). Obtain cpd **6** (0.40 g, 876μmol, 31.6% yield) as yellow solid.

**LCMS**: RT = 1.144 min

Charge cpd **6** (0.40 g, 876 μmol, 1.00 eq) into (100 mL bottle) R1. Charge methanamine (7.7 mL) into R1 under N_2_ protection. The reaction mixture was stirred for 4 hrs at 30 °C. Take sample for analysis. (**LCMS**: RT = 0.544 min). The solution was concentrated under reduced pressure to give a residue. The residue was added MeOH 1mL and the solution was added to 20 mL DCM. Increase the spinning speed of centrifuge to 4500 rpm and spinning for 5 min. Collect solid. The solid was added 5.00 mL ACN. Increase the spinning speed of centrifuge to 4500 rpm and spinning for 5 min. Collect solid. Obtain cpd **7** (0.131 g, 488 μmol, 55.7% yield) as off-white solid.

**LCMS:** RT = 0.819 min

**1H NMR:** 400 MHz MeOH

δ ppm 8.08 (s, 1H),7.29 (d, *J* = 4.0Hz, 1H), 6.73-6.70 (m, 1H), 6.62 (d, *J* = 4.0Hz, 1H), 3.77-3.74 (m, 2H), 2.66-2.59 (m, 2H).

**19F NMR:** 376 MHz, MeOH

δ ppm -127.0 (1F)

Charge cpd **7** (0.10 g, 373 μmol, 1.00 eq) into R1 (a thumb flask). Charge proton sponge (80.0 mg, 373μmol, 1.00 eq) into R1 under N_2_ protection. Charge PO(OMe)_3_ (1.00 mL 10.0 V) into R1 under N_2_ protection. Charge POCl_3_ (86.0 mg, 561μmol, 1.50 eq) into R1 at -10 °C under N_2_ protection. Stir R1 for 1 hr at 15 °C. Take sample for analysis. (**LCMS**: RT = 0.422 min). Obtain crude product (0.143 g, crude) as brown liquid.

To a solution of the crude product (0.143 g, 371 μmol 1.00 eq) in PO(OMe)_3_ (1.00 mL) was added (Bu_3_N)_2_H_4_P_2_O_7_ (0.6 M, 1.11 mmol, 1.86 mL, 3.00 eq) at -10 °C under N_2_ protection. Stir R1 for 1 hr at 15 °C. Take sample for analysis. (**LCMS**: RT = 0.338 min). The reaction solution was added TEAB 1mL. The solution was through a DEAE Sephadex column with an elution gradient of 0 to 1.00 M TEAB at 25°C. Fractions of gradient 0.6-0.9 M were collected and concentrated under reduced pressure to give a residue at 30 °C. The residue was purified by prep-HPLC (neutral condition: column: NanoQ-15L 250 x 150 x 15 UM; mobile phase: [A: H_2_O; B: H_2_O (1M TEAB)]; B%: 0.00%-45.00%, 70.00min). Obtain **V2870202** (16.7 mg, 32.8 μmol, 8.88% yield, TEA) as white solid

**LCMS:** RT = 9.043 min

**1H NMR:** 400 MHz, D_2_O

δ ppm 8.10 (s, 1H), 7.40 (m, 1H), 6.68 (m, 1H), 5.02-4.97 (m, 1H), 4.31-4.26 (m, 1H), 2.65-2.54 (m, 2H).

**31P NMR:** 162 MHz, D_2_O

δ ppm -10.5--10.6 (1P), -11.7--11.9 (1P), -23.9--23.2(1P)

**19F NMR:** 376 MHz, D_2_O

δ ppm -123.5 (1F)

## References

1. Draghia-Akli, R., Hill, N.M., Altevogt, B., Bradley, K., Chibale, K., Cihlar, T., Clinch, B., Demarest, J.F. and Neyts, J. The Indispensable Value of Small-Molecule Antivirals in Epidemic and Pandemic Preparedness. Clin Infect Dis, 2025. 10.1093/cid/ciaf476

2. (2024). World Health Organization. Pathogens prioritization: a scientific framework for epidemic and pandemic research preparedness. Meeting Report. https://cdn.who.int/media/docs/default-source/consultation-rdb/prioritization-pathogens-v6final.pdf?sfvrsn=c98effa7_7&download=true (13 January 2026).

3. Venkataraman, S., Prasad, B. and Selvarajan, R. RNA Dependent RNA Polymerases: Insights from Structure, Function and Evolution. Viruses, 2018; 10. 10.3390/v10020076

4. Seley-Radtke, K.L., Thames, J.E. and Waters, C.D., 3rd. Broad spectrum antiviral nucleosides-Our best hope for the future. Annu Rep Med Chem, 2021; 57: 109–132. 10.1016/bs.armc.2021.09.001

5. De Clercq, E. and Li, G. Approved Antiviral Drugs over the Past 50 Years. Clin Microbiol Rev, 2016; 29: 695–747. 10.1128/CMR.00102-15

6. Peersen, O.B. A Comprehensive Superposition of Viral Polymerase Structures. Viruses, 2019; 11. 10.3390/v11080745

7. Kamzeeva, P.N., Aralov, A.V., Alferova, V.A. and Korshun, V.A. Recent Advances in Molecular Mechanisms of Nucleoside Antivirals. Curr Issues Mol Biol, 2023; 45: 6851–6879. 10.3390/cimb45080433

8. Mulangu, S., Dodd, L.E., Davey, R.T., Jr., Tshiani Mbaya, O., Proschan, M., Mukadi, D., Lusakibanza Manzo, M., Nzolo, D., Tshomba Oloma, A., Ibanda, A. et al. A Randomized, Controlled Trial of Ebola Virus Disease Therapeutics. N Engl J Med, 2019; 381: 2293–2303. 10.1056/NEJMoa1910993

9. (2020). U.S. Food and Drug Administration Approves Gilead’s Antiviral Veklury® (remdesivir) for Treatment of COVID-19. https://www.gilead.com/news/news-details/2020/us-food-and-drug-administration-approves-gileads-antiviral-veklury-remdesivir-for-treatment-of-covid-19

10. Radoshitzky, S.R., Iversen, P., Lu, X., Zou, J., Kaptein, S.J.F., Stuthman, K.S., Van Tongeren, S.A., Steffens, J., Gong, R., Truong, H. et al. Expanded profiling of Remdesivir as a broad-spectrum antiviral and low potential for interaction with other medications in vitro. Sci Rep, 2023; 13: 3131. 10.1038/s41598-023-29517-9

11. Lo, M.K., Jordan, R., Arvey, A., Sudhamsu, J., Shrivastava-Ranjan, P., Hotard, A.L., Flint, M., McMullan, L.K., Siegel, D., Clarke, M.O. et al. GS-5734 and its parent nucleoside analog inhibit Filo-, Pneumo-, and Paramyxoviruses. Sci Rep, 2017; 7: 43395. 10.1038/srep43395

12. Walker, S.M., Gordon, C.J., Tchesnokov, E.P., Sun, L., Zou, J., Xie, X., Riola, N.C., Cutillas, V., Du Pont, V., Zhao, X. et al. 1′- and 4′-Cyano Modified Adenosine Analogs Against Prototypic Flavivirus RNA-Dependent RNA Polymerases. Viruses, 2026; 18: 257.

13. Pruijssers, A.J., George, A.S., Schafer, A., Leist, S.R., Gralinksi, L.E., Dinnon, K.H., 3rd, Yount, B.L., Agostini, M.L., Stevens, L.J., Chappell, J.D. et al. Remdesivir Inhibits SARS-CoV-2 in Human Lung Cells and Chimeric SARS-CoV Expressing the SARS-CoV-2 RNA Polymerase in Mice. Cell Rep, 2020; 32: 107940. 10.1016/j.celrep.2020.107940

14. Xie, X., Muruato, A.E., Zhang, X., Lokugamage, K.G., Fontes-Garfias, C.R., Zou, J., Liu, J., Ren, P., Balakrishnan, M., Cihlar, T. et al. A nanoluciferase SARS-CoV-2 for rapid neutralization testing and screening of anti-infective drugs for COVID-19. Nat Commun, 2020; 11: 5214. 10.1038/s41467-020-19055-7

15. Verwimp, S., Wagoner, J., Arenas, E.G., De Coninck, L., Abdelnabi, R., Hyde, J.L., Schiffer, J.T., White, J.M., Matthijnssens, J., Neyts, J. et al. Combinations of approved oral nucleoside analogues confer potent suppression of alphaviruses in vitro and in vivo. Antiviral Res, 2025; 239: 106186. 10.1016/j.antiviral.2025.106186

16. Wahl, A., Gralinski, L.E., Johnson, C.E., Yao, W., Kovarova, M., Dinnon, K.H., 3rd, Liu, H., Madden, V.J., Krzystek, H.M., De, C. et al. SARS-CoV-2 infection is effectively treated and prevented by EIDD-2801. Nature, 2021; 591: 451–457. 10.1038/s41586-021-03312-w

17. Sheahan, T.P., Sims, A.C., Zhou, S., Graham, R.L., Pruijssers, A.J., Agostini, M.L., Leist, S.R., Schafer, A., Dinnon, K.H., 3rd, Stevens, L.J. et al. An orally bioavailable broad-spectrum antiviral inhibits SARS-CoV-2 in human airway epithelial cell cultures and multiple coronaviruses in mice. Sci Transl Med, 2020; 12. 10.1126/scitranslmed.abb5883

18. Yoon, J.J., Toots, M., Lee, S., Lee, M.E., Ludeke, B., Luczo, J.M., Ganti, K., Cox, R.M., Sticher, Z.M., Edpuganti, V. et al. Orally Efficacious Broad-Spectrum Ribonucleoside Analog Inhibitor of Influenza and Respiratory Syncytial Viruses. Antimicrob Agents Chemother, 2018; 62. 10.1128/AAC.00766-18

19. Ehteshami, M., Tao, S., Zandi, K., Hsiao, H.M., Jiang, Y., Hammond, E., Amblard, F., Russell, O.O., Merits, A. and Schinazi, R.F. Characterization of beta-d-N(4)-Hydroxycytidine as a Novel Inhibitor of Chikungunya Virus. Antimicrob Agents Chemother, 2017; 61. 10.1128/AAC.02395-16

20. Reynard, O., Nguyen, X.N., Alazard-Dany, N., Barateau, V., Cimarelli, A. and Volchkov, V.E. Identification of a New Ribonucleoside Inhibitor of Ebola Virus Replication. Viruses, 2015; 7: 6233–6240. 10.3390/v7122934

21. Stuyver, L.J., Whitaker, T., McBrayer, T.R., Hernandez-Santiago, B.I., Lostia, S., Tharnish, P.M., Ramesh, M., Chu, C.K., Jordan, R., Shi, J. et al. Ribonucleoside analogue that blocks replication of bovine viral diarrhea and hepatitis C viruses in culture. Antimicrob Agents Chemother, 2003; 47: 244–254. 10.1128/AAC.47.1.244-254.2003

22. Bluemling, G.R., Mao, S., Natchus, M.G., Painter, W., Mulangu, S., Lockwood, M., De La Rosa, A., Brasel, T., Comer, J.E., Freiberg, A.N. et al. The prophylactic and therapeutic efficacy of the broadly active antiviral ribonucleoside N(4)-Hydroxycytidine (EIDD-1931) in a mouse model of lethal Ebola virus infection. Antiviral Res, 2023; 209: 105453. 10.1016/j.antiviral.2022.105453

23. U.S. Food & Drug Administration. Coronavirus (COVID-19) Update: FDA Authorizes Additional Oral Antiviral for Treatment of COVID-19 in Certain Adults. 2021.

24. Sloan, A., Gordon, C.J., Prevost, J., Audet, J., Fulton, K., Woolner, E., Chang, M.H., Walker, S.M., Tchesnokov, E.P., Das, K. et al. Inhibitory effects of molnupiravir on Crimean-Congo hemorrhagic fever virus polymerase. NAR Mol Med, 2026; 3: ugaf041. 10.1093/narmme/ugaf041

25. Schrell, L., Fuchs, H.L., Dickmanns, A., Scheibner, D., Olejnik, J., Hume, A.J., Reineking, W., Stork, T., Muller, M., Graaf-Rau, A. et al. Inhibitors of dihydroorotate dehydrogenase synergize with the broad antiviral activity of 4’-fluorouridine. Antiviral Res, 2025; 233: 106046. 10.1016/j.antiviral.2024.106046

26. Welch, S.R., Spengler, J.R., Westover, J.B., Bailey, K.W., Davies, K.A., Aida-Ficken, V., Bluemling, G.R., Boardman, K.M., Wasson, S.R., Mao, S. et al. Delayed low-dose oral administration of 4’-fluorouridine inhibits pathogenic arenaviruses in animal models of lethal disease. Sci Transl Med, 2024; 16: eado7034. 10.1126/scitranslmed.ado7034

27. Lieber, C.M., Kang, H.J., Aggarwal, M., Lieberman, N.A., Sobolik, E.B., Yoon, J.J., Natchus, M.G., Cox, R.M., Greninger, A.L. and Plemper, R.K. Influenza A virus resistance to 4’-fluorouridine coincides with viral attenuation in vitro and in vivo. PLoS Pathog, 2024; 20: e1011993. 10.1371/journal.ppat.1011993

28. Chen, Y., Li, X., Han, F., Ji, B., Li, Y., Yan, J., Wang, M., Fan, J., Zhang, S., Lu, L. et al. The nucleoside analog 4’-fluorouridine suppresses the replication of multiple enteroviruses by targeting 3D polymerase. Antimicrob Agents Chemother, 2024; 68: e0005424. 10.1128/aac.00054-24

29. Lieber, C.M., Aggarwal, M., Yoon, J.J., Cox, R.M., Kang, H.J., Sourimant, J., Toots, M., Johnson, S.K., Jones, C.A., Sticher, Z.M. et al. 4’-Fluorouridine mitigates lethal infection with pandemic human and highly pathogenic avian influenza viruses. PLoS Pathog, 2023; 19: e1011342. 10.1371/journal.ppat.1011342

30. Sourimant, J., Lieber, C.M., Aggarwal, M., Cox, R.M., Wolf, J.D., Yoon, J.J., Toots, M., Ye, C., Sticher, Z., Kolykhalov, A.A. et al. 4’-Fluorouridine is an oral antiviral that blocks respiratory syncytial virus and SARS-CoV-2 replication. Science, 2022; 375: 161–167. 10.1126/science.abj5508

31. Yin, P., May, N.A., Lello, L.S., Fayed, A., Parks, M.G., Drobish, A.M., Wang, S., Andrews, M., Sticher, Z., Kolykhalov, A.A. et al. 4’-Fluorouridine inhibits alphavirus replication and infection in vitro and in vivo. mBio, 2024; 15: e0042024. 10.1128/mbio.00420-24

32. Wang, R., Wang, X., Zhu, J., Li, H. and Liu, W. Effectiveness of nucleoside analogs against Wetland virus infection. Antiviral Res, 2025; 236: 106114. 10.1016/j.antiviral.2025.106114

33. Escaffre, O., Pearson, M.L., Juelich, T.L., Smith, J.K., Zhang, L., Lieber, C.M., Vyshenska, D., Ikegami, T., Haas, G.D., Krueger, R.E. et al. Efficacy of the nucleoside analog 4’-Fluorouridine against Nipah virus in the Syrian hamster model. PLoS Pathog, 2026; 22: e1014093. 10.1371/journal.ppat.1014093

34. Shannon, A. and Canard, B. Kill or corrupt: Mechanisms of action and drug-resistance of nucleotide analogues against SARS-CoV-2. Antiviral Res, 2023; 210: 105501. 10.1016/j.antiviral.2022.105501

35. Yu, C., Chatterjee, A., Riva, L., Wolff, K.C., McNamara, C.W., You, H., Gupta, A.K. and Saleh, O. 2024.

36. Ivanov, M.A., Liudva, G.S., Mukovnia, A.V., Kochetkov, S.N., Tunitskaia, V.L. and Aleksnadrova, L.A. [Synthesis and biological properties of pyrimidine 4’-fluoro nucleosides and 4’-fluoro uridine 5’-O-triphospate]. Bioorg Khim, 2010; 36: 526–534. 10.1134/s1068162010040072

37. Warfield, K.L., Barnard, D.L., Enterlein, S.G., Smee, D.F., Khaliq, M., Sampath, A., Callahan, M.V., Ramstedt, U. and Day, C.W. The Iminosugar UV-4 is a Broad Inhibitor of Influenza A and B Viruses ex Vivo and in Mice. Viruses, 2016; 8: 71. 10.3390/v8030071

38. Edwards, M.R., Pietzsch, C., Vausselin, T., Shaw, M.L., Bukreyev, A. and Basler, C.F. High-Throughput Minigenome System for Identifying Small-Molecule Inhibitors of Ebola Virus Replication. ACS Infect Dis, 2015; 1: 380–387. 10.1021/acsinfecdis.5b00053

39. Welch, S.R., Scholte, F.E.M., Flint, M., Chatterjee, P., Nichol, S.T., Bergeron, E. and Spiropoulou, C.F. Identification of 2’-deoxy-2’-fluorocytidine as a potent inhibitor of Crimean-Congo hemorrhagic fever virus replication using a recombinant fluorescent reporter virus. Antiviral Res, 2017; 147: 91–99. 10.1016/j.antiviral.2017.10.008

40. Bergeron, E., Zivcec, M., Chakrabarti, A.K., Nichol, S.T., Albarino, C.G. and Spiropoulou, C.F. Recovery of Recombinant Crimean Congo Hemorrhagic Fever Virus Reveals a Function for Non-structural Glycoproteins Cleavage by Furin. PLoS Pathog, 2015; 11: e1004879. 10.1371/journal.ppat.1004879

41. Beyleveld, G., White, K.M., Ayllon, J. and Shaw, M.L. New-generation screening assays for the detection of anti-influenza compounds targeting viral and host functions. Antiviral Res, 2013; 100: 120–132. 10.1016/j.antiviral.2013.07.018

42. Tchesnokov, E.P., Gordon, C.J., Woolner, E., Kocinkova, D., Perry, J.K., Feng, J.Y., Porter, D.P. and Gotte, M. Template-dependent inhibition of coronavirus RNA-dependent RNA polymerase by remdesivir reveals a second mechanism of action. J Biol Chem, 2020; 295: 16156–16165. 10.1074/jbc.AC120.015720

43. Baldwin, E.T., van Eeuwen, T., Hoyos, D., Zalevsky, A., Tchesnokov, E.P., Sanchez, R., Miller, B.D., Di Stefano, L.H., Ruiz, F.X., Hancock, M. et al. Structures, functions and adaptations of the human LINE-1 ORF2 protein. Nature, 2024; 626: 194–206. 10.1038/s41586-023-06947-z

44. Lee, H.W., Tchesnokov, E.P., Stevens, L.J., Hughes, T.M., Diefenbacher, M.V., Woolner, E., Kocincova, D., Schultz, D.C., Cherry, S., Sheahan, T.P. et al. Mechanism and spectrum of inhibition of viral polymerases by 2’-deoxy-2’-beta-fluoro-4’-azidocytidine or azvudine. NAR Mol Med, 2025; 2: ugaf029. 10.1093/narmme/ugaf029

45. Gordon, C.J., Walker, S.M., Tchesnokov, E.P., Kocincova, D., Pitts, J., Siegel, D.S., Perry, J.K., Feng, J.Y., Bilello, J.P. and Gotte, M. Mechanism and spectrum of inhibition of a 4’-cyano modified nucleotide analog against diverse RNA polymerases of prototypic respiratory RNA viruses. J Biol Chem, 2024; 300: 107514. 10.1016/j.jbc.2024.107514

46. Tchesnokov, E.P., Bailey-Elkin, B.A., Mark, B.L. and Gotte, M. Independent inhibition of the polymerase and deubiquitinase activities of the Crimean-Congo Hemorrhagic Fever Virus full-length L-protein. PLoS Negl Trop Dis, 2020; 14: e0008283. 10.1371/journal.pntd.0008283

47. Gordon, C.J., Tchesnokov, E.P., Woolner, E., Perry, J.K., Feng, J.Y., Porter, D.P. and Gotte, M. Remdesivir is a direct-acting antiviral that inhibits RNA-dependent RNA polymerase from severe acute respiratory syndrome coronavirus 2 with high potency. J Biol Chem, 2020; 295: 6785–6797. 10.1074/jbc.RA120.013679

48. Tchesnokov, E.P., Feng, J.Y., Porter, D.P. and Gotte, M. Mechanism of Inhibition of Ebola Virus RNA-Dependent RNA Polymerase by Remdesivir. Viruses, 2019; 11. 10.3390/v11040326

49. Tchesnokov, E.P., Raeisimakiani, P., Ngure, M., Marchant, D. and Gotte, M. Recombinant RNA-Dependent RNA Polymerase Complex of Ebola Virus. Sci Rep, 2018; 8: 3970. 10.1038/s41598-018-22328-3

50. Pilotto, S., Sykora, M., Cackett, G., Dulson, C. and Werner, F. Structure of the recombinant RNA polymerase from African Swine Fever Virus. Nat Commun, 2024; 15: 1606. 10.1038/s41467-024-45842-7

51. Fan, H., Walker, A.P., Carrique, L., Keown, J.R., Serna Martin, I., Karia, D., Sharps, J., Hengrung, N., Pardon, E., Steyaert, J. et al. Structures of influenza A virus RNA polymerase offer insight into viral genome replication. Nature, 2019; 573: 287–290. 10.1038/s41586-019-1530-7

52. Weissmann, F., Petzold, G., VanderLinden, R., Huis In ’t Veld, P.J., Brown, N.G., Lampert, F., Westermann, S., Stark, H., Schulman, B.A. and Peters, J.M. biGBac enables rapid gene assembly for the expression of large multisubunit protein complexes. Proc Natl Acad Sci U S A, 2016; 113: E2564–2569. 10.1073/pnas.1604935113

53. Bieniossek, C., Richmond, T.J. and Berger, I. MultiBac: multigene baculovirus-based eukaryotic protein complex production. Curr Protoc Protein Sci, 2008; **Chapter 5**: Unit 5 20. 10.1002/0471140864.ps0520s51

54. Berger, I., Fitzgerald, D.J. and Richmond, T.J. Baculovirus expression system for heterologous multiprotein complexes. Nat Biotechnol, 2004; 22: 1583–1587. 10.1038/nbt1036

55. Gordon, C.J., Lee, H.W., Tchesnokov, E.P., Perry, J.K., Feng, J.Y., Bilello, J.P., Porter, D.P. and Gotte, M. Efficient incorporation and template-dependent polymerase inhibition are major determinants for the broad-spectrum antiviral activity of remdesivir. J Biol Chem, 2022; 298: 101529. 10.1016/j.jbc.2021.101529

56. Loutan, A.J., Yang, B., Connolly, G., Montoya, A., Smiley, R.J., Chatterjee, A.K. and Gotte, M. Bunyaviral Cap-Snatching Endonuclease Activity and Inhibition with Baloxavir-like Inhibitors in the Context of Full-Length L Proteins. Viruses, 2025; 17. 10.3390/v17030420

57. Yin, W., Mao, C., Luan, X., Shen, D.D., Shen, Q., Su, H., Wang, X., Zhou, F., Zhao, W., Gao, M. et al. Structural basis for inhibition of the RNA-dependent RNA polymerase from SARS-CoV-2 by remdesivir. Science, 2020; 368: 1499–1504. 10.1126/science.abc1560

58. Punjani, A., Rubinstein, J.L., Fleet, D.J. and Brubaker, M.A. cryoSPARC: algorithms for rapid unsupervised cryo-EM structure determination. Nat Methods, 2017; 14: 290–296. 10.1038/nmeth.4169

59. Wang, R.Y., Song, Y., Barad, B.A., Cheng, Y., Fraser, J.S. and DiMaio, F. Automated structure refinement of macromolecular assemblies from cryo-EM maps using Rosetta. Elife, 2016; 5. 10.7554/eLife.17219

60. Casanal, A., Lohkamp, B. and Emsley, P. Current developments in Coot for macromolecular model building of Electron Cryo-microscopy and Crystallographic Data. Protein Sci, 2020; 29: 1069–1078. 10.1002/pro.3791

61. Afonine, P.V., Poon, B.K., Read, R.J., Sobolev, O.V., Terwilliger, T.C., Urzhumtsev, A. and Adams, P.D. Real-space refinement in PHENIX for cryo-EM and crystallography. Acta Crystallogr D Struct Biol, 2018; 74: 531–544. 10.1107/S2059798318006551

62. Barad, B.A., Echols, N., Wang, R.Y., Cheng, Y., DiMaio, F., Adams, P.D. and Fraser, J.S. EMRinger: side chain-directed model and map validation for 3D cryo-electron microscopy. Nat Methods, 2015; 12: 943–946. 10.1038/nmeth.3541

63. Williams, C.J., Headd, J.J., Moriarty, N.W., Prisant, M.G., Videau, L.L., Deis, L.N., Verma, V., Keedy, D.A., Hintze, B.J., Chen, V.B. et al. MolProbity: More and better reference data for improved all-atom structure validation. Protein Sci, 2018; 27: 293–315. 10.1002/pro.3330

64. Meng, E.C., Goddard, T.D., Pettersen, E.F., Couch, G.S., Pearson, Z.J., Morris, J.H. and Ferrin, T.E. UCSF ChimeraX: Tools for structure building and analysis. Protein Sci, 2023; 32: e4792. 10.1002/pro.4792

65. Schrodinger, L.L.C. (2015). The PyMOL Molecular Graphics System, Version 1.8.

66. Shen, J., Goovaerts, Q., Ajjugal, Y., De Wijngaert, B., Das, K. and Patel, S.S. Human mitochondrial RNA polymerase structures reveal transcription start site and slippage mechanism. Mol Cell, 2025; 85: 3137–3150 e3137. 10.1016/j.molcel.2025.07.002

67. Cho, A., Saunders, O.L., Butler, T., Zhang, L., Xu, J., Vela, J.E., Feng, J.Y., Ray, A.S. and Kim, C.U. Synthesis and antiviral activity of a series of 1’-substituted 4-aza-7,9-dideazaadenosine C-nucleosides. Bioorg Med Chem Lett, 2012; 22: 2705–2707. 10.1016/j.bmcl.2012.02.105

68. Fung, A., Jin, Z., Dyatkina, N., Wang, G., Beigelman, L. and Deval, J. Efficiency of incorporation and chain termination determines the inhibition potency of 2’-modified nucleotide analogs against hepatitis C virus polymerase. Antimicrob Agents Chemother, 2014; 58: 3636–3645. 10.1128/AAC.02666-14

69. Malone, B.F., Perry, J.K., Olinares, P.D.B., Lee, H.W., Chen, J., Appleby, T.C., Feng, J.Y., Bilello, J.P., Ng, H., Sotiris, J. et al. Structural basis for substrate selection by the SARS-CoV-2 replicase. Nature, 2023; 614: 781–787. 10.1038/s41586-022-05664-3

70. Wu, J., Wang, X., Liu, Q., Lu, G. and Gong, P. Structural basis of transition from initiation to elongation in de novo viral RNA-dependent RNA polymerases. Proc Natl Acad Sci U S A, 2023; 120: e2211425120. 10.1073/pnas.2211425120

71. Kouba, T., Drncova, P. and Cusack, S. Structural snapshots of actively transcribing influenza polymerase. Nat Struct Mol Biol, 2019; 26: 460–470. 10.1038/s41594-019-0232-z

72. Kokic, G., Hillen, H.S., Tegunov, D., Dienemann, C., Seitz, F., Schmitzova, J., Farnung, L., Siewert, A., Hobartner, C. and Cramer, P. Mechanism of SARS-CoV-2 polymerase stalling by remdesivir. Nat Commun, 2021; 12: 279. 10.1038/s41467-020-20542-0

73. Wu, J., Wang, H., Liu, Q., Li, R., Gao, Y., Fang, X., Zhong, Y., Wang, M., Wang, Q., Rao, Z. et al. Remdesivir overcomes the S861 roadblock in SARS-CoV-2 polymerase elongation complex. Cell Rep, 2021; 37: 109882. 10.1016/j.celrep.2021.109882

74. Jin, Z., Kinkade, A., Behera, I., Chaudhuri, S., Tucker, K., Dyatkina, N., Rajwanshi, V.K., Wang, G., Jekle, A., Smith, D.B. et al. Structure-activity relationship analysis of mitochondrial toxicity caused by antiviral ribonucleoside analogs. Antiviral Res, 2017; 143: 151–161. 10.1016/j.antiviral.2017.04.005

75. Lee, H., Hanes, J. and Johnson, K.A. Toxicity of nucleoside analogues used to treat AIDS and the selectivity of the mitochondrial DNA polymerase. Biochemistry, 2003; 42: 14711–14719. 10.1021/bi035596s

76. Johnson, A.A., Ray, A.S., Hanes, J., Suo, Z., Colacino, J.M., Anderson, K.S. and Johnson, K.A. Toxicity of antiviral nucleoside analogs and the human mitochondrial DNA polymerase. J Biol Chem, 2001; 276: 40847–40857. 10.1074/jbc.M106743200

77. Arnold, J.J., Sharma, S.D., Feng, J.Y., Ray, A.S., Smidansky, E.D., Kireeva, M.L., Cho, A., Perry, J., Vela, J.E., Park, Y. et al. Sensitivity of mitochondrial transcription and resistance of RNA polymerase II dependent nuclear transcription to antiviral ribonucleosides. PLoS Pathog, 2012; 8: e1003030. 10.1371/journal.ppat.1003030

78. Keating, G.M. Sofosbuvir: a review of its use in patients with chronic hepatitis C. Drugs, 2014; 74: 1127–1146. 10.1007/s40265-014-0247-z

79. (2013). Gilead Sciences Inc. Sovaldi™ (sofosbuvir) tablets, for oral use: US prescribing information. https://www.gilead.com/-/media/files/pdfs/medicines/liver-disease/sovaldi/sovaldi_patient_pi.pdf (January 30, 2026).

80. Sticher, Z.M., Lu, G., Mitchell, D.G., Marlow, J., Moellering, L., Bluemling, G.R., Guthrie, D.B., Natchus, M.G., Painter, G.R. and Kolykhalov, A.A. Analysis of the Potential for N(4)-Hydroxycytidine To Inhibit Mitochondrial Replication and Function. Antimicrob Agents Chemother, 2020; 64. 10.1128/AAC.01719-19

81. Fenaux, M., Lin, X., Yokokawa, F., Sweeney, Z., Saunders, O., Xie, L., Lim, S.P., Uteng, M., Uehara, K., Warne, R. et al. Antiviral Nucleotide Incorporation by Recombinant Human Mitochondrial RNA Polymerase Is Predictive of Increased In Vivo Mitochondrial Toxicity Risk. Antimicrob Agents Chemother, 2016; 60: 7077–7085. 10.1128/AAC.01253-16

82. Feng, J.Y., Xu, Y., Barauskas, O., Perry, J.K., Ahmadyar, S., Stepan, G., Yu, H., Babusis, D., Park, Y., McCutcheon, K. et al. Role of Mitochondrial RNA Polymerase in the Toxicity of Nucleotide Inhibitors of Hepatitis C Virus. Antimicrob Agents Chemother, 2016; 60: 806–817. 10.1128/AAC.01922-15

83. Eklund, H., Uhlin, U., Farnegardh, M., Logan, D.T. and Nordlund, P. Structure and function of the radical enzyme ribonucleotide reductase. Prog Biophys Mol Biol, 2001; 77: 177–268. 10.1016/s0079-6107(01)00014-1

84. Zhou, S., Hill, C.S., Sarkar, S., Tse, L.V., Woodburn, B.M.D., Schinazi, R.F., Sheahan, T.P., Baric, R.S., Heise, M.T. and Swanstrom, R. beta-d-N4-hydroxycytidine Inhibits SARS-CoV-2 Through Lethal Mutagenesis But Is Also Mutagenic To Mammalian Cells. J Infect Dis, 2021; 224: 415–419. 10.1093/infdis/jiab247

85. Moyle, G. Clinical manifestations and management of antiretroviral nucleoside analog-related mitochondrial toxicity. Clin Ther, 2000; 22: 911–936; discussion 898. 10.1016/S0149-2918(00)80064-8

86. Kewn, S., Hoggard, P.G., Henry-Mowatt, J.S., Veal, G.J., Sales, S.D., Barry, M.G. and Back, D.J. Intracellular activation of 2’,3’-dideoxyinosine and drug interactions in vitro. AIDS Res Hum Retroviruses, 1999; 15: 793–802. 10.1089/088922299310692

87. Kakuda, T.N. Pharmacology of nucleoside and nucleotide reverse transcriptase inhibitor-induced mitochondrial toxicity. Clin Ther, 2000; 22: 685–708. 10.1016/S0149-2918(00)90004-3

88. Kearney, B.P., Flaherty, J.F. and Shah, J. Tenofovir disoproxil fumarate: clinical pharmacology and pharmacokinetics. Clin Pharmacokinet, 2004; 43: 595–612. 10.2165/00003088-200443090-00003

89. Deval, J., Hong, J., Wang, G., Taylor, J., Smith, L.K., Fung, A., Stevens, S.K., Liu, H., Jin, Z., Dyatkina, N. et al. Molecular Basis for the Selective Inhibition of Respiratory Syncytial Virus RNA Polymerase by 2’-Fluoro-4’-Chloromethyl-Cytidine Triphosphate. PLoS Pathog, 2015; 11: e1004995. 10.1371/journal.ppat.1004995

90. Salie, Z.L., Kirby, K.A., Michailidis, E., Marchand, B., Singh, K., Rohan, L.C., Kodama, E.N., Mitsuya, H., Parniak, M.A. and Sarafianos, S.G. Structural basis of HIV inhibition by translocation-defective RT inhibitor 4’-ethynyl-2-fluoro-2’-deoxyadenosine (EFdA). Proc Natl Acad Sci U S A, 2016; 113: 9274–9279. 10.1073/pnas.1605223113

91. Smith, D.B., Kalayanov, G., Sund, C., Winqvist, A., Maltseva, T., Leveque, V.J., Rajyaguru, S., Le Pogam, S., Najera, I., Benkestock, K. et al. The design, synthesis, and antiviral activity of monofluoro and difluoro analogues of 4’-azidocytidine against hepatitis C virus replication: the discovery of 4’-azido-2’-deoxy-2’-fluorocytidine and 4’-azido-2’-dideoxy-2’,2’-difluorocytidine. J Med Chem, 2009; 52: 2971–2978. 10.1021/jm801595c

92. Smith, D.B., Martin, J.A., Klumpp, K., Baker, S.J., Blomgren, P.A., Devos, R., Granycome, C., Hang, J., Hobbs, C.J., Jiang, W.R. et al. Design, synthesis, and antiviral properties of 4’-substituted ribonucleosides as inhibitors of hepatitis C virus replication: the discovery of R1479. Bioorg Med Chem Lett, 2007; 17: 2570–2576. 10.1016/j.bmcl.2007.02.004

93. Rondla, R., Coats, S.J., McBrayer, T.R., Grier, J., Johns, M., Tharnish, P.M., Whitaker, T., Zhou, L. and Schinazi, R.F. Anti-hepatitis C virus activity of novel beta-d-2’-C-methyl-4’-azido pyrimidine nucleoside phosphoramidate prodrugs. Antivir Chem Chemother, 2009; 20: 99–106. 10.3851/IMP1400

94. Klumpp, K., Kalayanov, G., Ma, H., Le Pogam, S., Leveque, V., Jiang, W.R., Inocencio, N., De Witte, A., Rajyaguru, S., Tai, E. et al. 2’-deoxy-4’-azido nucleoside analogs are highly potent inhibitors of hepatitis C virus replication despite the lack of 2’-alpha-hydroxyl groups. J Biol Chem, 2008; 283: 2167–2175. 10.1074/jbc.M708929200

95. Wang, G., Deval, J., Hong, J., Dyatkina, N., Prhavc, M., Taylor, J., Fung, A., Jin, Z., Stevens, S.K., Serebryany, V. et al. Discovery of 4’-chloromethyl-2’-deoxy-3’,5’-di-O-isobutyryl-2’-fluorocytidine (ALS-8176), a first-in-class RSV polymerase inhibitor for treatment of human respiratory syncytial virus infection. J Med Chem, 2015; 58: 1862–1878. 10.1021/jm5017279

96. Siegel, D., Hui, H.C., Doerffler, E., Clarke, M.O., Chun, K., Zhang, L., Neville, S., Carra, E., Lew, W., Ross, B. et al. Discovery and Synthesis of a Phosphoramidate Prodrug of a Pyrrolo[2,1-f][triazin-4-amino] Adenine C-Nucleoside (GS-5734) for the Treatment of Ebola and Emerging Viruses. J Med Chem, 2017; 60: 1648–1661. 10.1021/acs.jmedchem.6b01594

97. Michailidis, E., Marchand, B., Kodama, E.N., Singh, K., Matsuoka, M., Kirby, K.A., Ryan, E.M., Sawani, A.M., Nagy, E., Ashida, N. et al. Mechanism of inhibition of HIV-1 reverse transcriptase by 4’-Ethynyl-2-fluoro-2’-deoxyadenosine triphosphate, a translocation-defective reverse transcriptase inhibitor. J Biol Chem, 2009; 284: 35681–35691. 10.1074/jbc.M109.036616

98. Bonawitz, N.D., Clayton, D.A. and Shadel, G.S. Initiation and beyond: multiple functions of the human mitochondrial transcription machinery. Mol Cell, 2006; 24: 813–825. 10.1016/j.molcel.2006.11.024

